# Density-Preserving Data Visualization Unveils Dynamic Patterns of Single-Cell Transcriptomic Variability

**DOI:** 10.1101/2020.05.12.077776

**Authors:** Ashwin Narayan, Bonnie Berger, Hyunghoon Cho

## Abstract

Nonlinear data-visualization methods, such as t-SNE and UMAP, have become staple tools for summarizing the complex transcriptomic landscape of single cells in 2D or 3D. However, existing approaches neglect the local density of data points in the original space, often resulting in misleading visualizations where densely populated subpopulations of cells are given more visual space even if they account for only a small fraction of transcriptional diversity within the dataset. We present den-SNE and densMAP, our density-preserving visualization tools based on t-SNE and UMAP, respectively, and demonstrate their ability to facilitate more accurate visual interpretation of single-cell RNA-seq data. On recently published datasets, our methods newly reveal significant changes in transcriptomic variability within a range of biological processes, including cancer, immune cell specialization in human, and the developmental trajectory of *C. elegans*. Our methods are readily applicable to visualizing high-dimensional data in other scientific domains.

## Introduction

Exploratory analyses of large-scale biological datasets typically begin with visualizing the data in low dimensions, in the hopes of revealing high-level structural insights that can be probed in downstream analyses. This approach has been especially critical in the rapidly emerging field of single-cell transcriptomics, where high-throughput single-cell RNA sequencing (scRNA-seq) technologies are empowering researchers to study gene expression at an unprecedented resolution across diverse tissues, organisms, and biological conditions. Driven by the high-dimensionality of scRNA-seq datasets (thousands of different transcripts per cell) and their increasingly large-scale (hundreds of thousands of cells), many researchers rely on 2D or 3D data visualizations for quickly and intuitively finding structural patterns (e.g. clusters or trajectories) and communicating relevant biological findings with the scientific community^1, 2^.

Two of the most popular techniques for visualization of high-dimensional data are t-stochastic neighborhood embedding^3^ (t-SNE) and uniform manifold approximation and projection^4^ (UMAP), both of which have been widely adopted in scRNA-seq data analysis pipelines^5, 6^. In contrast to traditional methods for dimensionality reduction, e.g. principal component analysis (PCA), both t-SNE and UMAP learn a *nonlinear* embedding of the original space by directly optimizing the embedding coordinates of individual data points using iterative algorithms. Both methods aim to accurately preserve the original local neighborhood of each data point in the visualization, while being more permissive of distortions in long-range distances. Because of the expressiveness of nonlinear embeddings, t-SNE and UMAP are both well-regarded for their empirical performance in better elucidating sophisticated manifold structures and clustering patterns in high-dimensional data^1, 2^.

Despite their strengths, t-SNE and UMAP suffer from a major, often-overlooked pitfall: both methods neglect information about the local density of data points in the source dataset. In other words, data points whose neighbors are all very close-by in the original data are not distinguished in the visualization from those whose neighbors are far away. This limitation leads to misleading visualizations where the apparent size of a cluster largely reflects the number of points in the cluster rather than its underlying heterogeneity, as we demonstrate in our results. In scRNA-seq data, this omitted information about heterogeneity corresponds to the *variability* of gene expression within a subpopulation of cells. By inaccurately portraying differences in local density, these algorithms produce potentially misleading visualizations of the transcriptomic landscape of single cells. Capturing differences in variability within a dataset could provide another “dimension” of information in the visualization, reflecting heretofore hidden biological insights.

Here, we introduce *density-preserving* data visualization methods den-SNE and densMAP that build upon t-SNE and UMAP, respectively, to enable researchers to more accurately visualize and communicate deeper biological insights from the growing compendium of single-cell transcriptomic experiments. Our methods leverage the insight that, since both t-SNE and UMAP construct their embeddings by iteratively optimizing an objective function, we can augment that objective function with an auxiliary term that measures the distortion of local density at each data point in the visualization. To this end, we develop a general, differentiable measure of local density, called the *local radius*, which intuitively represents the average distance to the nearest neighbors of a given point. Our design of this measure enables efficient optimization of the density-augmented visualization objective. The algorithmic techniques we introduce for transforming t-SNE and UMAP into their density-preserving counterparts could be used to enhance other visualization tools based on iterative optimization and thus are of general interest.

To demonstrate the utility of density-preserving visualization, we applied den-SNE and densMAP to a range of published scRNA-seq datasets from lung cancer patients^7^, human peripheral blood cells^8^ and embryonic roundworm *Caenorhabditis elegans*^9^, as well as the UK Biobank human genotype profiles and the canonical MNIST hand-written digit images. Our density-preserving methods not only capture additional information beyond existing visualization techniques but also biological insights others miss, including immune cell transcriptomic variability in tumors; specialization of monocytes and dendritic cells; and temporally modulated transcriptomic variability across developmental lineages of *C. elegans*. We validated our key findings from the lung cancer and peripheral blood data on additional datasets. Our work shows that density-preserving data visualization can efficiently unveil unforeseen patterns in single-cell transcriptomic landscapes and enrich our understanding of biology.

## Results

### Overview of density-preserving data visualization

Our density-preserving visualization methods, den-SNE and densMAP, augment t-SNE and UMAP respectively, generating embeddings that preserve both local structure and variability in the original data (Figure 1 and Methods). In order to capture the local structure of the data, t-SNE and UMAP both create a nearest-neighbors graph and preserve only the distances between points that are neighbors in this graph. We use the same nearest-neighbors graphs that underpin each of the original methods to calculate a *local radius* around each point, which represents the average distance from the point to its nearest neighbors; this conveys the density of that point’s neighborhood. The two original algorithms both have an objective function that quantifies the agreement between a given embedding and the original nearest-neighbors graph, and they rearrange the embedding to maximize this agreement. We thus augment these objective functions to enable density preservation by incorporating a term that measures the agreement between local radii in the original dataset and in the embedding. The resulting embedding thus jointly optimizes the original objective and our density-preservation objective, ensuring local structure is still preserved while also conveying information about variability. Our techniques have strong theoretical foundations, enable efficient optimization, and can be easily generalized to other data visualization algorithms that similarly use gradient-based optimization, allowing density-preservation to be broadly achievable (Methods, Supplementary Notes 1–3).

**Figure 1:**
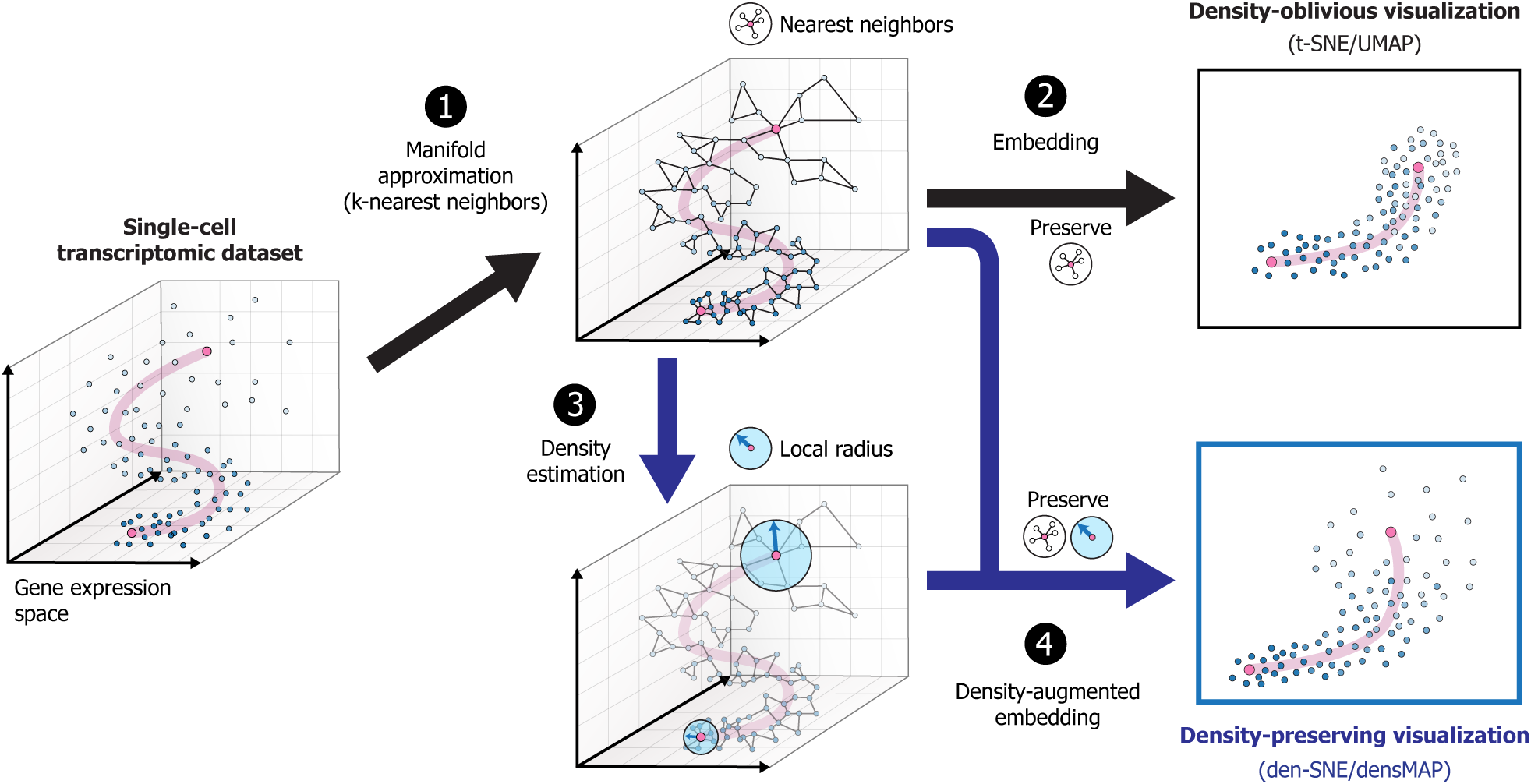
Overview of density-preserving data visualization. Given a set of points in a high-dimensional space as input (e.g. gene expression profiles from single-cell RNA-seq experiments), the goal of data visualization is to embed these points in 2D or 3D while preserving the structure of the original data. To this end, standard visualization tools t-SNE and UMAP first construct the *k*-nearest neighbor (KNN) graph as a compact summary of the data manifold (**1**). These methods then optimize the visualization coordinates of the points to maximally preserve local distances between neighbors in the graph (**2**). However, because t-SNE and UMAP adaptively choose length-scale to normalize local distances within each neighborhood, they produce visualizations that neglect information about density in the original space, thus omitting a key structural feature of the data. To enhance data visualization by incorporating density information, we introduce a general, differentiable measure of density called the local radius (Methods), which is efficiently calculated on the same KNN graphs that t-SNE and UMAP leverage (**3**). By augmenting the original visualization objective with a new term that encourages local radii to be consistent between the original space and the visualization, we transform both t-SNE and UMAP into density-preserving counterparts, den-SNE and densMAP, which more accurately portray the structure of the underlying data (**4**).

Applying our methods to visualize simulated datasets featuring heterogeneous density landscapes revealed the misleading visual conclusions that could be made without density preservation (Figure 2). In visualizing a mixture-of-Gaussian point clouds with different variances, t-SNE and UMAP generate clusters that are all of the same size, while den-SNE and densMAP accurately depict the different variances (Figure 2a). When the point clouds are translated linearly with overlap, reflecting a trajectory, the lack of density preservation in t-SNE and UMAP obscures the dynamic changes in variability over the trajectory (Figure 2c). Conversely, when size is constant but a region is oversampled, t-SNE and UMAP overrepresent this oversampled region, giving the impression of increased variability and downplaying the undersampled regions (Figures 2b and d). Our results that follow show that these considerations are critical in biological analyses.

**Figure 2:**
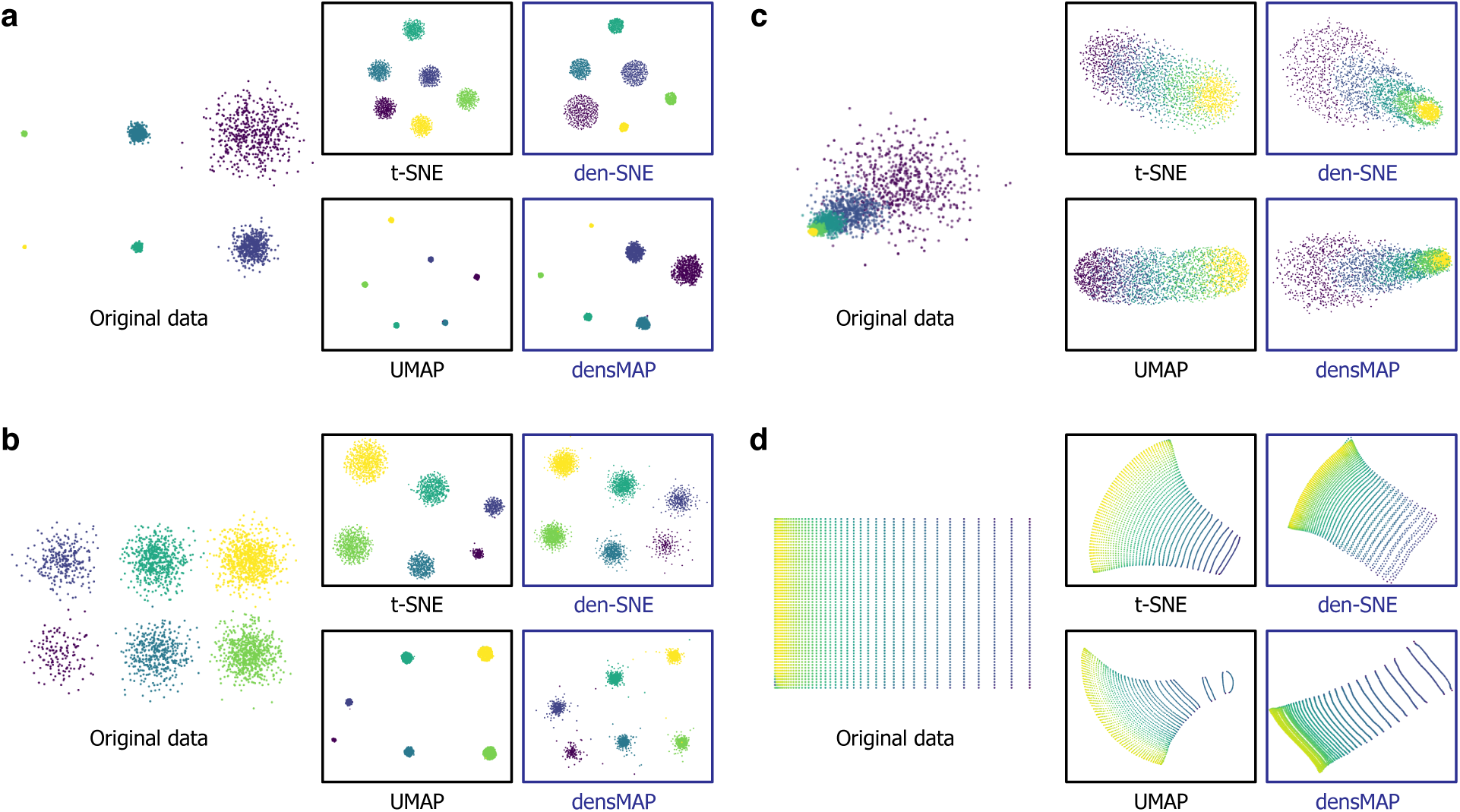
Density-preserving visualization more accurately captures the true underlying shape of synthetic datasets than existing tools. We compared the visualizations of our density-preserving methods den-SNE and densMAP to those of t-SNE and UMAP on different synthetic datasets: mixture-of-Gaussian point clouds with (**a**) increasing variances with the same sampling rate; (**b**) same variance, but with increasing sampling rates; (**c**) increasing variances in a linear translational motion with overlap, representing a temporal trajectory; and (**d**) a grid of points, whereby the density grows linearly in one direction. The synthetic datasets are generated in twenty dimensions for the point clouds and two dimensions for the grid, and the depictions of the original data in the figure represent two-dimensional linear projections for the former. While t-SNE and UMAP produce misleading visualizations where the apparent size of a cluster of points (marked by different colors) is unrelated to the amount of space it occupies in the original space and is biased by sampling rate, den-SNE and densMAP more accurately portray the shape of the original data by preserving density information.

### Visualizing the heterogeneity of immune cells in tumor

To illustrate the value of density-preserving visualization for biological studies, we first applied our methods to a published scRNA-seq dataset of 41,861 immune cells in matched tumor and peripheral blood samples from seven non-small-cell lung cancer (NSCLC) patients^7^. The original study identified distinct transcriptomic states spanned by tumor-infiltrating myeloid cells that were reproducibly observed across different individuals, suggesting their potential relevance for cancer immunotherapies. We asked whether our visualization tools could capture the gene expression landscape of tumor-infiltrating immune cells more accurately than existing methods and help uncover new biological insights.

Comparison of den-SNE and t-SNE embeddings revealed several immune cell types with noticeable differences between the visualizations (Figure 3). For example, tumor-infiltrating neutrophils and plasma cells occupy considerably more space in the den-SNE visualization than their t-SNE counterparts, while tumor-infiltrating T cells are relatively smaller in den-SNE. These discrepancies arise because visual size of a cluster in t-SNE corresponds more closely to the number of cells in the cluster than to underlying variability. Thus, in t-SNE, tumor-infiltrating neutrophils (*n* = 2861) occupy much less space than circulating neutrophils (*n* = 9217) despite den-SNE indicating they have comparable variability. The rich transcriptomic diversity of tumor-infiltrating plasma cells is also lost in t-SNE. Conversely, T cells, the most populous cell type in tumors (*n* = 10701) are visually overrepresented in t-SNE relative to their actual variability.

**Figure 3:**
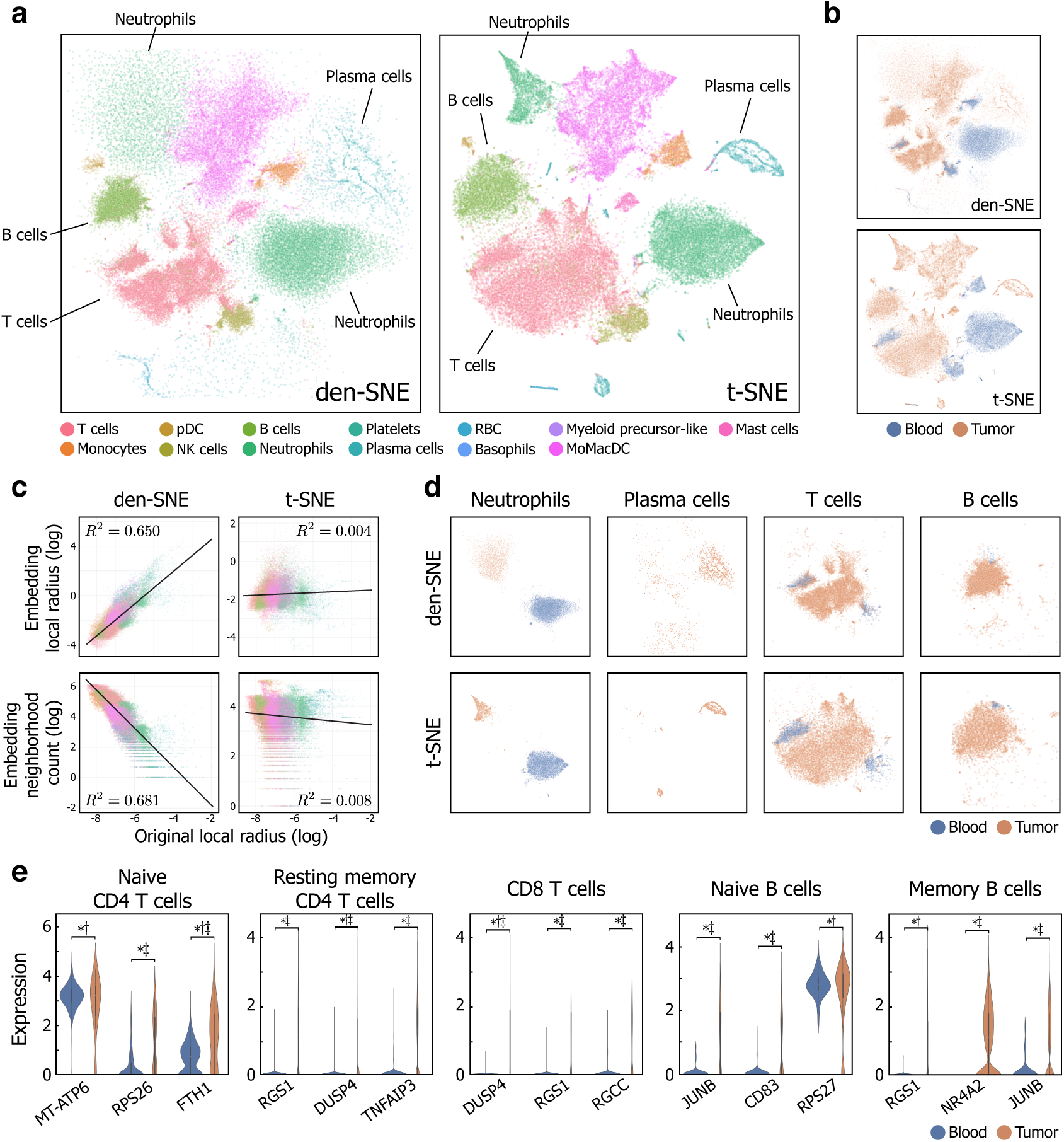
Density-preserving visualization reveals heterogeneity in transcriptomic variability of immune cells in blood and tumor. We visualized a dataset of tumor and blood immune cells from lung cancer patients^7^ using den-SNE and t-SNE, colored by (**a**) cell type and (**b**) tissue type (tumor or blood); den-SNE exposes striking density differences between immune cell types and between blood and tumor, which cannot be analyzed based on t-SNE due to its theoretical lack of density-preservation (Methods). Note that the relative heterogeneity of neutrophils, plasma cells, and T cells are misleadingly portrayed in the t-SNE visualization. **c.** Scatter plots comparing the local radii, our measure of local density (Methods), in the original space and two measures of visual density (local radius and neighborhood count; see Methods) in the visualization (embedding) for den-SNE and t-SNE. Points are colored by cell type, and the *R*^2^ value of the correlation is shown for each plot. Higher correlations of den-SNE (inverse correlation for neighborhood count) show that den-SNE more accurately conveys the density landscape of the original data than t-SNE. The radius for neighborhood count is set to two times the average length-scale of each visualization (Methods); other choices of length-scale show a similar improvement for den-SNE (Supplementary Figure 2). **d.** For detailed comparison, we plot the same visualizations for den-SNE (top) and t-SNE (bottom), restricted to each of four notable cell types (neutrophils, plasma cells, T cells, and B cells) and colored by tissue type (tumor or blood). Neutrophils and plasma cells in tumor considerably expand in size in den-SNE, reflecting transcriptomic variability previously hidden in t-SNE. T and B cells show a large increase in heterogeneity in tumor compared to blood in den-SNE. Although t-SNE shows a similar pattern, its lack of a density-preservation property precludes reasoning about differences in heterogeneity. **e.** Violin plots showing the distributions of gene expression in tumor and blood for the top three genes with the highest increase in variance in tumor for each subtype of T and B cells. A more comprehensive list of genes for each cell type is included in Supplementary Tables 1–5. These genes indicate potential biological mechanisms underlying the increased heterogeneity (revealed by den-SNE) of T and B cells in tumor. The markers ∗, †, and ‡ denote a statistically-significant difference in variance, dispersion, and mean, respectively, between blood and tumor (Bonferroni-corrected *p <* 0.01; Methods). All genes shown have significant variance difference, and several of them are not accompanied by a shift in mean expression (e.g. RPS27 in naive B cells), suggesting new biological insights about tumor not captured by conventional differential expression analysis. We provide the same plots for densMAP and UMAP in Supplementary Figure 1.

To quantitatively evaluate the improvement in density preservation that our algorithms offer, we calculate two complementary measures of local density in the visualization—(i) local radius and (ii) neighborhood count (see Methods)—and assess their correlation with the local radii computed in the original data space, which represent the underlying variability in the dataset. Both measures aim to quantify our perception of density in the visualizations (inversely-related for local radius); intuitively, the local radius captures the size of a neighborhood that contains a fixed number of nearest neighbors, and the neighborhood count captures the number of points within a fixed radius around each point. The former is consistent with how our algorithms model visual density for efficient optimization, while the latter is arguably a more direct notion of density previously used in the literature on visual perception^10^.

The accuracy of den-SNE’s visualization of local density is confirmed by the high correlation based on both measures (*R*^2^ = 0.650 for local radius; average *R*^2^ = 0.657 for neighborhood count across different length-scales) compared to t-SNE (*R*^2^ = 0.004; *R*^2^ = 0.023) (Figure 3c; Supplementary Figure 2). Results with densMAP (*R*^2^ = 0.590 for local radius; average *R*^2^ = 0.632 for neighborhood count) and UMAP (*R*^2^ = 0.045; *R*^2^ = 0.008) are analogous (Supplementary Figures 1 and 2).

Our visualizations motivate *transcriptomic variability* as a key distinguishing factor among cell types and biological conditions. To illustrate, we examined tumor-infiltrating lymphocytes (TILs; white blood cells, including T and B cells, that migrate to tumor tissue) compared to those in blood. While essential in the anti-tumor immune response^11^, these cells’ molecular mechanisms in cancer remain poorly understood. A key aspect of TIL biology newly highlighted by density-preserving visualization is the increased transcriptomic heterogeneity of T and B cells compared to their counterparts in blood (Figure 3d). Despite an apparent size-difference between the tumor and blood TILs in t-SNE, lack of density-preservation means this pattern could only imply a difference in cell counts, not in heterogeneity of expression.

Ranking genes by their contribution to the increase in transcriptomic heterogeneity in tumor implicated several biological processes as potential driving factors of TIL diversity (Methods; Supplementary Tables 1–5). Top genes for CD8 T cells and CD4 memory T cells were significantly enriched in negative regulation of IL2 production, transcription, and metabolic processes, suggesting that T cells in tumor are subjected to variable degrees of proliferation control, likely in response to complex biochemical signals in the tumor microenvironments (Supplementary Tables 7 and 8). Notably, RGS1 and DUSP4 showed the largest difference in variability for both T cell types. RGS1 encodes a regulator for the G-protein signaling pathway known to be involved in chemokine-induced lymphocyte migration^12^, and DUSP4 encodes a phosphatase that modulates a T cell receptor signaling pathway with known association with immunological disorders^13^. We validated the variability difference of these two genes in CD8 T cells between tumor and blood based on another scRNA-seq dataset of TILs from NSCLC patients^14^, along with 7 other genes in our list of genes ranked by contribution to variability (9 out of 19 genes were found to have significant increase in variance in tumor in the validation dataset; Supplementary Table 6). On the other hand, top genes for naïve CD4 T cells are enriched in proteins targeting membranes and in those that ensure the decay of mis-transcribed mRNA (Supplementary Table 9). For B cells, key biological processes underlying the variability difference included leukocyte activation and protein complex assembly for memory B cells, and response to cyclic AMP (a known modulator of cell proliferation) and biotic stimulus for naïve B cells, along with transcriptional and metabolic regulation processes similar to those implicated for T cells (Supplementary Tables 10 and 11).

While many genes implicated here are lowly-expressed in blood and activated in tumor, we also found a substantial portion (42% among top 20 genes across all cell types) that show statistically significant *overdispersion* in tumor, whereby the increase in variance cannot be explained by an increase in mean expression (Methods, Supplementary Tables 1–5). In fact, some genes, e.g. RPS27 in naïve B cells, which encodes the MPS-1 protein that modulates the activity of tumor-suppressor p53^15^, show a significant increase in variance *without* a significant change in mean. These genes are especially common in the top genes for naïve CD4 T cells. Their lack of difference in mean expression implies that these key distinguishing genes cannot be identified by conventional differential expression analysis. Moreover, since standard visualization algorithms separate clusters largely based on difference in mean expression, the effects of these genes are lost in their visualizations. Our findings demonstrate that the transcriptomic variability landscape uncovered by our visualizations helps open new analytic directions for the study of anti-tumor immune response.

### Visualizing immune cell specialization and diversification in peripheral blood

While the above illustrates changing patterns of variability that come about due to disease, we show here that variability of expression *within* cellular subtypes itself reveals interesting underlying biology. We used densMAP to visualize a benchmark scRNA-seq experiment that profiled 68,551 peripheral blood mononuclear cells (PBMC) from 10X Genomics^8^. While both UMAP and densMAP separate the various clusters corresponding to different cell types, the densMAP embedding considerably expands the sizes of natural killer (NK) cells, cytotoxic T cells, CD14+ monocytes and dendritic cells (DCs), and shrinks the size of naïve cytotoxic T cells (Figure 4a). Similar to the cancer dataset, the sizes of these clusters in UMAP correspond to the number of cells belonging to them and thus do not accurately reflect their variability of expression. By quantifying the agreement between the local radius in the original dataset and the local density measures in each of the visualizations (Methods), we confirmed that densMAP more accurately preserves density (*R*^2^ = 0.712 for local radius; average *R*^2^ = 0.727 for neighborhood count), compared to UMAP (*R*^2^ = 0.000; *R*^2^ = 0.000) (Figure 4c; Supplementary Figure 4). The same pattern is observed when comparing den-SNE to t-SNE, with the density correlations in den-SNE much higher (*R*^2^ = 0.704 for local radius; average *R*^2^ = 0.696 for neighborhood count) than in t-SNE (*R*^2^ = 0.052; *R*^2^ = 0.037) (Supplementary Figures 3 and 4).

**Figure 4:**
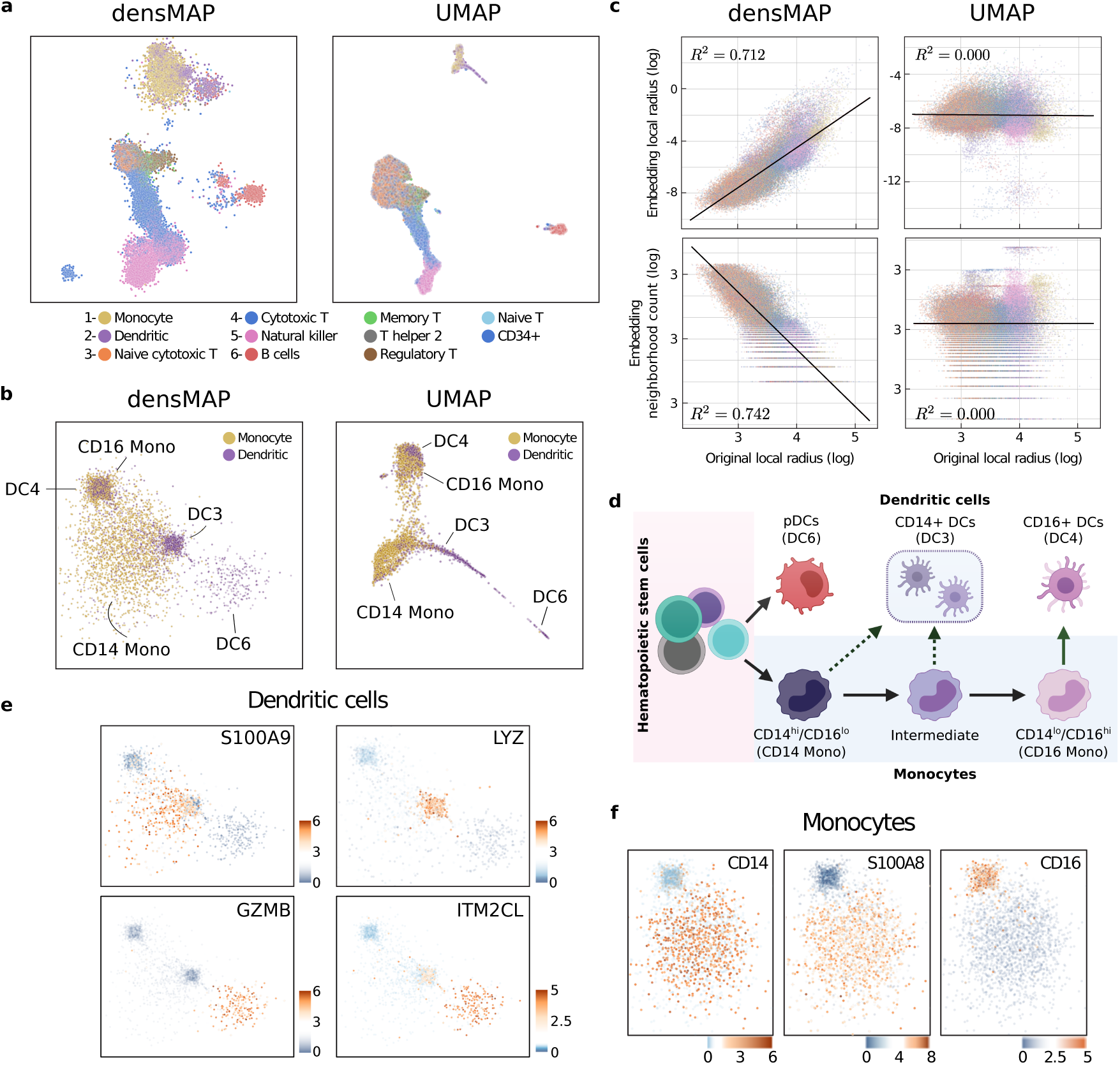
Density-preserving visualization of peripheral blood mononuclear cells reveals monocyte and dendritic cell subsets that differ in transcriptomic variability. **a.** We visualized the PBMC dataset^8^ using densMAP (left) and UMAP (right), colored by cell type. The group of clusters corresponding to monocytes (cluster 1) and dendritic cells (DCs; cluster 2) showed the most pronounced difference between the two visualizations. **b.** For a detailed comparison, we plotted the same visualizations restricted to the monocyte-DC subset, which revealed distinct subtypes of monocytes (CD16 Mono and CD14 Mono) and DCs (DC3, DC4, and DC6) with clear density differences in densMAP. Each subtype is annotated using the classification from the PBMC2 study^19^ based on marker gene expression. Although the same subtypes are visible in UMAP, their relative density differences are lost. **c.** Scatter plots comparing the local radii, our measure of local density (Methods), in the original space and two measures of visual density (local radius and neighborhood count; see Methods) in the visualization (embedding) for densMAP and UMAP. Points are colored by cell type, and the *R*^2^ value of the correlation is shown for each plot. Higher correlations in densMAP (inverse correlation for neighborhood count) support the validity of the observed density differences between the monocyte and DC subtypes in the densMAP visualization. The radius for neighborhood count is set to the average length-scale of each visualization (Methods); other choices of length-scale show a similar improvement for densMAP (Supplementary Figure 4). **d.** Graphical illustration showing the biological relationships among the five monocyte and DC subtypes we found in the monocyte-DC subset. Under inflammatory conditions, CD14 Mono (classical monocytes) differentiate into CD16 Mono (non-classical monocytes) for immune response. Both CD14 Mono and CD16 Mono can differentiate into DCs (classified as DC3 and DC4, respectively). DC6 represents plasmacytoid DCs (pDCs), which come from a different differentiation trajectory than the rest. densMAP visualization suggests that the differentiation paths from CD14 Mono to CD16 Mono and DC3 both represent specialization with considerable decrease in transcriptomic variability. densMAP also reveals rich heterogeneity of DC6 previously hidden in UMAP. **e.** Gene expression heatmaps of DC marker genes from the PBMC2 study^19^ for DC3 (top) and DC6 (bottom) in the densMAP visualization restricted to DCs. These support our assignment of DC clusters to DC3 and DC6. A comprehensive set of heatmaps as well as violin plots of all marker genes for DC3, DC4, and DC6 are provided in Supplementary Figure 5. **f.** Gene expression heatmaps of monocyte marker genes CD14, S100A8, and CD16 in the densMAP visualization restricted to monocytes. The patterns of expression support our classification of the dense cluster as CD16 Mono and the sparse cluster as CD14 Mono. We provide the same plots for den-SNE and t-SNE in Supplementary Figure 3. Validation of our newly observed variability differences among monocyte and DC subtypes on two additional datasets^19, 20^ is included in Supplementary Figure 6.

We focus here on the visualization of the monocyte and DC clusters, which are strikingly different between the two visualizations (Figure 4b). While both reveal two subtypes of monocytes, the densMAP visualization separates them by density, with a dense subcluster adjacent to a much sparser one. These monocytes begin life as *classical* monocytes, which are characterized by expression of CD14 and a lack of CD16 (also called FCGR3A); these can then differentiate into CD16 monocytes, macrophages, or dendritic cells (DCs)^16^ (Figure 4d). Overlaying the expression of marker genes reveals that the sparse cluster is populated with classical monocytes and the dense cluster, with CD16 monocytes (Figure 4f). This trajectory has intriguing biological significance. Recent work has upended the neat categorization of monocytes into classical (CD14) and non-classical (CD16), and revealed that monocytes are an extremely heterogeneous cell type with complex intermediate states^17^ and high transcriptional diversity^18^. However, non-classical monocytes are more specialized: they are thought to emerge from a small population of intermediate (CD14+CD16+) monocytes and spike very rapidly during infections^17^; since their progenitor cell is rare, this supports the notion of a bottleneck in transcriptional diversity for non-classical monocytes.

We validated this difference in variability between classical and non-classical monocytes in two other scRNA-seq datasets of immune cells, one that profiled 1,078 monocytes, DCs and their subtypes^19^ (PBMC2) and the other that profiled 13k PBMCs from two healthy donors^20^ (PBMC3). In both, the classical monocytes were indeed sparser than the non-classical ones (Mann-Whitney U test, *p* = 6.61 *×* 10^−7^ for the PBMC2 dataset, *p* = 2.89 *×* 10^−4^ for the PBMC3 dataset, see Methods and Supplementary Figure 6). Unlike UMAP, densMAP clearly depicts this dichotomy of variability in expression.

A similar analysis can be performed on the DC subset: this cell type shows (i) a dense cluster of cells adjacent to the CD14 monocytes, (ii) a dense cluster overlapping the CD16 monocytes, and (iii) a sparser cluster near the CD14 monocytes (Figure 4b). While the classification of dendritic cells is still an area of active research^21^, the colocalization of the DCs (i) and (ii) and the monocytes in the densMAP visualization indicates that these DCs originate from monocytes. By analyzing the expression of the marker genes of DC subtypes identified by the the PBMC2 study^19^ in these DC subsets, we hypothesize that (i) corresponds to classical monocyte-derived DCs (cDCs), with marker genes S100A9, S100A8, VCAN, LYZ, and ANXA1 (DC3 in PBMC2); (ii) corresponds to the poorly understood CD141– CD1C– DCs, with marker genes CD16, FTL, SERPINA1, LST1, and AIF1 (DC4 in PBMC2); and (iii) corresponds to plasmacytoid DCs (pDCs), with marker genes GZMB, IGJ, SERPINF1, ITM2C (DC6 in PBMC2) (see Supplementary Figure 5).

**Figure 5:**
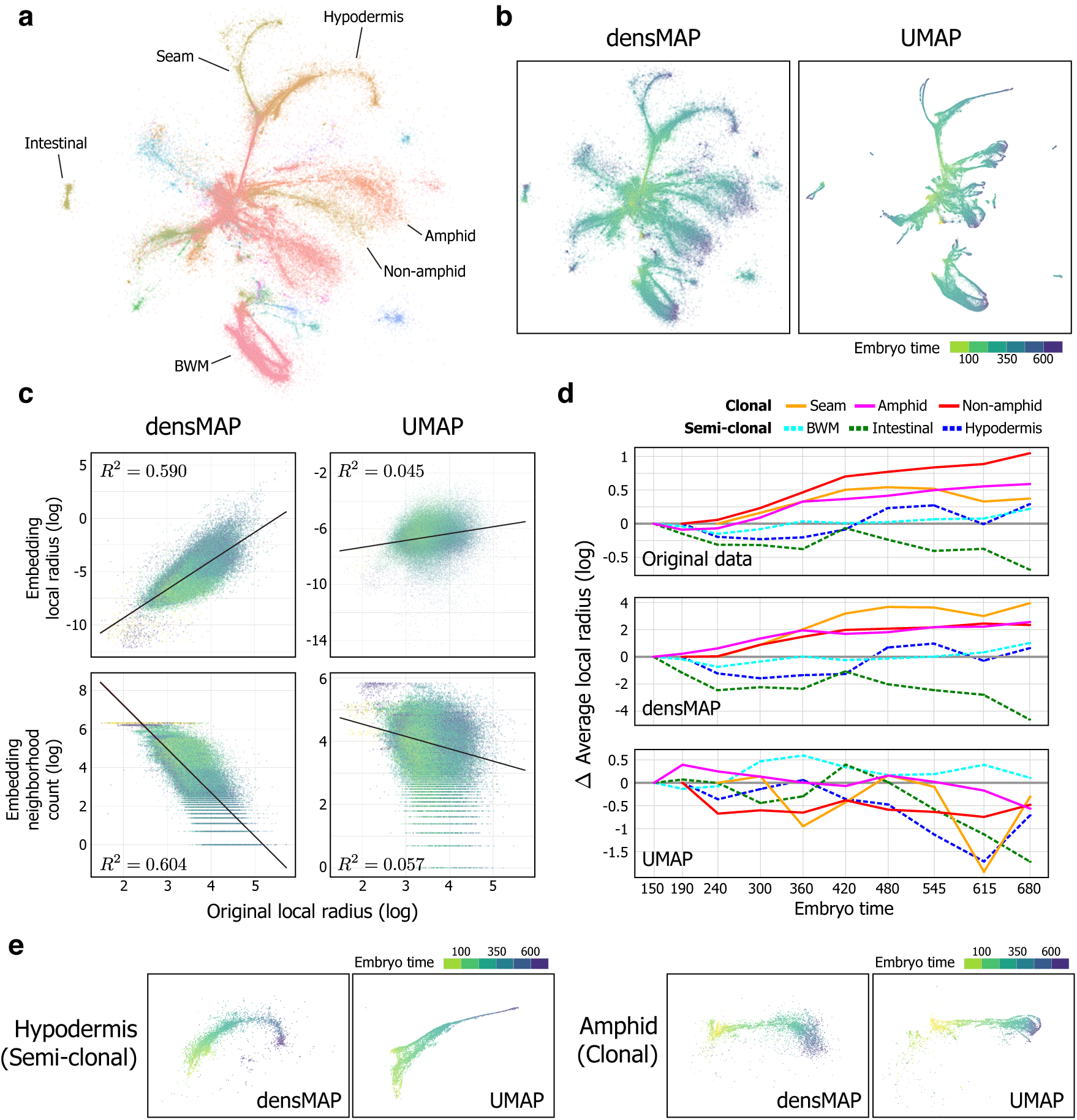
Density-preserving visualization of *C. elegans* development reveals temporal dynamics of transcriptomic variability in different developmental lineages. We visualized the *C. elegans* dataset^9^ using densMAP and UMAP, colored by (**a**) cell type (major cell types labeled) and (**b**) embryo time. UMAP visualization with cell type coloring is omitted for space. In contrast to UMAP, densMAP clearly conveys an overall increase in transcriptomic variability as the organism develops and realizes a wider range of biological functions. **c.** Scatter plots comparing the local radii, our measure of local density (Methods), in the original space and two measures of visual density (local radius and neighborhood count; see Methods) in the visualization (embedding) for densMAP and UMAP. Points are colored by embryo time, and the *R*^2^ value of the correlation is shown for each plot. Higher correlations of densMAP (inverse correlation for neighborhood count) support the validity of the overall increase in transcriptomic variability over the course of development observed in densMAP. The radius for neighborhood count is set to the average length-scale of each visualization (Methods); other choices of length-scale show a similar improvement for densMAP (Supplementary Figure 8). **d.** To assess lineage-specific patterns of transcriptomic variability, we summarized the average local radius of each cell type (marked by different line style) within each embryo time interval for the original data (top), densMAP (middle), and UMAP (bottom). The plot for original data represents the temporal changes in the underlying transcriptomic variability of each cell type, and the plots for densMAP and UMAP represent apparent changes in variability based on the respective visualizations. We used the time intervals provided by the original study, and the y-axis shows the change in average local radius compared to the earliest time interval in log scale. densMAP closely follows the temporal patterns of each cell type in the original dataset, a structural insight that is lost in UMAP. These patterns uniquely captured by densMAP highlight the relatively constant variability of semi-clonal lineages (BWM, intestinal, and hypodermis) in contrast to the increasing variability of clonal lineages (seam, amphid and non-amphid neurons), which can be explained by the more intermixed nature of semi-clonal development. **e.** densMAP and UMAP visualizations restricted to hypodermis and amphid cells for comparison, colored by embryo time. densMAP captures constant variability of hypodermis cells during development, whereas UMAP vastly under-represents the variability of the terminal cell state. Similarly, for amphid cells, densMAP accurately portrays expanding variability, a pattern that is lost in UMAP. We provide the visualizations of other cell types and repeat the analyses for den-SNE and t-SNE in Supplementary Figure 7. BWM: body-wall muscle.

Our visualizations of the DCs reveal striking differences in variability compared to the standard visualization algorithms. The DC3 cluster is far denser than the CD14 monocytes colocated with it, hinting that, as with CD16 monocytes, these cells specialize as they develop from CD14 monocytes. This pattern also holds in PBMC2, where the DC3 cluster is significantly denser than the classical monocyte cluster (Mann-Whitney U test, *p* = 5.43 *×* 10^−14^; Methods and Supplementary Figure 6). The PBMC3 dataset was omitted from this analysis as it contained too few DCs to draw conclusions about subtypes. In addition, the pDC cluster expands drastically in the density-preserving visualization compared to the standard visualization, revealing previously hidden variability (Figure 4b).

We also note the DCs dispersed throughout the CD14 monocytes (Figure 4b). When we classify the DC3 subset into dense and sparse categories based on their original local radius (with a threshold of 3.9 for the log local radius), we find that the sparse subset has *intermediate* expression of the genes that mark DC3 and those that mark CD14 monocytes (Supplementary Figure 6). While this could be due to misclassification (the original study assigned cell types based on similarity to purified samples), it could also indicate a bridging state between the two cell types. These results suggest that there are key differences in transcriptomic variability among immune cell subtypes that are hidden by existing visualization tools.

### Visualizing time-varying transcriptomic heterogeneity in *C. elegans* development

To explore embryo development at high-resolution, Packer et al. (2019) performed scRNA-seq profiling of *C. elegans* to create an atlas of gene expression at almost every cell division of the embryo^9^. Given that visualizing these data illustrates trajectories of development and differentiation, we asked whether density-preserving visualization could better capture the diversification (or lack thereof) of different developmental lineages, complementing investigations into time-dependent patterns of gene expression in organism development^22–24^.

For most of the cell types profiled, the lineage distance between cells correlates strongly with transcriptome dissimilarity, and many cells from the same progenitor diverge after gastrulation^9^. Thus, an accurate visualization should show that the density of cells for most cell types decreases over time (reflecting increasing diversity), as the cells adopt their terminal fates. While both densMAP and UMAP show a central “progenitor” region that branches into the different major tissues, densMAP more clearly highlights the increase in variability in the outer branches of the lineages (Figures 5a and b). Evaluating the agreement between the local radius in the original dataset and both measures of local density in the visualization show that densMAP (*R*^2^ = 0.590 for local radius; average *R*^2^ = 0.585 for neighborhood count) more accurately preserves density than UMAP (*R*^2^ = 0.045; *R*^2^ = 0.052) (Figure 5c and Supplementary Figure 8). Results are analogous when comparing den-SNE (*R*^2^ = 0.619 for local radius; average *R*^2^ = 0.596 for neighborhood count) to t-SNE (*R*^2^ = 0.000; *R*^2^ = 0.063) (Supplementary Figures 7 and 8).

While transcriptomic variability generally increases over the course of differentiation, notable exceptions are also made apparent by densMAP. Specifically, of the cell types well-represented (greater than 1000 cells), the intestinal, body-wall muscle (BWM), and hypodermis cells show a relative homogeneneity in density (as measured by the average local radius in the original dataset across time) throughout embryo development when compared to the other cell types, e.g. both non-amphid and amphid neurons and seam cells; densMAP more accurately preserves these temporal density trajectories than UMAP (Figures 5d and e).

These visual patterns are supported by the underlying biology, since intestinal, BWM, and hypodermis cells are so-called *semi-clonal lineage clades*^9^. A semi-clonal lineage model is intermediate between *clonal* development, which closely adheres to the lineage structure whereby branching of cell fates leads to increasingly more divergent cells, and *non-clonal* development, where the daughter cells are only loosely associated with their progenitors and different lineage branches share more commonalities through horizontal transitions^25^. As semi-clonal cell types, the intestinal, BWM, and hypodermis cells are thus expected to remain more transcriptionally coherent across time (and therefore more compact in expression space) than clonal lineages. When we compare the average change in density over embryo time for semi-clonal cells, this change is considerably lower than the average change for the other cell types (Supplementary Figure 7). The difference in density between these semi-clonal cell types and the rest is made clear in our density-preserving visualization but is completely hidden in a standard UMAP. In fact, the UMAP plots tend to show a *decrease* in density in many cell lines because fewer cells were profiled at the late time-points (Supplementary Figure 7). These results show that our methods can accurately portray continuous changes in transcriptomic variability in developmental trajectories, which are not captured by existing visualization tools.

### General applicability of density-preserving data visualization

Visualizing high-dimensional data is broadly useful, with applications both within and outside biology. Like t-SNE and UMAP, our density-preserving methods require only a distance metric defined between data points. To illustrate the performance of our methods on other data domains, we analyzed a genotype dataset from the UK Biobank and the MNIST image dataset widely used by the machine learning community (Methods).

The UK Biobank^26^ (UKBB) is an ambitious project collecting extensive genotypic and phenotypic data from British individuals for use in health-related research. Due to the skew in ethnicity of the British population, most of the individuals in the dataset self-identify as white (94% of the 534k individuals). This lack of diversity has raised important concerns about ethnic biases in downstream scientific analyses^27^. When visualizing the individuals in the dataset based on their genotype profiles, an analytic approach that is increasingly being explored^28^, t-SNE and UMAP show the cluster corresponding to whites disproportionately large, while the clusters corresponding to Asian and black people can scarcely be seen (Supplementary Figure 9). Visualizing this data using den-SNE and densMAP results in a more balanced representation of the ethnicities, with considerable expansion of people-of-color clusters and shrinking of the white cluster (Supplementary Figure 9). Thus, it is clear that the existing visualization tools grossly under-represent the genetic diversity of minority populations due to their limited sample sizes. Even among the white population, the density-preserving visualizations obtain a more balanced representation of subpopulations (computationally identified; Methods). In the UMAP and t-SNE visualizations, only the two most populous subgroups take up significant space, whereas densMAP and den-SNE show five subgroups with comparable diversity.

A complementary situation occurs in the MNIST dataset, a dataset of handwritten digit images (Methods). Here, t-SNE and UMAP generate ten evenly sized clusters; den-SNE and densMAP visualizations, however, reveal that the cluster corresponding to the digit **1** is strikingly less variable than the other digits (Supplementary Figure 11). This is as expected, since **1** is drawn with considerably limited degrees of freedom. Analyzing the local radii in the original data reveals that, indeed, **1** has a much higher density than the other digits (Supplementary Figure 11). The improved accuracy of our visualizations for UKBB and MNIST datasets are supported by both density-preservation metrics based on local radius and neighborhood count (Supplementary Figures 9 to 12). Taken together, these results show that density-preserving visualization is able to reveal important insights about the data not captured by the existing methods on diverse types of datasets.

### Density-preserving visualization is almost as efficient as existing approaches

As experimental methods continue to generate larger and larger datasets, computational tools to analyze them need to scale as well. By leveraging computations already done by t-SNE and UMAP, our density-preserving methods have the same asymptotic scaling as those methods. For both den-SNE and densMAP, the additional computations are *O*(*n*) in dataset size. Given the *O*(*n* log *n*) complexity of t-SNE (with the standard Barnes-Hut approximations^29^), the ratio of den-SNE computation time to t-SNE computation time decreases as dataset size grows (Supplementary Figure 13). densMAP largely retains the computational efficiency of UMAP by similarly leveraging stochastic gradient descent (Methods), and the relative overhead of densMAP remains constant as the dataset becomes larger (Supplementary Figure 13). Although density preservation increases the overall runtime of den-SNE and densMAP (about 20% overhead for both methods on 86k cells; see Supplementary Figure 13), we believe that this additional cost is not onerous, when weighed against additional information conveyed by accurately depicting density. The memory requirement of den-SNE is virtually identical to that of t-SNE and densMAP uses about 150MB more peak memory across the range of data sizes (20k to 86k) we profiled (Supplementary Figure 13).

## Discussion

Effective tools for visualizing the single-cell landscapes captured by ever-larger single-cell experiments are pivotal for accelerating discoveries and disseminating them among researchers. In this work, we presented novel density-preserving visualization tools den-SNE and densMAP. These overcome a major limitation of the state-of-the-art tools t-SNE and UMAP: they neglect differences in the *local variability* of gene expression across the transcriptomic landscape spanned by the cells in the dataset, resulting in misleading distortions in the visualizations. While t-SNE and UMAP are still useful for revealing clustering or trajectory patterns, we demonstrated on a range of published scRNA-seq datasets that the new “dimension” of information, i.e. local density, that we incorporate into our visualizations harbors biological insights that can help enrich our understanding of the underlying biological processes beyond what can be achieved with existing visualization tools. Our techniques for augmenting density information are also broadly applicable to other visualization algorithms, including recent extensions of t-SNE^30, 31^ and force-directed layout embedding^32, 33^ (FDLE).

Theoretically, targeted analyses could capture directly the changes in transcriptomic variability made apparent by our visualizations (e.g. by directly comparing variability between cell types^34^). However, by visualizing local variability over the entire dataset, our approach allows easier interpretation and understanding. This methodological shift is akin to how t-SNE and UMAP have significantly streamlined cell type identification workflows by visually revealing clustering patterns in the dataset, despite the fact that one could potentially apply and analyze the results of clustering algorithms independently of data visualization.

Its analytical benefits aside, density-preserving visualization, as our results illustrate, is a more faithful representation of the underlying structure of the dataset. Even as the community becomes increasingly aware of the intricate limitations of existing visualization tools, inaccurate visualizations will continue to expose researchers to potential biases in data interpretation. A large body of work in the social sciences highlights the problematic nature of inaccurate visualizations: for example, even though distortions in Mercator projections of the world map are well-known, they still suggest biased conclusions to viewers^35, 36^. Our suite of density-preserving visualization tools will reduce such distortions and can help prevent unintentional biases and misdirection when researchers interpret and share insights from these data.

Our work motivates a number of interesting methodological directions of further research for analyzing single-cell datasets. First, the systematic changes in transcriptomic variability we discovered in tumor-infiltrating immune cells suggest *differential variability* as a general tool for characterizing the biological differences between different cell populations. A change in transcriptomic variability likely reflects underlying changes in gene regulatory mechanisms, and identifying the key drivers of this pattern and studying their roles merits further exploration. Our visualizations also motivate local density measures for noise reduction, as they often reveal fine-grain structure within a cell type, typically a dense “core” surrounded by a sparse cloud of cells with more divergent expression patterns. By focusing on only this core, one could obtain crisper canonical representations of individual cell states and developmental trajectories. Lastly, given the popularity of *k*-nearest neighbor-based methods in scRNA-seq analysis^1^ our results suggest that many popular tools may also benefit from information about local variability. It would be interesting to explore density-augmented algorithms for clustering, trajectory analysis, and data integration that could better exploit the true underlying structure of the data to enhance the accuracy of downstream analyses. Our work represents a key step forward in understanding the dynamic structure of complex single-cell transcriptomic landscapes.

## Methods

### Review of t-SNE and UMAP

The most widely-used nonlinear visualization algorithms in single-cell transcriptomic analysis are t-SNE^3^ and UMAP^4^, and both follow a similar methodology. They first compute a nearest-neighbor graph of the high-dimensional data and introduce a probability distribution on the edges of this graph that assigns larger weights on smaller distances. They then choose an embedding that minimizes the distance between this original probability distribution and a similar distribution computed on the embedding. The key differences between the two algorithms lie in their choices of these distributions and the objective function quantifying the difference between the two distributions.

Let 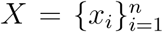 be our input dataset with *n* data points, where each *x*_*i*_ ∈ ℝ^*d*^ (e.g. gene expression profile of a cell). Let *E* be the set of edges (*i, j*) in the (directed) *k*-nearest neighbor graph constructed on this dataset, where *j* is one of the *k* points closest to *i*. For t-SNE, the probability distribution on the original data, 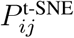, is given by normalizing and symmetrizing Gaussian kernel distances:

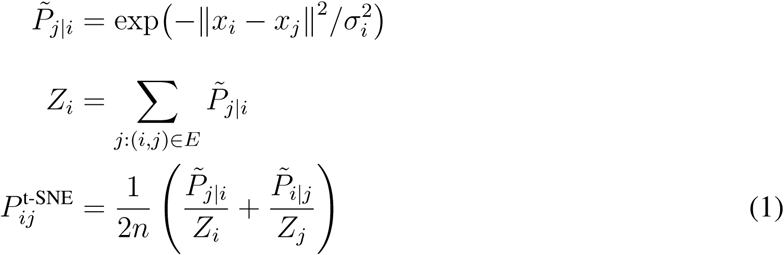

where *σ*_*i*_ is chosen adaptively for each *i* and corresponds the length-scale at *x*_*i*_.

UMAP uses a slightly different kernel, representing a rescaled exponential distribution:

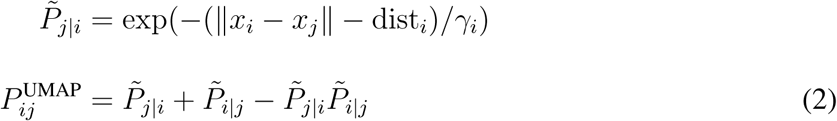

where *γ*_*i*_ is chosen adaptively and also corresponds to the length-scale, and dist_*i*_ is the distance from *x*_*i*_ to its nearest neighbor. We expand on the role of *σ*_*i*_ and *γ*_*i*_ in the next section.

For the probability distributions computed on the embedding, both t-SNE and UMAP use a heavy-tailed distribution (e.g. Student’s *t*-distribution for t-SNE), which emphasizes preserving local structure in the original dataset while being more lenient towards longer distances (see the original papers^3, 4^ for a thorough explanation). Formally, the probability distributions 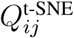 and 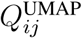 in the embedding are defined as

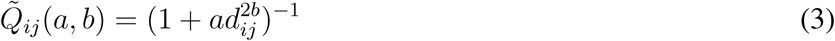

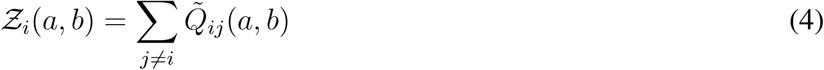

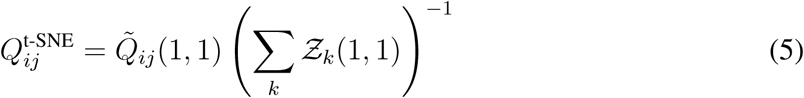

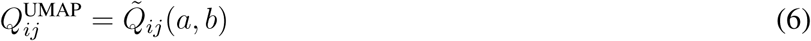

where *d*_*ij*_ represents the distance between points *i* and *j* in the embedding (Euclidean for both methods), and *a* and *b* are additional shape parameters UMAP introduces to control the spread of the distribution according to a user parameter. In the following, we omit the superscripts of *P* and *Q* when they are clear from the context.

The goal of both algorithms is to generate an embedding that minimizes the difference between *P* and *Q*. The loss function used by t-SNE to quantify this difference is the Kullback-Leibler (KL) divergence:

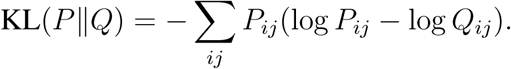

UMAP instead uses the cross-entropy (CE) loss summed over all the edges:

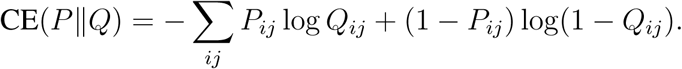

Both methods optimize the embedding coordinates to minimize the respective loss functions using standard gradient descent optimization techniques (see Supplementary Note 2 for details). Notably, the fact that UMAP does not require *Q* to be renormalized over all edges allows UMAP to use *stochastic* gradient descent (whereby the embedding coordinates are updated for one data point at a time), making it more computationally efficient than t-SNE in general.

### Adaptive length-scale selection in t-SNE and UMAP erases density information

The length-scale parameters *σ*_*i*_ and *γ*_*i*_ play an important role. The exponentially-decaying tails of the *P* distribution in both t-SNE and UMAP mean that the points a few multiples of the length-scale away from *x*_*i*_ are effectively omitted from the conditional distribution *P*_*·|i*_. Thus, the choice of the length-scale at point *x*_*i*_ determines the radius of the local structure around *x*_*i*_ that the embedding aims to preserve. Since different points in the dataset can have vastly different distribution of distances to their respective nearest neighbors, it is desirable to use a different *σ*_*i*_ or *γ*_*i*_ for each point *x*_*i*_ in order to evenly capture the local structure across all parts of the data.

In t-SNE, the *σ*_*i*_’s are chosen by setting the *perplexity* of each conditional distribution *P*_*·|I*_ constant. Perplexity can be thought of as a “smooth” analog of the number of nearest neighbors and is formally defined as 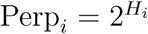, where *H*_*i*_ denotes the entropy of the conditional distribution *P*_·|*i*_:

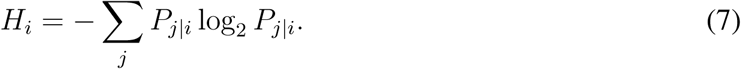

Since perplexity monotonically increases in *σ*_*i*_ (more points are significantly represented in *P*_*·|i*_ as *σ*_*i*_ increases), t-SNE performs a binary search on each *σ*_*i*_ to obtain a constant perplexity for all *i*. UMAP’s length-scale selection is analogous, but instead of fixing the value of perplexity, it fixes the marginal sum of probabilities at each point *i*, Σ_*j*_ *P*_*ij*_, by choosing an appropriate *γ*_*i*_.

Although it is effective for capturing local structure, adaptive choice of length-scale has the undesirable consequence of canceling out differences in density around each point in the original data, as t-SNE (implicitly) and UMAP (explicitly) both assume the data points are distributed uniformly on an underlying manifold. Note that, in both t-SNE and UMAP, a sparse neighborhood of *x*_*i*_ leads to a large length-scale, whereas a dense neighborhood leads to a small length-scale. Since the distance between points is divided by the length-scale parameter in the computation of *P*, we can intuitively see that this normalization removes density information from the data.

More formally, consider a dataset of points 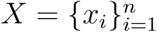 with Euclidean pairwise distances. 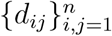 Suppose we dilate the data space by a factor of *α >* 1 to generate a sparser dataset 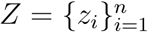 with the same underlying structure, where the new pairwise distances are scaled by *α*, i.e. ∥*z*_*i*_ − *z*_*j*_∥ = *αd*_*ij*_. A key observation is that the distribution *P* computed on *X* by t-SNE or UMAP will be *identical* to *P* computed on *Z*, even though *Z* represents a more heterogeneous set of points than *X*. Intuitively, this is because obtaining the same perplexity/marginal sum of probabilities on *Z* requires that the respective length-scales be scaled by *α*, which cancels out the increase in distances and leaves the resulting *P unchanged*. Since *P* is the only information about the dataset provided as input to the embedding step of each algorithm, the original differences in density in different regions of the data space are entirely lost in the embedding. We provide a more detailed description of this property and its generalization to a broader class of generative models for the underlying data in Supplementary Note 3.

### Our approach: capturing density information using the local radius

To generate embeddings that retain information about the density at each point, we introduce the notion of a *local radius* to make concrete our intuition of spatial density. Intuitively, a point is in a *dense* region if its nearest neighbors are very close to it, and in a *sparse* region if its nearest neighbors are far away. Thus, we use average distance to nearest neighbors as a measure of density for a given point.

To formalize this notion, for a point *x*_*i*_, we require two components: (i) a pairwise distance function *d*(*x*_*i*_, *x*_*j*_), and (ii) a probability distribution *ρ*_*j|i*_ that weighs each *x*_*j*_ based on its distance from *x*_*i*_, with faraway points having lower weights. We define the local radius at *x*_*i*_, denoted *R*_*ρ*_(*x*_*i*_), as the expectation of the distance function over *x*_*j*_ with respect to *ρ*_*j|i*_, thus capturing the average distance from *x*_*i*_ to nearby points:

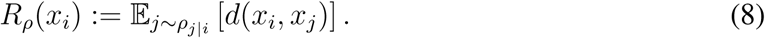

In the following, we let the distance function be the squared Euclidean distance, i.e. *d*(*x*_*i*_, *x*_*j*_) = ∥*x*_*i*_ − *x*_*j*_∥^2^. Other choices of distance function can be easily incorporated into our framework.

In den-SNE and densMAP, we take advantage of the probability distributions *P* ^t-SNE^ and *P* ^UMAP^ which already capture local relationships; for the local radius in the original embedding, we set *ρ*_*j|i*_ = *P*_*ij*_*/* Σ_*j*_ *P*_*ij*_ to get

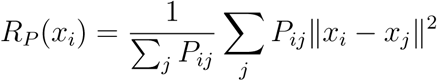

for both methods. Note that *P* vanishes rapidly outside the neighborhood of each *x*_*i*_ and is thus well-suited for density estimation. We can show in fact that this representation of density (inversely-related) has the desirable property that it scales with the *variance* of a range of data-generating distributions and increases when the length-scale term *σ*_*i*_ increases (Supplementary Note 3).

Next, we define the local radius in the embedding. Let *y*_*i*_ be the embedding coordinates of the point *x*_*i*_ given by the algorithm of choice. We need a distribution analogous to *P* for calculating the expected distance between *y*_*i*_ and its neighbors in the embedding. Although it would be ideal for this distribution to have adaptive length-scales like *P*, this would present a major hurdle for optimization because the binary search used to determine *σ*_*i*_ and *γ*_*i*_ is not differentiable. Instead, we leverage the embedding distribution *Q* computed by t-SNE and UMAP as an approximation for the adaptive scheme. It is worth noting that, in the case of t-SNE, *Q* is based on a Cauchy distribution, which can be interpreted as the marginalization of a Gaussian distribution over an unknown variance^37^. Thus, *Q* intuitively reflects an average over all length-scales. Letting *ρ*_*j|i*_ = *Q*_*ij*_*/* Σ_*j*_ *Q*_*ij*_ and *d*(*y*_*i*_, *y*_*j*_) = ∥*y*_*i*_ − *y*_*j*_∥^2^, the local radius in the embedding is given as

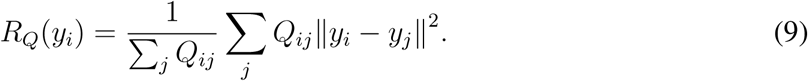

For ease of notation, we denote the local radius in the original data as *R*_*o*_ and the local radius in the embedding as *R*_*e*_ in the following sections.

### Augmenting the visualization objective to induce density preservation

To preserve density, we aim for a *power-law* relationship between the local radius in the original dataset and in the embedding, i.e. *R*_*e*_(*y*_*i*_) ≈ *α* [*R*_*o*_(*x*_*i*_)]^*β*^ for some *α* and *β*, inspired by the exponential scaling of density with respect to dimensionality (see Supplementary Note 1). This can be reframed as an *affine* relationship between the logarithms of the local radii, i.e.,

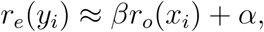

where we define *r* _*o*_(*x*_*i*_) : = log *R* _*o*_(*x*_*i*_) a nd *r* _*e*_(*y*_*i*_) : = log *R* _*e*_(*y*_*i*_). T he g oodness-of-fit of this relationship can be measured by the *correlation coefficient*

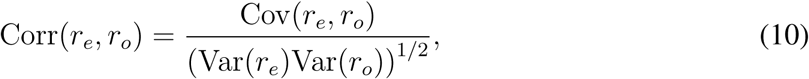

which is invariant to the parameters *α* and *β*. Cov(*·, ·*) denotes the covariance function, and Var(*·*) denotes the variance function. Note that these quantities are estimated by considering the tuples 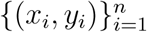 as *n* independent samples from the same distribution; e.g., the mean of *r*_*e*_ is estimated as 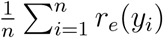.

Our density-preservation objective is to choose the embedding 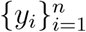 such that correlation between the log local radii of the original dataset and the embedding is maximized. This approach is closely related to canonical correlation analysis^38^ (CCA), which finds a linear transformation of a dataset that maximizes its correlation with another. We are further motivated by recent work that extends CCA to nonlinear transformations^39^.

Augmenting the loss functions of t-SNE and UMAP with this density-preservation objective yields the den-SNE and densMAP objectives, respectively:

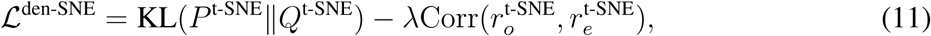

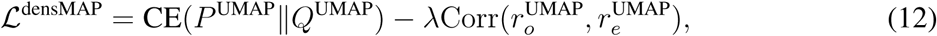

where *λ* is a user-chosen parameter that determines the relative importance of the density-preservation term compared to the original objective.

### Optimizing the embedding with respect to density-augmented objectives

Our differentiable formulation of the local radius enables us to optimize the density-augmented objective functions (11) and (12) using standard gradient descent techniques. Since both t-SNE and UMAP are also based on gradient descent, it suffices for us to calculate the contribution of the density-preservation objective to the overall gradient and add it to the existing t-SNE and UMAP gradients.

The gradient of the density-preservation objective with respect to the embedding coordinates *y*_*i*_ is given by

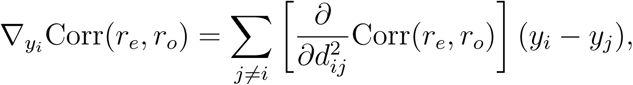

where *d*_*ij*_ = ∥*y*_*i*_ − *y*_*j*_∥. To simplify the notation, let 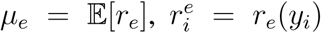, and 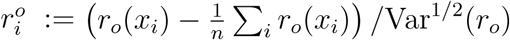. Note that the centering of 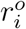 and normalizing by standard deviation does not depend on the embedding and thus can be precomputed. Now, the inner gradient term with respect to 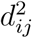 can be calculated as

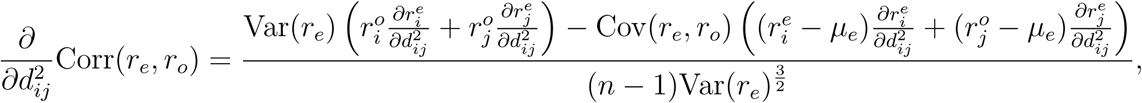

where

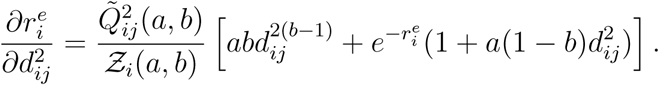

The terms 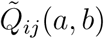 and *Z*_*i*_(*a, b*), defined in (3) and (4), respectively, are quantities computed by t-SNE and UMAP to capture the local structure of the embedding. (*Z*_*i*_(*a, b*) is required only in t-SNE.) Setting the parameters *a* = *b* = 1 results in the t-SNE formulation, whereas UMAP sets these two parameters as a function of a user parameter. A detailed derivation of our gradients above is provided in Supplementary Note 2.

Optimizing the densMAP objective requires special consideration because UMAP uses stochastic gradient descent (SGD), whereby edges are sampled according to *P*_*ij*_ and the gradient update is performed for one edge at a time. Since the gradient formula (10) involves a sum over its neighbors with equal weights, edges sampled from *P* must be re-weighted to obtain unbiased estimates of our gradient. To this end, we multiply the density term in the gradient for an edge {*i, j*} by *Z/nP*_*ij*_ where *Z* = Σ_{*k,l*, }∈*E*_ *P*_*kl*_, to correct for sampling bias. In addition, there are a number of global terms that are computationally burdensome to update for every edge, which include Var(*r*_*e*_), Cov(*r*_*e*_, *r*_*o*_), and *µ*_*e*_. We compute these terms in the beginning of each epoch (a round of edge-wise updates for the entire dataset) and consider them as fixed during the epoch. This can be viewed as a form of coordinate descent, where the objective is optimized with respect to a subset of variables at a time while conditioning on the rest. We describe these techniques in detail in Supplementary Note 2.

### Implementation details

To ensure that our methods find good local optima of (11) and (12) that are as effective as t-SNE and UMAP in separating clusters, we take a two-step approach where we run the original algorithms *without* the density-preserving objective for the first *q* fraction of iterations, then optimize the full objective for the remaining 1−*q* fraction of iterations. This approach is akin to t-SNE’s “early exaggeration”, whereby the first several iterations of the optimization emphasize attractive forces to help guide the direction of the optimization.

For computational efficiency, we approximate the embedding distribution *Q* used in our local radius computation (9) by allowing *Q*_*ij*_ to be non-zero only when *P*_*ij*_ is non-zero (i.e. *i* and *j* are *k*-nearest neighbors in the original space), thus inducing sparsity in *Q*. This technique is especially well-suited for the aforementioned two-step approach, since the embedding already closely follows the nearest-neighbor structure in *P* when this approximation takes effect.

There are several parameters of den-SNE and densMAP that the users can modify to tailor the behavior of these algorithms. We inherit all of the parameters from t-SNE and UMAP, including perplexity (t-SNE) or number of neighbors (UMAP), number of iterations/epochs, and the “mindist” parameter for UMAP (which controls the *a* and *b* parameters in *Q*_*ij*_; see (6)). We refer the readers to the original publications for a detailed discussion of these parameters. There are two additional parameters we introduce in den-SNE and densMAP: the weight *λ* ≥ 0 given to the density-preserving objective, and the fraction *q* ∈ [0, 1] of iterations that take the density term into account. All of our experimental results are based on the following default parameter settings that we recommend. For den-SNE, we use perplexity of 50 and 1000 iterations (same as the default setting of t-SNE), along with *q* = 0.3 and *λ* = 0.1. For densMAP, we use 30 neighbors, 750 epochs, *q* = 0.3, and *λ* = 2. We note that changing the value of *λ* leads to qualitatively different embeddings that achieve different trade-offs between the original visualization objective and the density-preservation term (Supplementary Figure 14).

### Quantitative evaluation of density preservation

To assess the performance of visualization algorithms at preserving density, we compute the correlation between the log local radii in the original dataset and two measures of visual density in the embedding generated by the algorithm.

The first measure of visual density is the local radius computed in the same manner as in the original space. Recall that during the optimization, we compute the local radius in the embedding *approximately* using the heavy-tailed distribution *Q* computed by t-SNE or UMAP and consider only the edges present in the nearest-neighbors graph of the original data. For accurate evaluation, here we compute the local radius more directly as follows. Given the embedding points 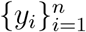, we compute the analog of the *P* matrix on the original data on these embedding points, denoted *P*. For t-SNE and den-SNE, we define *P* as: where *σ*_*i*_, the length-scale parameter, is chosen to achieve the same perplexity as in the original *P′* matrix. For UMAP and densMAP, we define *P′* as:

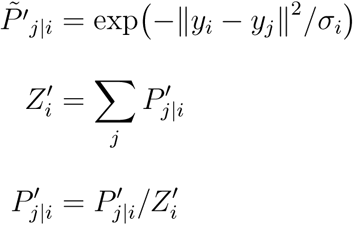

where 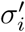, the length-scale parameter, is chosen to achieve the same perplexity as in the original *P* matrix.

For UMAP and densMAP, we define *P′* as:

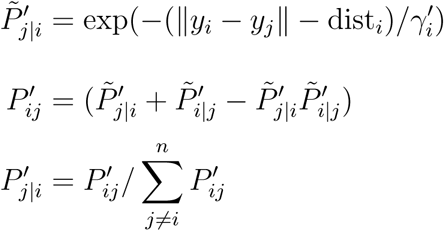

where dist_*i*_ is the distance to the nearest neighbor of *y*_*i*_, and 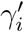 is chosen to achieve the same constant marginal 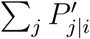 as the original *P* matrix.

Since *P′* more explicitly focuses on the local neighborhoods of points in the embedding than *Q* by adaptively choosing the length-scale, calculating the local radius using this distribution more accurately reflects the actual density of each point in the embedding:

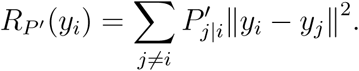

Our quantitative metric of density preservation is then the Pearson correlation coefficient (*R*^2^) between log *R*_*P*_ ′(*y*_*i*_) and *r*_*o*_(*x*_*i*_) = log *R*_*P*_ (*x*_*i*_), where the latter is the log local radius in the original data space.

The second measure of visual density in the embedding is the *neighborhood count*, which is motivated by the visual perception of density as the number of points in a given area. For a point *y*_*i*_ in the embedding and a radius *l*, the *l*-neighborhood count of *y*_*i*_ is the number of points *y*_*j*_ that are within a distance of *l* from *y*_*i*_ in the embedding. Thus, dense regions will have large neighborhood counts and sparse regions, small counts. This natural notion of local density has been extensively used in the psychology of vision^10, 40^.

To systematically choose *l* for each dataset, we first compute the area *A* of the smallest bounding box of the embedded points, then calculate an average length-scale 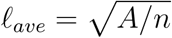, where *n* is the number of points in the dataset. To assess density preservation across different length-scales, we tested different multiples of *l*_*ave*_; for den-SNE and t-SNE, we chose *l* from {*l*_*ave*_, 2*l*_*ave*_, 4*l*_*ave*_}, and for densMAP and UMAP, from 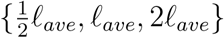. We chose smaller values for densMAP and UMAP because those embeddings are more compact in general than those of den-SNE and t-SNE for our parameter choices. For each chosen *l*, we calculate the *l*-neighborhood count for each point in the embedding and calculate the correlation (in log space) with the local radii in the original space as a quantitative metric of density preservation. A strong negative correlation is desirable, which indicates that points with a higher neighborhood count (higher visual density) tend to have a smaller local radius in the original dataset (smaller underlying variability).

### Data preprocessing

We obtained three publicly available scRNA-seq datasets for the main analyses: a dataset of immune cells in lung cancer and blood^7^, a dataset of peripheral blood mononuclear cells (PBMCs) in healthy individuals^8^, and a dataset that profiled the developmental trajectory of *C. elegans*^9^. We used three additional scRNA-seq datasets for validation experiments, including another lung cancer dataset^14^ and two blood immune cell datasets^19, 20^. For each dataset, we applied the same cell and gene filtering schemes used by the original publications, then normalized the data so that each cell has the same total number of counts (10k). Following the standard in scRNA-seq analysis, we then log-transformed the normalized counts, i.e. *x* → log(1 + *x*). Principal component analysis (PCA) was then used to produce lower-dimensional representations of individual cells, which are provided as input to the visualization algorithms. We used the number of principal com-ponents (PCs) prescribed by the original publications if present, or used 50 dimensions otherwise. The resulting datasets for the main experiments included 48,969 cells and 306 PCs for lung cancer, 68,551 cells and 50 PCs for PBMCs, and 86,024 cells and 100 PCs for *C. elegans*. We used the cell type labels provided by the original datasets for visualization.

For the UK Biobank dataset^26^, we used the 40 PC loadings provided as part of the genetic data for visualization. We analyzed a 20% subsample of the dataset including 97,676 individuals, for computational efficiency. Ethnicity labels for the individuals were obtained from Data Field 21000, which was collected from the participants via a touchscreen questionnaire. To visualize subpopulation structure within the white British individuals, we performed spectral clustering using the 40 PCs as input to identify five subclusters.

For the MNIST dataset, we flattened each of the 60,000 28*×* 28 pixel images to a 784-dimensional vector and used the top 50 PCs as our input to the visualization algorithms. Labels classifying the handwritten digits were provided in the dataset.

### Differential analysis of gene expression variability in the lung cancer data

For each cell type with visible expansion of transcriptomic variability in tumor in our visualizations (memory B cells, naïve B cells, CD4 memory resting T cells, CD4 naïve T cells, and CD8 T cells), we identified twenty genes with the largest increase in variance in tumor compared to blood for further analysis. For each gene and cell type, we calculated the differences in the mean and variance of expression between tumor and blood. The statistical significance of the observed differences is assessed using a permutation test, whereby the assignment of cells to tumor or blood is randomly permuted, and the statistic computed on the permuted dataset is viewed as samples from the null distribution where there is no difference between tumor and blood. For comparing the variance, we centered the expression levels for each group (tumor or blood) before the permutation procedure to control for the shift in mean. The *p*-value is calculated as the fraction of permutations that result in a statistic whose magnitude is larger than the statistic computed on the original dataset. We used 100k permutations to estimate the *p*-values and applied Bonferroni correction within each cell type to account for multiple hypothesis testing.

When considering changes in the variance of gene expression, it is important to note that an increase in variance can often be explained by an increase in mean. For example, under the Poisson process model of underlying count distributions, variance of the observed counts naturally scales with the mean^34^. Thus, we additionally calculated the difference in *dispersion index* to assess the extent to which the change in variance is unexplained by a corresponding change in mean. The dispersion index (DI) is given by *σ*^2^*/µ*, where *µ* and *σ*^2^ are the mean and variance of expression. We assessed the statistical significance of the difference in DI also using a permutation test. For the null distribution, we assume that in the absence of excess difference in dispersion, the variance of expression has a linear dependence on the mean (as suggested by the dispersion index). A permutation scheme that correctly reflects this null distribution is one where the expression levels within each group (tumor or blood) are transformed as *x* → *µ*^−1*/*2^(*x* − *µ*)+1 before the permutation, where *µ* is the sample mean of the group. This transformation maps both groups to the same mean (*µ* = 1) while preserving the DI, so that permuting the labels leads to a valid sample from the null distribution. Similar to the mean and variance tests, we used 100k permutations to estimate the *p* values and applied Bonferroni correction.

### Assessing significance of density differences in monocytes and dendritic cells

To verify our claims that classical (CD14+) monocytes have more variability of expression than both CD16+ monocytes and DC3 dendritic cells (as characterized by the PBMC2 dataset), we compared the distribution of the log local radius in the original data for each of these cell types in the PBMC2 and PBMC3 datasets. To assess significance, we used the one-sided Mann-Whitney U (MWU) test^41^, which tests the hypothesis that values drawn from one distribution are larger than those drawn from another. We calculated the MWU test statistic for: CD14+ monocytes and CD16+ monocytes in the PBMC2 and PBMC3 datasets; and for CD14+ monocytes and DC3 dendritic cells in PBMC2.

### Runtime and memory benchmarking

To evaluate runtime and memory usage of our density-preserving visualization methods, we used the *C. elegans* dataset from Packer et al. (2019) with 86,024 cells, which is the largest scRNA-seq dataset used in this paper. In addition to the full dataset, we subsampled it into smaller datasets, including 43,012 cells, 21,506 cells, 10,753 cells, and 5,376 cells. We measured the runtimes of denSNE, densMAP, t-SNE, and UMAP on each of the datasets with the default parameter settings and profiled memory usage using the psrecord package (https://github.com/astrofrog/psrecord). All experiments were run on an Intel Xeon Gold 6130 (2.30 GHz) processor and used a single core.

### Data availability

The lung cancer^7^ and *C. elegans*^9^ datasets are available from the Gene Expression Omnibus (GEO) database with accession numbers GSE127465 and GSE126954, respectively. The PBMC dataset^8^ is available from 10x Genomics at: https://support.10xgenomics.com/single-cell-gene-expression/datasets. For our validation datasets, the secondary lung cancer dataset^14^ is available from GEO (GSE99254), and the PBMC2^19^ and PBMC3^20^ datasets can be accessed through the Broad Institute’s Single Cell Portal (https://singlecell.broadinstitute.org/) with dataset IDs SCP43 and SCP345, respectively. Data access applications for the UK Biobank data can be submitted at: https://www.ukbiobank.ac.uk/. The MNIST dataset is available at: http://yann.lecun.com/exdb/mnist/. We also provide our preprocessed data for the main datasets (lung cancer, PBMC, and *C. elegans*) at: http://densvis.csail.mit.edu/datasets.

### Code availability

We provide the software for den-SNE and densMAP in the densVis package available at: http://densvis.csail.mit.edu/ and https://github.com/hhcho/densvis.

## Acknowledgements

The authors thank Brian Hie, Benjamin DeMeo, Ellen Zhong, and Joshua Peters for helpful discussions. Our analysis of genotype data was conducted using the UK Biobank Resource under Application Number 46341. BioRender.com was used to generate Figure 4d.

## Competing Interests

The authors declare that they have no competing financial interests.

## Notes

### Competing Interest Statement

The authors have declared no competing interest.

