## Supplementary Materials for "Density-Preserving Data Visualization Unveils Dynamic Patterns of Single-Cell Transcriptomic Variability"

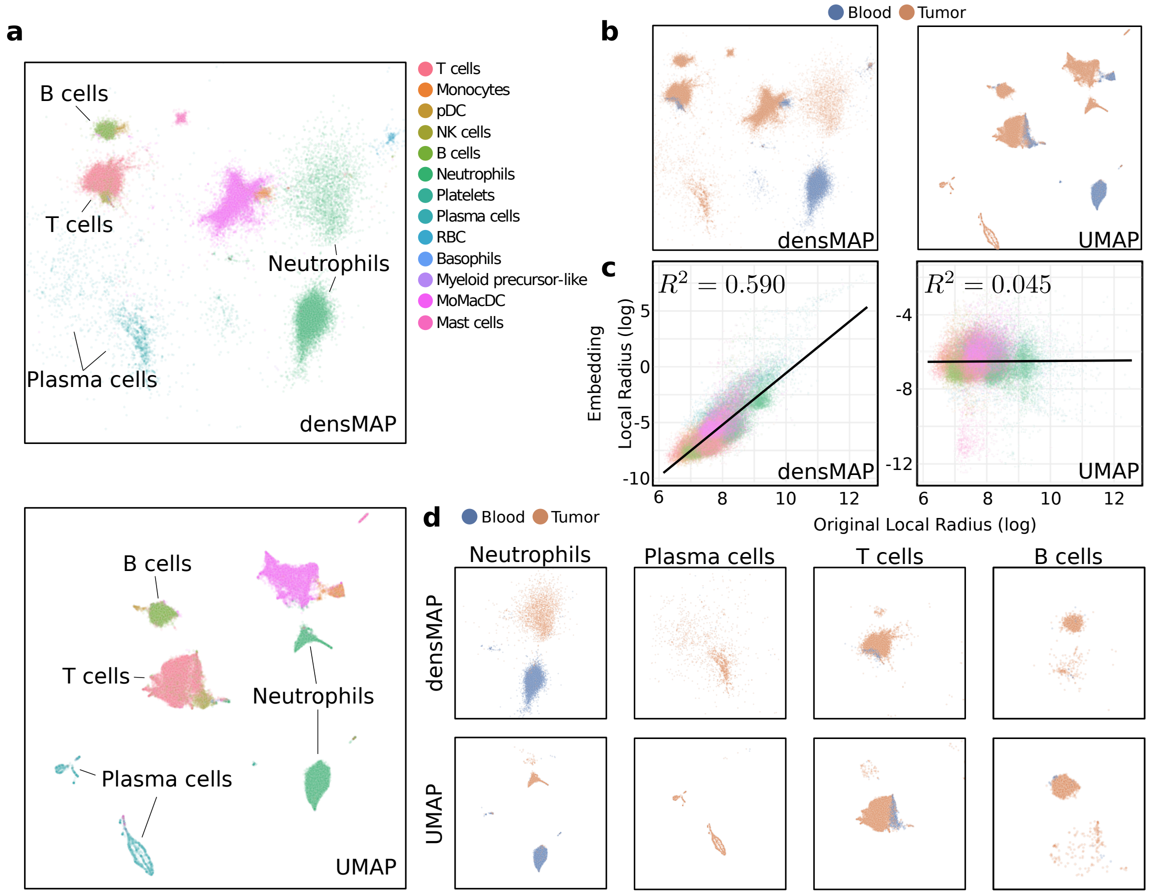


Supplementary Figure 1: **Visualizing lung cancer using densMAP recapitulates den-SNE results.** We repeat the analysis presented in Figure 3 using densMAP and UMAP. **a.** Top is a densMAP embedding and bottom is a UMAP embedding; points are colored by cell type. Note that the relative heterogeneity of neutrophils, plasma cells, and T cells are misleadingly portrayed in the UMAP visualization. **b.** densMAP (left) and UMAP (right) plots, now colored by tissue type (blood or tumor). **c.** Scatter plots comparing the local radii, our measure of local density (Methods), in the original space and in the visualization (embedding). Points are colored by cell type, and the $R^2$ value of the correlation is shown for each plot. Higher correlation of densMAP shows that densMAP more accurately conveys the density landscape of the original data than UMAP. Scatter plots based on neighborhood count (another measure of visual density; Methods) are included in Supplementary Figure 2. **d.** densMAP (top) and UMAP (bottom) plots restricted to each of four notable cell types (neutrophils, plasma cells, T cells, and B cells) and colored by tissue type (tumor or blood). Neutrophils and plasma cells in tumor considerably expand in size in densMAP, reflecting transcriptomic variability previously hidden in UMAP. T and B cells show a large increase in heterogeneity in tumor compared to blood in densMAP. Although UMAP shows a similar pattern, its lack of density-preservation property precludes reasoning about differences in heterogeneity.

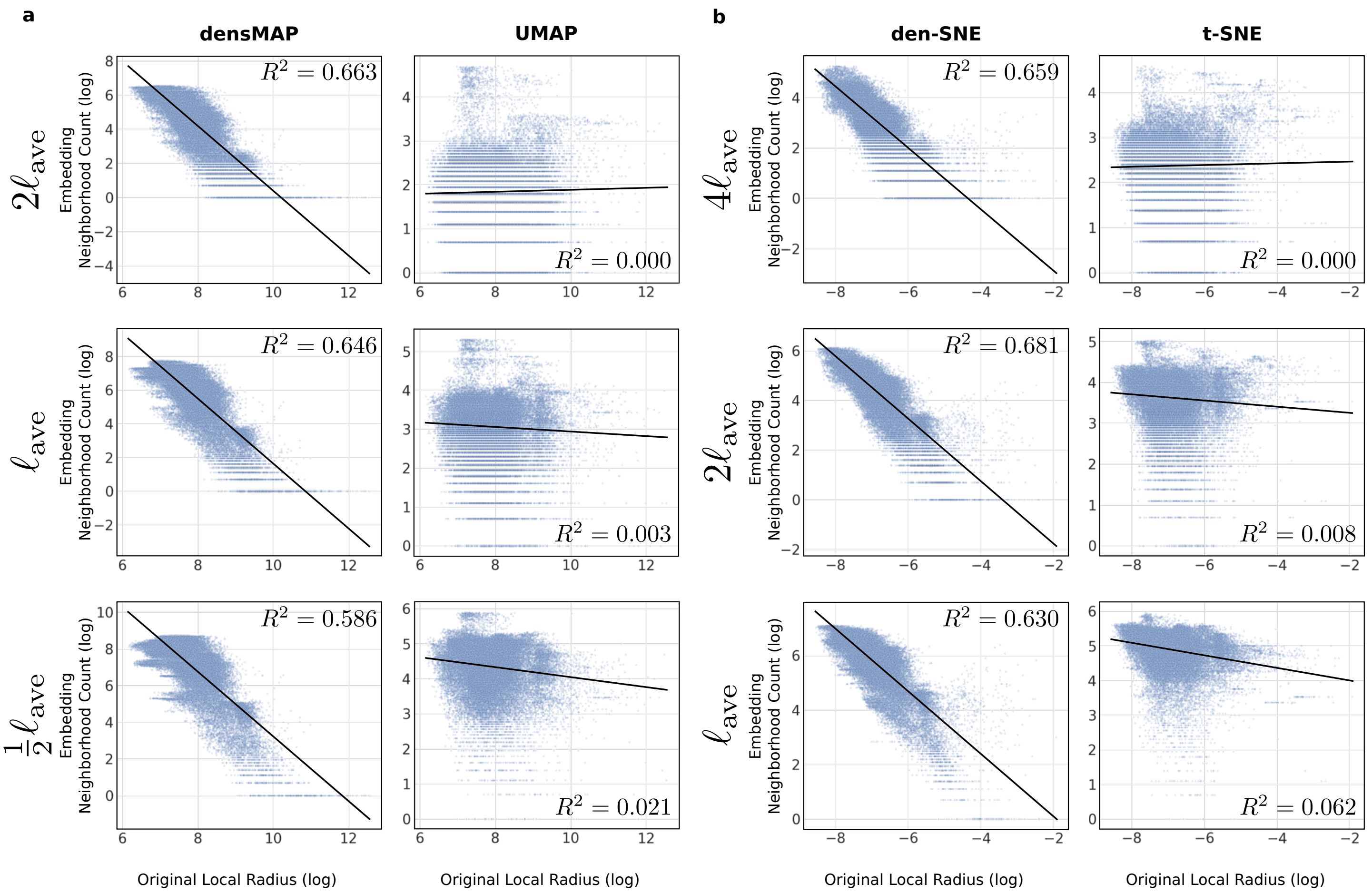


Supplementary Figure 2: **Density-preserving methods preserve density robustly at different scales on lung cancer data based on neighborhood count.** We compared the local radius of each point in the original lung cancer dataset to its neighborhood count in the visualizations (a measure of visual density; see Methods) for (**a**) densMAP and UMAP; and (**b**) den-SNE and t-SNE. We chose for each embedding, a length-scale $\ell_{\text{ave}}$ and multiples of that length-scale for which to compute the neighborhood counts (Methods): $\{\frac{1}{2}\ell_{\text{ave}}, \ell_{\text{ave}}, 2\ell_{\text{ave}}\}$ for densMAP and UMAP, and $\{\ell_{\text{ave}}, 2\ell_{\text{ave}}, 4\ell_{\text{ave}}\}$ for den-SNE and t-SNE. Since neighborhood count represents the density around a given point, a visualization that preserves density information will have higher neighborhood counts for points with smaller local radii in the original space. Note that this negative correlation is significantly stronger for our density-preserving tools than for t-SNE and UMAP, and this pattern holds across the different length-scales.

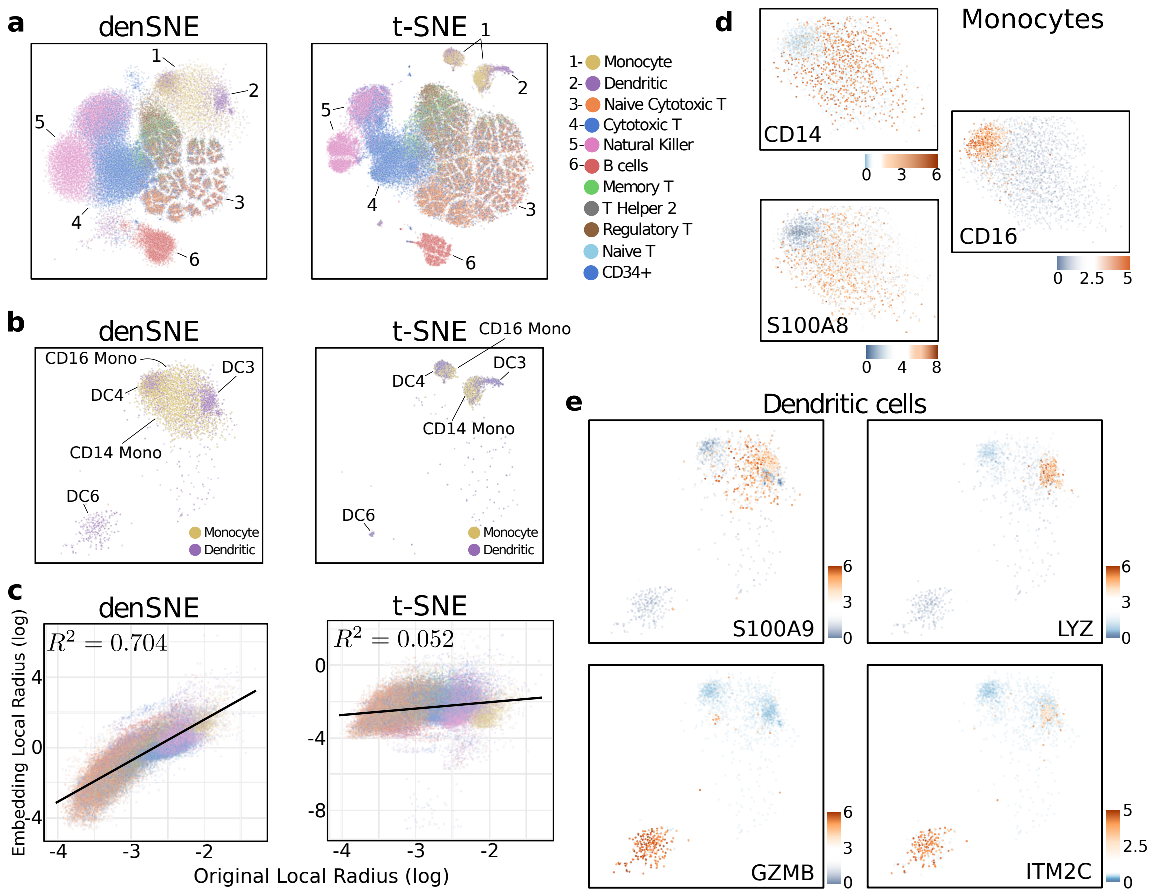


Supplementary Figure 3: **Visualizing PBMC data using den-SNE recapitulates densMAP results.** We repeat the analyses presented in Figure 4 using den-SNE and t-SNE. **a.** den-SNE (left) and t-SNE (right) visualizations of the data, colored by cell-type. The group of clusters corresponding to monocytes (cluster 1) and dendritic cells (DCs; cluster 2) showed the most pronounced difference between the two visualizations. **b.** For a detailed comparison, we plotted the same visualizations restricted to the monocyte-DC subset, which revealed distinct subtypes of monocytes (CD16 Mono and CD14 Mono) and DCs (DC3, DC4, and DC6) with clear density differences in den-SNE. Each subtype is annotated using the classification from the PBMC2 study (Villani et al., 2017; Methods) based on marker gene expression. Although the same subtypes are visible in t-SNE, their relative density differences are lost. **c.** Scatter plots comparing the local radii, our measure of local density (Methods), in the original space and in the visualization (embedding). Points are colored by cell type, and the $R^2$ value of the correlation is shown for each plot. Higher correlation of den-SNE shows that den-SNE more accurately conveys the density landscape of the original data than t-SNE. Scatter plots based on neighborhood count (another measure of visual density; Methods) are included in Supplementary Figure 4. **d.** Gene expression heatmaps of monocyte marker genes CD14, S100A8, and CD16 in the den-SNE visualization restricted to monocytes. The patterns of expression support our classification of the dense cluster as CD16 Mono and the sparse cluster as CD14 Mono. **e.** Gene expression heatmaps of DC marker genes from the PBMC2 study (Villani et al., 2017; Methods) for DC3 (top) and DC6 (bottom) in the densMAP visualization restricted to DCs. These support our assignment of DC clusters to DC3 and DC6.

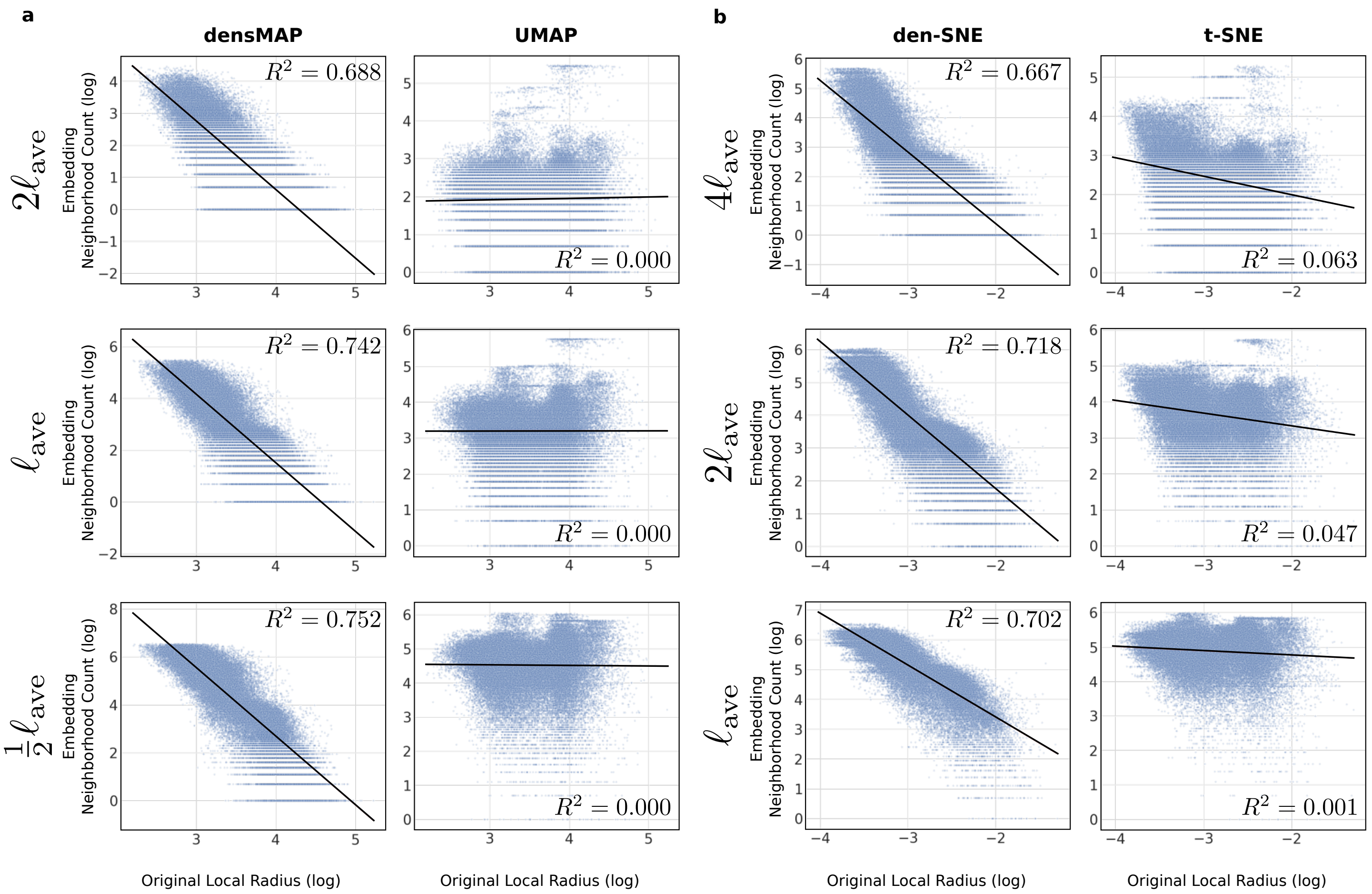


Supplementary Figure 4: **Density-preserving methods preserve density robustly at different scales on PBMC data based on neighborhood count.** We compared the local radius of each point in the original PBMC dataset to its neighborhood count in the visualizations (a measure of visual density; see Methods) for (**a**) densMAP and UMAP; and (**b**) den-SNE and t-SNE. We chose for each embedding, a length-scale $\ell_{\text{ave}}$ and multiples of that length-scale for which to compute the neighborhood counts (Methods): $\{\frac{1}{2}\ell_{\text{ave}}, \ell_{\text{ave}}, 2\ell_{\text{ave}}\}$ for densMAP and UMAP, and $\{\ell_{\text{ave}}, 2\ell_{\text{ave}}, 4\ell_{\text{ave}}\}$ for den-SNE and t-SNE. Since neighborhood count represents the density around a given point, a visualization that preserves density information will have higher neighborhood counts for points with smaller local radii in the original space. Note that this negative correlation is significantly stronger for our density-preserving tools than for t-SNE and UMAP, and this pattern holds across the different length-scales.

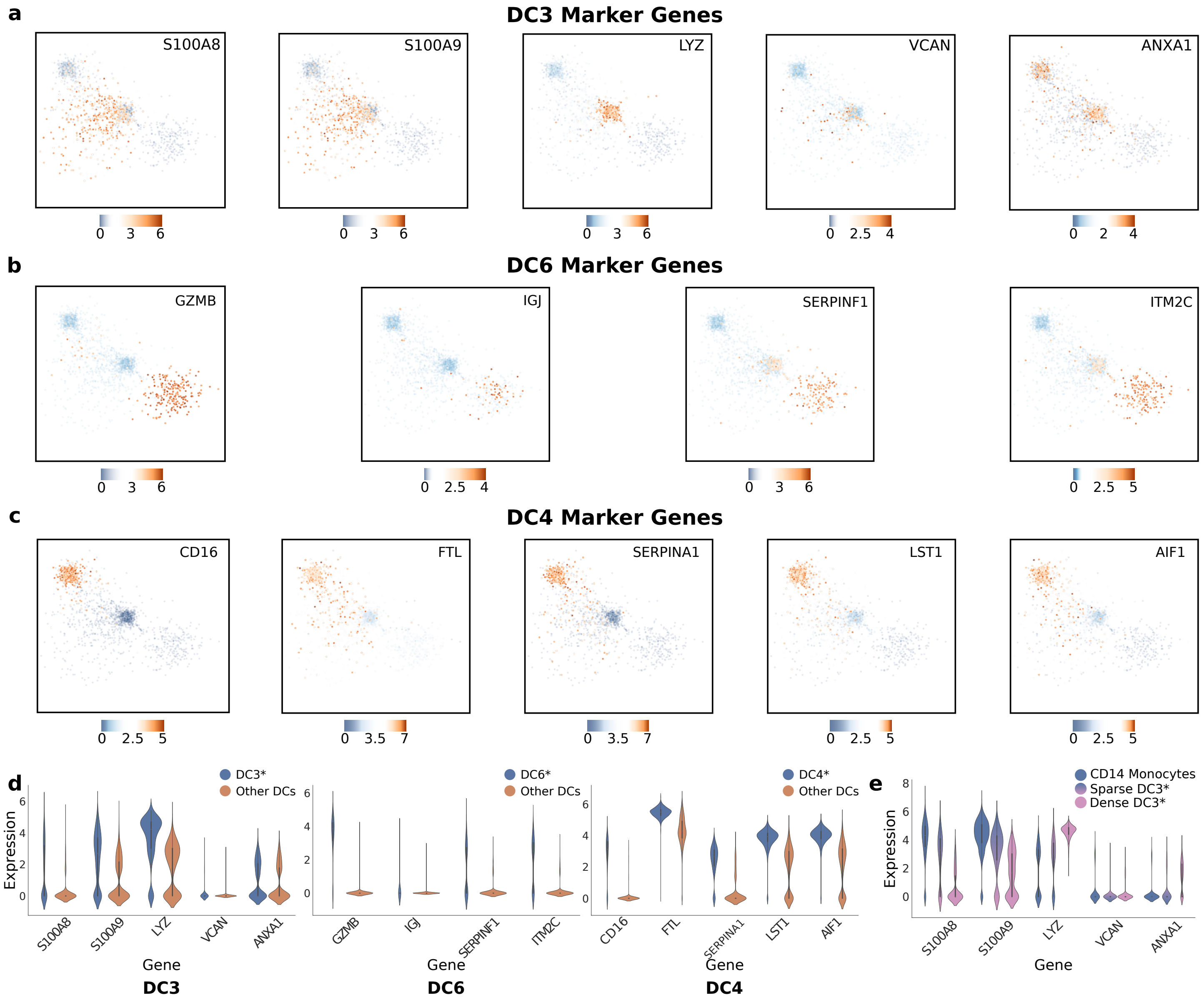


Supplementary Figure 5: **Marker gene expressions for dendritic cell subtypes in the PBMC dataset.** We plot gene expression heatmaps for the marker genes identified in the original study of PBMC2 for the dendritic cell (DC) subtypes (**a**) DC3, (**b**) DC6, and (**c**) DC4 on our densMAP visualization of the PBMC data, restricted to DCs. **d.** Violin plots showing the expression of marker genes identified by the PBMC2 study in our putative DC subtypes in the PBMC dataset: DC3 (left), DC6 (middle), and DC4 (right). The higher expression of these genes in our assigned subtypes and the expression patterns in (**a**) through (**c**) support our assignment of the DCs in the original PBMC dataset to the known subtypes in the PBMC2 dataset. The asterisk indicates our putative cell type assignment based on marker gene expression. **e.** Noting the existence of sparse and dense parts of the DC3 cluster in Figure 4, we compare the expression of DC3 and classical monocyte marker genes in the dense DCs (log local radius less than 3.9), sparse DCs (log local radius greater than 3.9), and classical monocytes; the violin plot indicates that sparse DCs are intermediate in expression of these marker genes between dense DCs and classical monocytes, potentially indicating a transition between the two states. The asterisk indicates our putative cell type assignment based on marker gene expression.

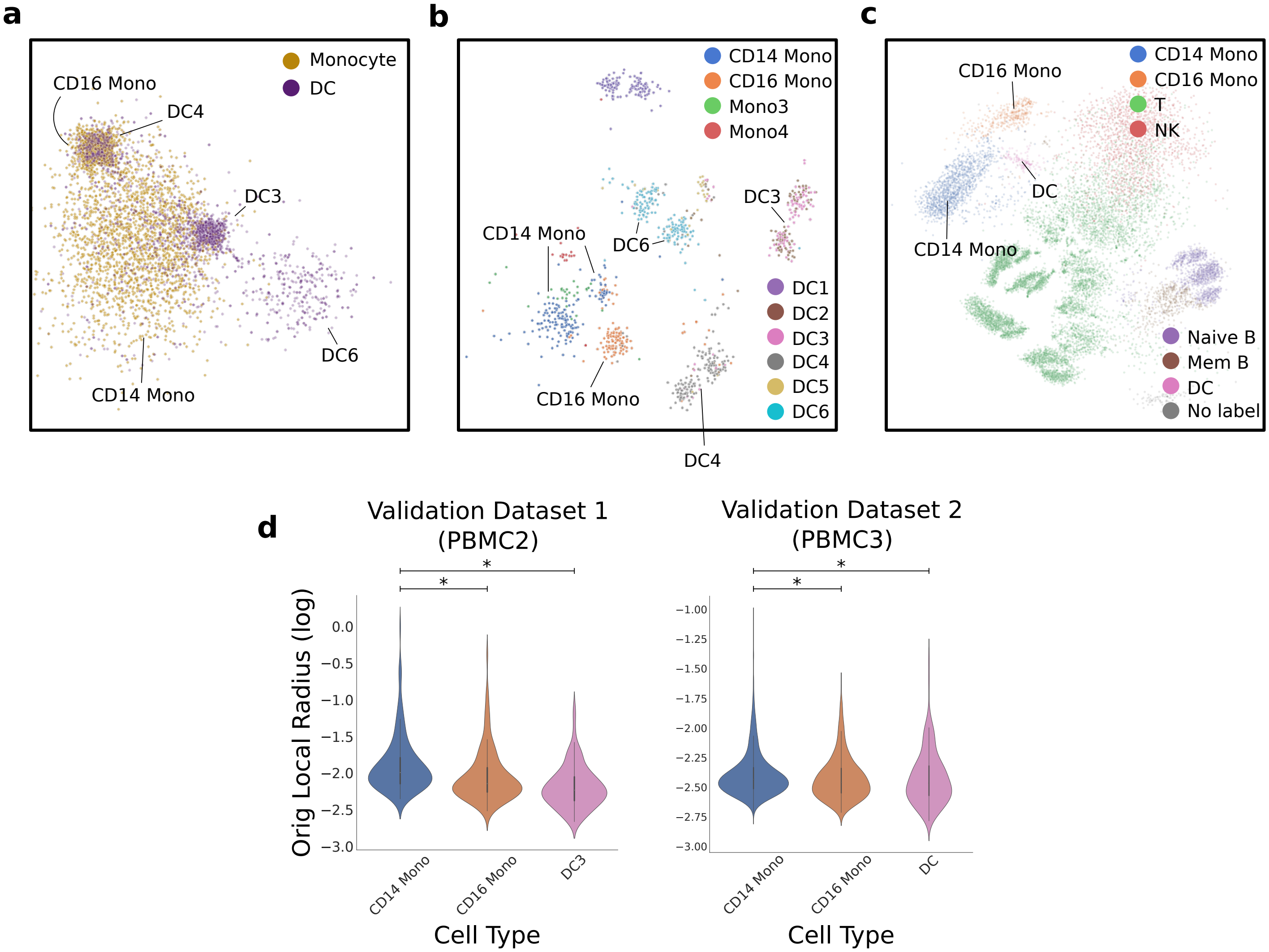


Supplementary Figure 6: **Density differences among monocytes and dendritic cell subtypes are validated on additional datasets.** Density-preserving visualizations of the (**a**) PBMC data, zoomed in on the monocytes and DCs (densMAP); (**b**) PBMC2 data (den-SNE); and (**c**) PBMC3 data (den-SNE) (Methods). In (**a**), the labels assigned are hypothesized based on subtypes determined in the PBMC2 dataset (see Supplementary Figure 5). The visualizations of PBMC2 and PBMC3 data recapitulate the density differences between CD14 Mono (which are sparse) and CD16 Mono/DC3 subsets (which are dense) observed in PBMC data. **d**. We further validated these density difference on the PBMC2 and PBMC3 data based on the original local radii computed for each of these datasets. For both PBMC2 (left) and PBMC3 (right), the local radius in the original data is significantly larger in CD14 monocytes than in CD16 monocytes. Similarly, DC3 has significantly smaller local radii than CD14 monocytes in the PBMC2 data; in the PBMC3 dataset, DC subtype labels were not available but the local radii of CD14 monocytes are still larger than that of the DCs. The $*$ indicates a $p$-value less than $5 \times 10^{-4}$ for a one-sided Mann-Whitney U test statistic (see Methods). NK: Natural killer cells; Mem B: Memory B cells

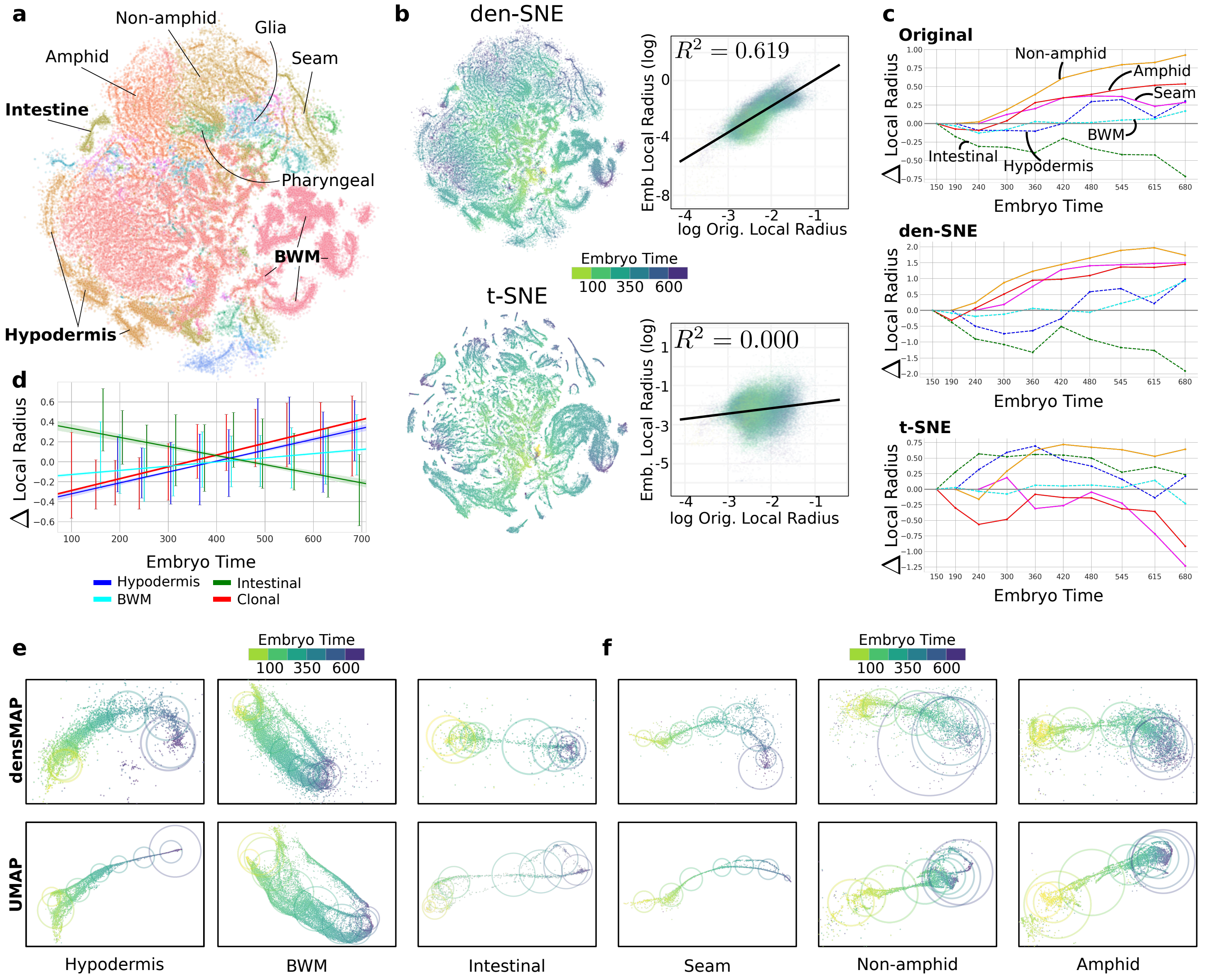


Supplementary Figure 7: **Visualizing the *C. elegans* embryo development data with den-SNE recapitulates densMAP results.** We repeat the analysis presented in Figure 5 using t-SNE and den-SNE. **a.** den-SNE embedding of dataset, colored by cell-type (t-SNE ommitted for space). **b.** Same data, now colored by embryo time, with den-SNE above and UMAP below. The scatter plots on the right compare the local radius, our measure of local density, in the original space, with the local radius in the embedding (Methods). Points are colored by embryo time, and the $R^2$ value of the correlation is shown for each plot. The higher correlation in den-SNE than in t-SNE supports the increase in transcriptomic variability over time seen in the den-SNE plot. Analogous correlation plots based on neighborhood count, our other measure of local distance (Methods), are included in Supplementary Figure 8. **c.** To assess lineage-specific patterns, we consider the mean local radius within different time bins for six different cell-types in the original data (top), den-SNE (middle), and t-SNE (bottom). In the original data, the plots illustrate the temporal changes in the underlying local density, while for den-SNE and t-SNE, they illustrate the apparent changes in density based on the visualizations. Time intervals were given in the original dataset, and the $y$-axis shows the change in average local radius compared to the earliest time interval in log-scale. Note that the trajectories traced out by den-SNE follow those of the original dataset more closely than those from t-SNE, supporting the relative temporal homogeneity seen in the den-SNE plots of semi-clonal cell types (hypodermis, intestinal, and BWM) compared to the other cell types. **d.** We show the best linear fit of local radius v. embryo time for the three semi-clonal cell-types against all clonal cells, again aggregating cells within the time-intervals given in the publication, with error bars showing one standard deviation in local radius for all cells within the interval. The slope of the linear fit for clonal cells is significantly higher than those of the semi-clonal cells: 99% confidence intervals of the slope coefficients do not overlap. We show densMAP and UMAP plots restricted to the examples of semi-clonal (**e**) and clonal (**f**) cell types; circles are centered at the centroid of the points in each time bin, and radius indicates one standard deviation of these coordinates (both based on visualization coordinates). densMAP more accurately portrays that the variability of the semi-clonal cell types (**e**) is more homogeneous compared to the clonal cell types (**f**), whereas UMAP produces misleading visualizations.

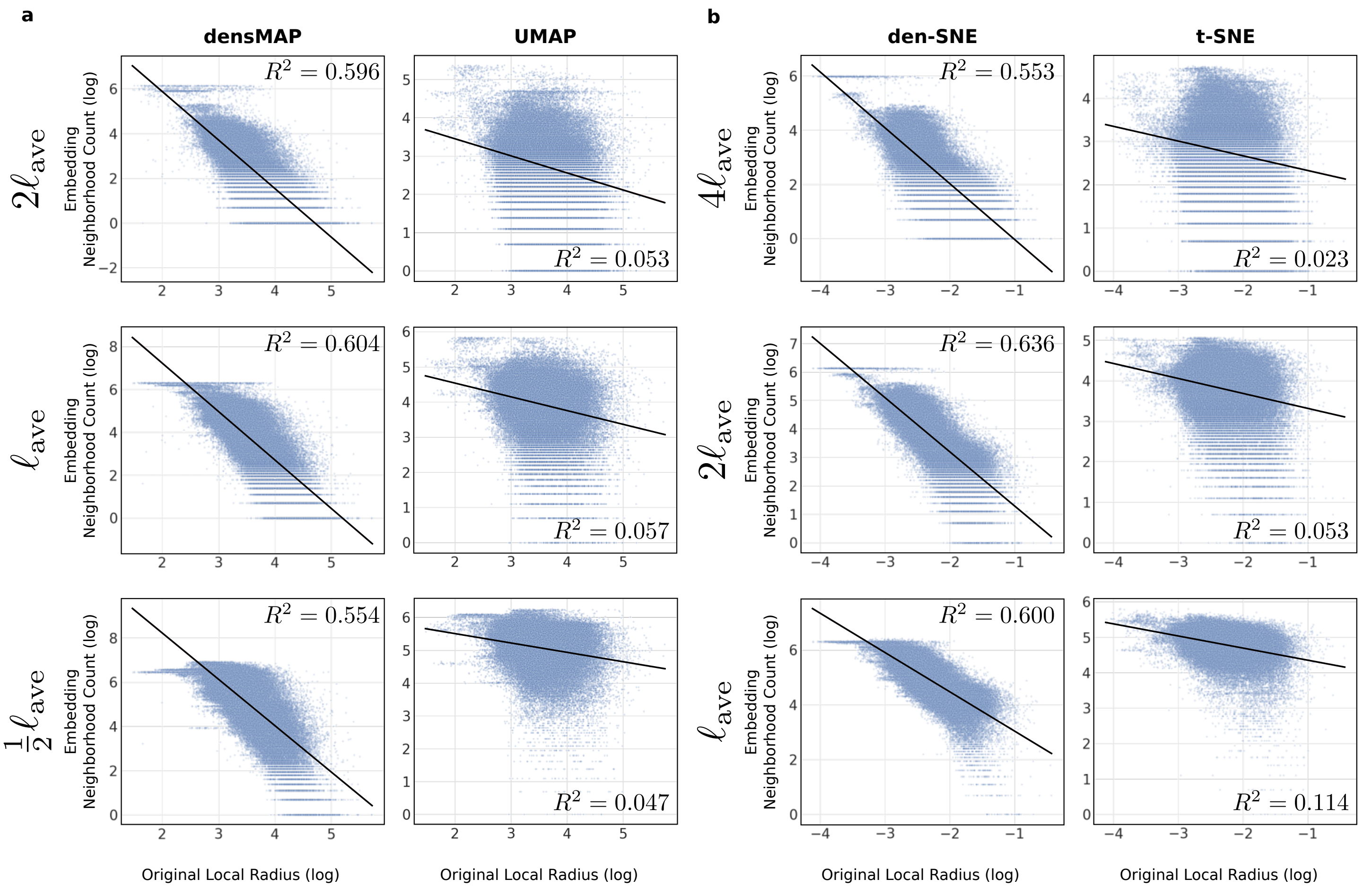


Supplementary Figure 8: **Density-preserving methods preserve density robustly at different scales on *C. elegans* embryo development data based on neighborhood count.** We compared the local radius of each point in the original *C. elegans* embryo development dataset to its neighborhood count in the visualizations (a measure of visual density; see Methods) for (**a**) densMAP and UMAP; and (**b**) den-SNE and t-SNE. We chose for each embedding, a length-scale $\ell_{\text{ave}}$ and multiples of that length-scale for which to compute the neighborhood counts (Methods): $\{\frac{1}{2}\ell_{\text{ave}}, \ell_{\text{ave}}, 2\ell_{\text{ave}}\}$ for densMAP and UMAP, and $\{\ell_{\text{ave}}, 2\ell_{\text{ave}}, 4\ell_{\text{ave}}\}$ for den-SNE and t-SNE. Since neighborhood count represents the density around a given point, a visualization that preserves density information will have higher neighborhood counts for points with smaller local radii in the original space. Note that this negative correlation is significantly stronger for our density-preserving tools than for t-SNE and UMAP, and this pattern holds across the different length-scales.

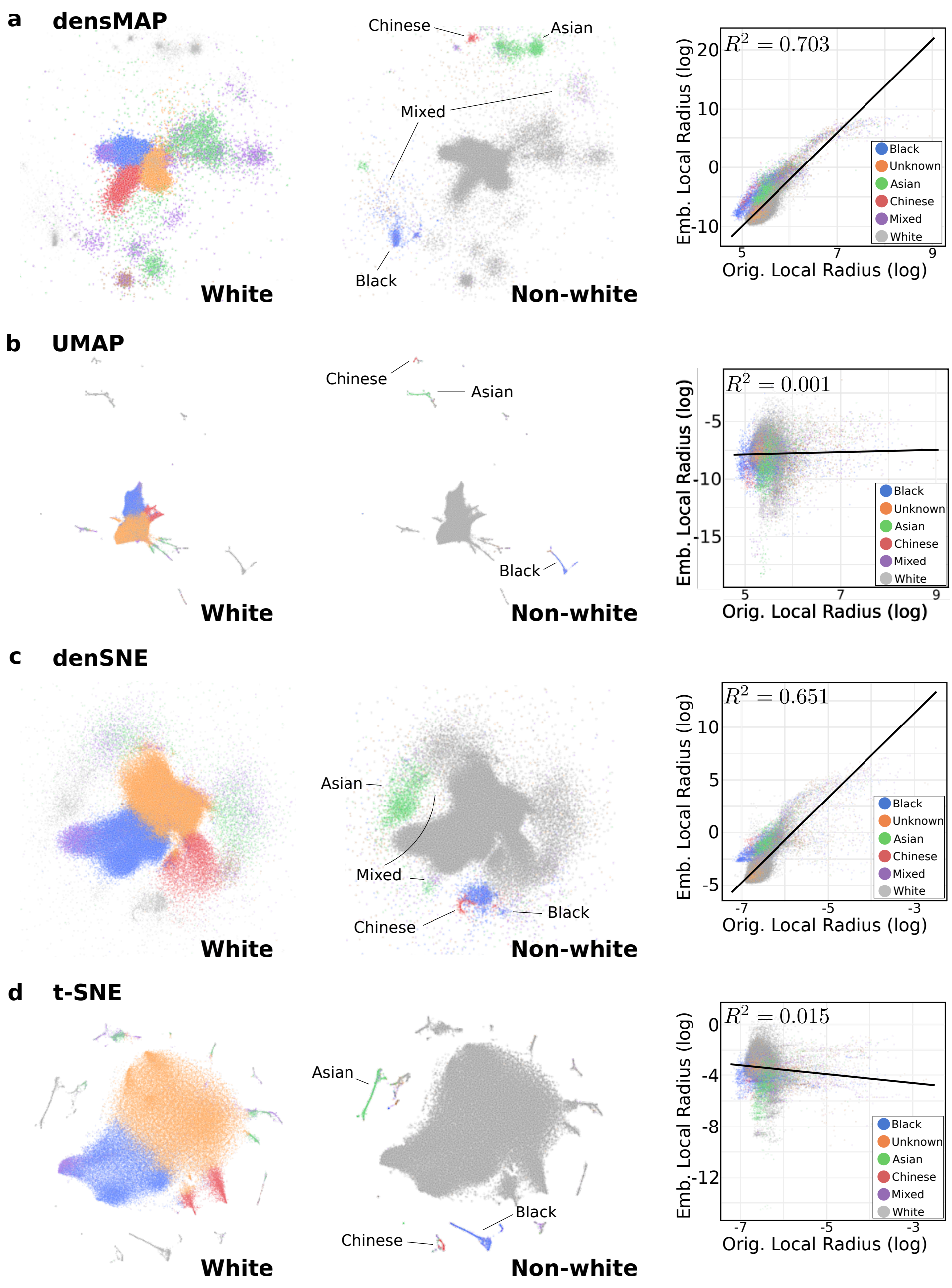


Supplementary Figure 9: **Density-preserving methods more accurately visualize diversity of small subpopulations in UKBB data.** We visualize the genotype profiles of 97,676 UKBB participants (a 20% subsample of the dataset) using (**a**) densMAP, (**b**) UMAP, (**c**) den-SNE and (**d**) t-SNE. For each, in the left plot, points corresponding to white people are colored by five computationally-identified subpopulations (Methods); in the middle plot, non-white people are colored according to their ethnicity; right shows correlation of local radius between the original dataset and the embedding, with points colored by ethnicity and $R^2$ reported. We show the analogous scatter plots using neighborhood count to measure local density in the visualization in Supplementary Figure 10. As 94% of the the people in the UKB dataset self-identified as white, the UMAP and t-SNE plots give overwhelming visual space to this group, hiding the genetic variability of the other ethnic groups. The density-preserving plots, however, clearly expand the clusters of non-white people as well as certain white subpopulations, more accurately conveying their genetic diversity.

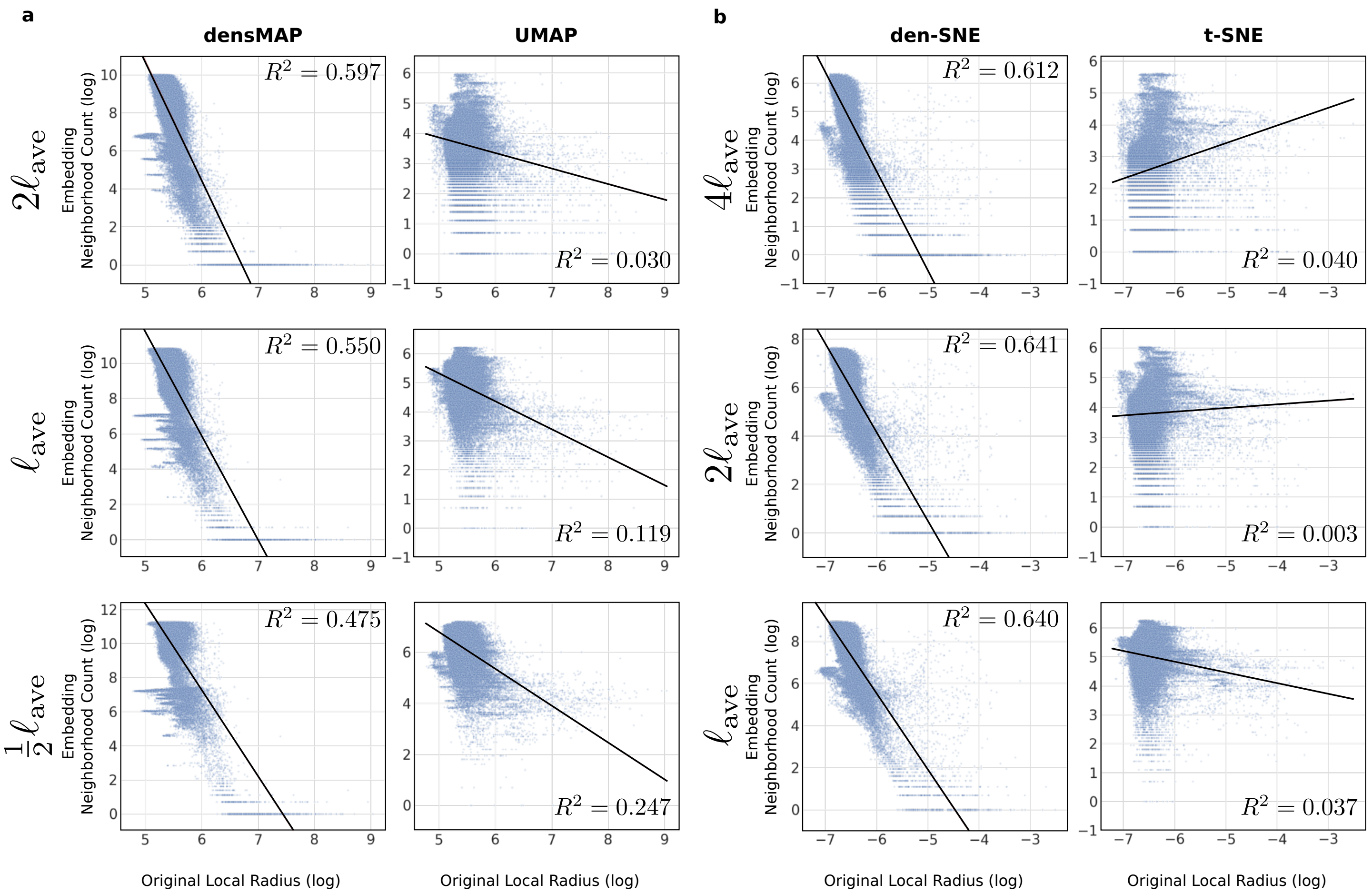


Supplementary Figure 10: **Density-preserving methods preserve density robustly at different scales on UKBB data based on neighborhood count.** We compared the local radius of each point in the original UKBB dataset (20% subsample) to its neighborhood count in the visualizations (a measure of visual density; see Methods) for (**a**) densMAP and UMAP; and (**b**) den-SNE and t-SNE. We chose for each embedding, a length-scale $\ell_{\text{ave}}$ and multiples of that length-scale for which to compute the neighborhood counts (Methods): $\{\frac{1}{2}\ell_{\text{ave}}, \ell_{\text{ave}}, 2\ell_{\text{ave}}\}$ for densMAP and UMAP, and $\{\ell_{\text{ave}}, 2\ell_{\text{ave}}, 4\ell_{\text{ave}}\}$ for den-SNE and t-SNE. Since neighborhood count represents the density around a given point, a visualization that preserves density information will have higher neighborhood counts for points with smaller local radii in the original space. Note that this negative correlation is significantly stronger for our density-preserving tools than for t-SNE and UMAP, and this pattern holds across the different length-scales.

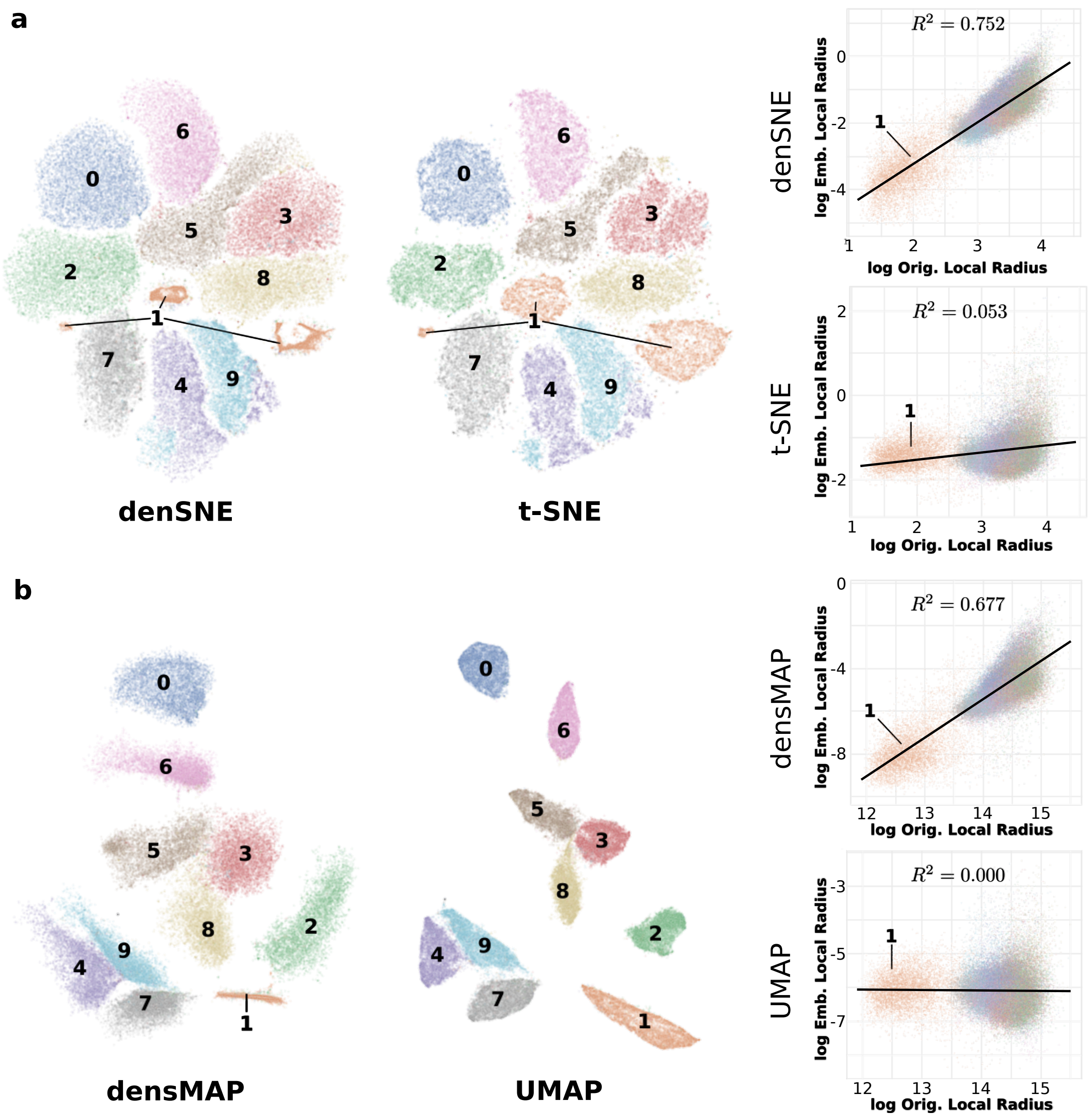


Supplementary Figure 11: **Density-preserving visualization of MNIST handwritten digit image dataset reveals the relative homogeneity of the digit 1.** We visualize the MNIST handwritten digits with (**a**) denSNE and t-SNE and (**b**) densMAP and UMAP, with points colored by digit. Note that the size of the cluster corresponding to the digit **1** shrinks under both density-preserving algorithms. Plots on the right show the correlation of the local radii between the original dataset and the embedding in each algorithm, with points colored by digit and the $R^2$ score reported. The higher $R^2$ for the density-preserving methods illustrates that the digit **1** indeed has higher density than the other digits. We show the analogous scatter plots using neighborhood count to measure local density in Supplementary Figure 12.

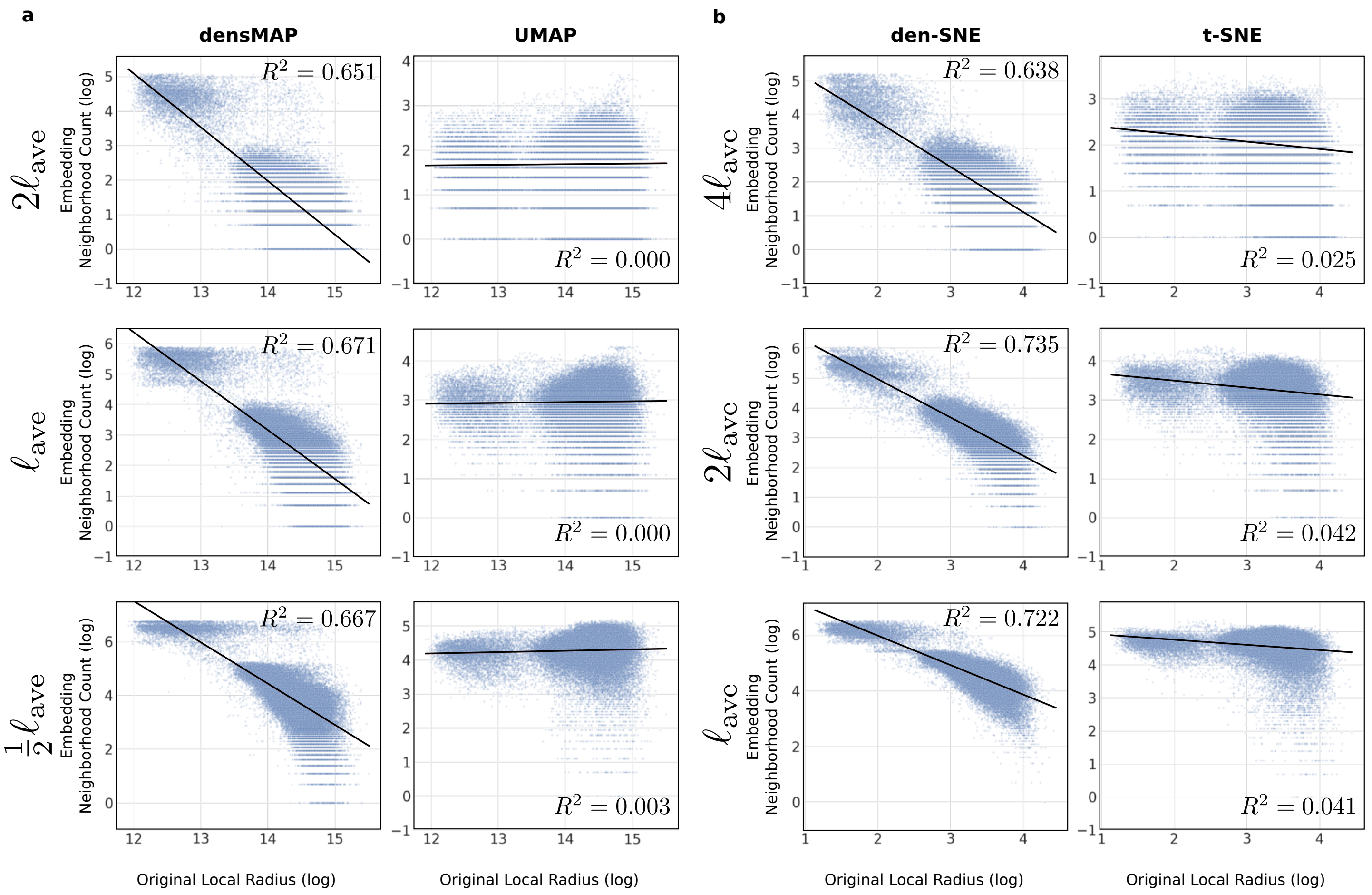


Supplementary Figure 12: **Density-preserving methods preserve density robustly at different scales on MNIST data based on neighborhood count.** We compared the local radius of each point in the original MNIST dataset to its neighborhood count in the visualizations (a measure of visual density; see Methods) for (**a**) densMAP and UMAP; and (**b**) den-SNE, and t-SNE. We chose for each embedding, a length-scale $\ell_{\text{ave}}$ and multiples of that length-scale for which to compute the neighborhood counts (Methods): $\{\frac{1}{2}\ell_{\text{ave}}, \ell_{\text{ave}}, 2\ell_{\text{ave}}\}$ for densMAP and UMAP, and $\{\ell_{\text{ave}}, 2\ell_{\text{ave}}, 4\ell_{\text{ave}}\}$ for den-SNE and t-SNE. Since neighborhood count represents the density around a given point, a visualization that preserves density information will have higher neighborhood counts for points with smaller local radii in the original space. Note that this negative correlation is significantly stronger for our density-preserving tools than for t-SNE and UMAP, and this pattern holds across the different length-scales.

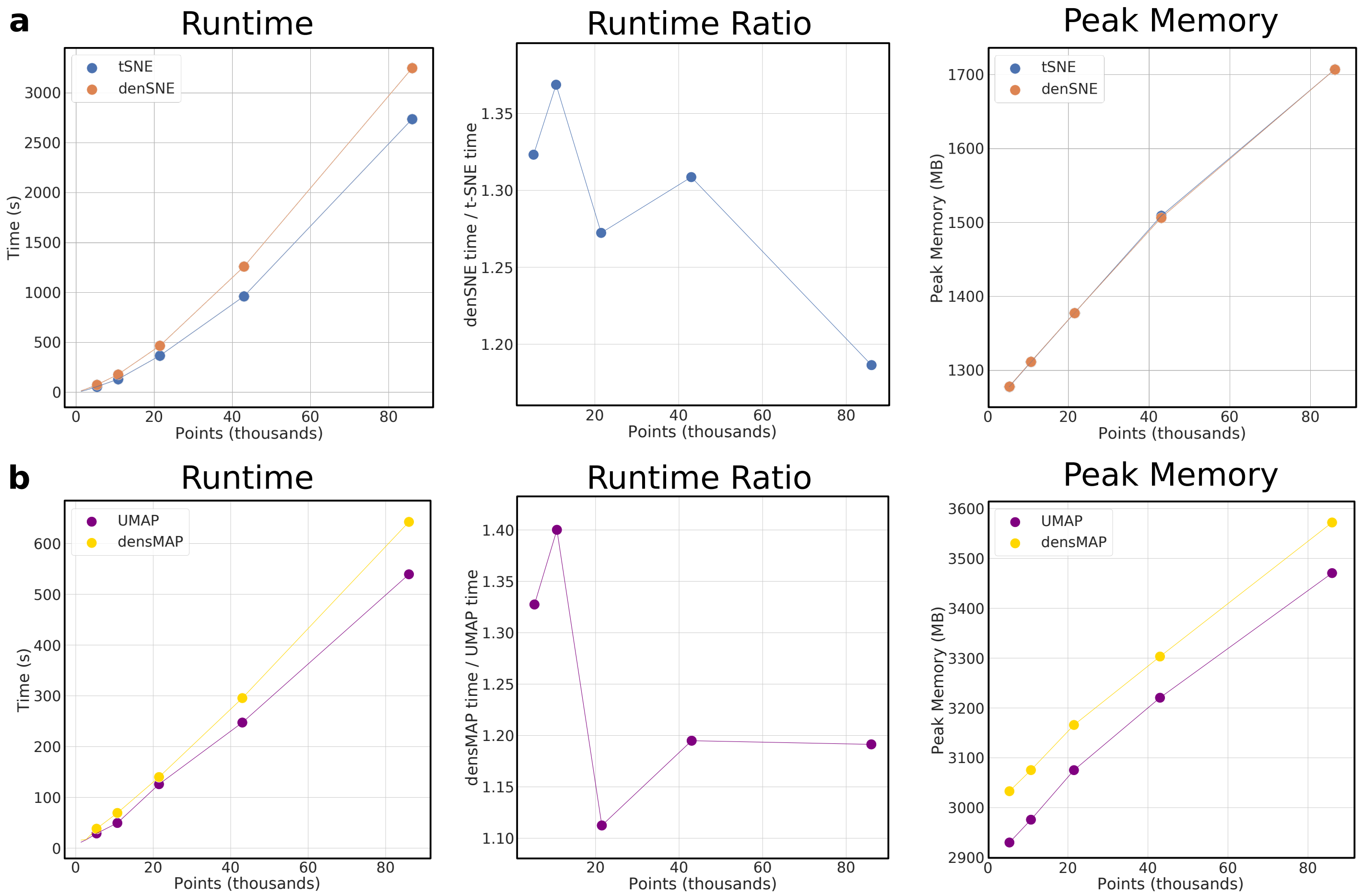


Supplementary Figure 13: **den-SNE and densMAP are nearly as efficient as t-SNE and UMAP in runtime and memory.** We compare (**a**) den-SNE and t-SNE and (**b**) densMAP and UMAP with respect to runtime and peak memory usage. For these tests, we exclude the time taken to compute the local radii of the final embedding, which is used only for evaluation and does not affect the embedding. Left plots running time in seconds at different data sizes (achieved by subsampling the *C. elegans* dataset; Methods); middle shows the ratio of the density-preserving algorithm's runtime to that of the original method; right shows peak memory usage over different data sizes. Although density-preserving methods take longer, the overhead is small (around 20% additional runtime at 86k cells for both methods). For den-SNE, the relative overhead of density-preservation becomes smaller for larger datasets. Both densMAP and UMAP obtain fast runtimes for large datasets, taking less than ∼10 minutes for all our datasets. Peak memory usage is the same between t-SNE and den-SNE, and differs by a small constant between UMAP and densMAP.

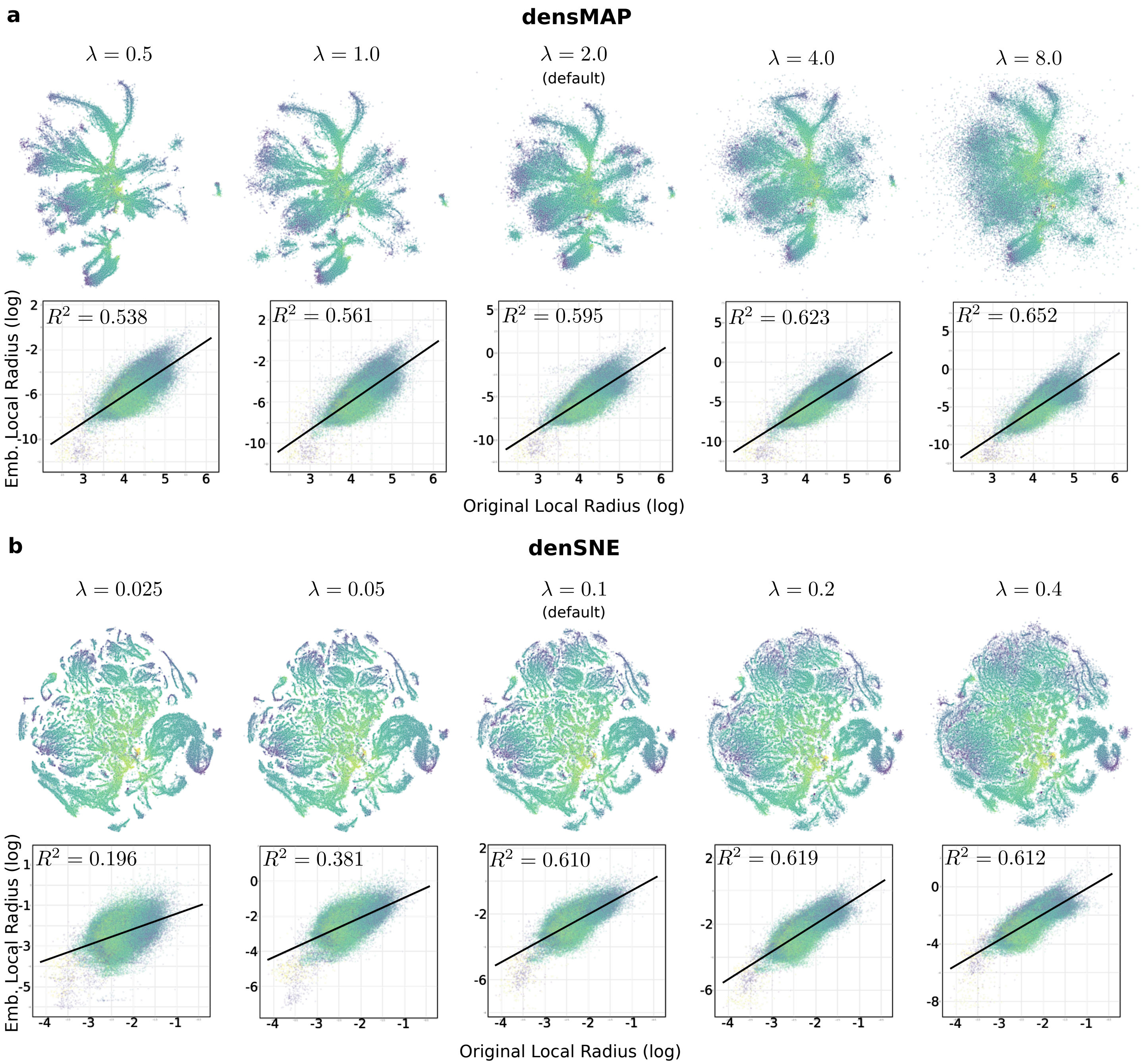


Supplementary Figure 14: **Varying the density weight parameter in densMAP and den-SNE controls the trade-off between density preservation and cluster separation.** We demonstrate on the *C. elegans* dataset the effects of varying the weight $\lambda$ of the density-preservation term in the objective function of (**a**) densMAP and (**b**) den-SNE. As $\lambda$ increases, so does the correlation between the log local radii in the original data and the embedding, but the clusters begin to fade into each other, likely due to lack of space in the visualization. As $\lambda$ decreases, the correlation becomes worse and the embeddings become closer to those of t-SNE and UMAP. Based on our results from a wide range of datasets, we recommend the default values of $\lambda = 0.1$ for den-SNE and $\lambda = 2.0$ for densMAP.

Gene Expression in Tumor vs. Blood (T Cells CD8)

| | | | | | Permutation Test $p$-value (Bonferroni-corrected) | | |
|---|---|---|---|---|---|---|---|
| Rank | Gene | $\Delta$(Variance) | $\Delta$(Dispersion) | $\Delta$(Mean) | Dispersion | Mean | Variance |
| 1 | DUSP4 | 1.034 | 0.492 | 0.919 | **<2E-4** | **<2E-4** | **<2E-4** |
| 2 | RGS1 | 0.995 | 0.009 | 0.926 | 1.0 | **<2E-4** | **<2E-4** |
| 3 | RGCC | 0.922 | 0.045 | 1.024 | 1.0 | **<2E-4** | **<2E-4** |
| 4 | TNFAIP3 | 0.816 | -0.173 | 0.970 | 1.0 | **<2E-4** | **<2E-4** |
| 5 | NR4A2 | 0.800 | 0.023 | 0.933 | 1.0 | **<2E-4** | **<2E-4** |
| 6 | ZFP36 | 0.792 | 0.128 | 0.864 | 0.1346 | **<2E-4** | **<2E-4** |
| 7 | CCL4 | 0.783 | 0.619 | 0.358 | **<2E-4** | **<2E-4** | **<2E-4** |
| 8 | CREM | 0.764 | -0.201 | 0.811 | 1.0 | **<2E-4** | **<2E-4** |
| 9 | JUNB | 0.742 | 0.116 | 0.736 | 0.676 | **<2E-4** | **<2E-4** |
| 10 | HSP90AA1 | 0.701 | 0.252 | 0.605 | **<2E-4** | **<2E-4** | **<2E-4** |
| 11 | DNAJB1 | 0.693 | 0.055 | 0.616 | 1.0 | **<2E-4** | **<2E-4** |
| 12 | FOSB | 0.678 | 0.594 | 0.565 | **<2E-4** | **<2E-4** | **<2E-4** |
| 13 | RPS26 | 0.675 | 0.199 | 0.516 | **<2E-4** | **<2E-4** | **<2E-4** |
| 14 | ZNF331 | 0.670 | 0.456 | 0.641 | **<2E-4** | **<2E-4** | **<2E-4** |
| 15 | JUND | 0.656 | -0.027 | 0.906 | 1.0 | **<2E-4** | **<2E-4** |
| 16 | FTH1 | 0.588 | 0.141 | 0.618 | 0.0516 | **<2E-4** | **<2E-4** |
| 17 | FOSL2 | 0.585 | 0.008 | 0.599 | 1.0 | **<2E-4** | **<2E-4** |
| 18 | IGKC | 0.579 | 1.135 | 0.335 | **<2E-4** | **<2E-4** | **<2E-4** |
| 19 | YPEL5 | 0.569 | 0.193 | 0.575 | **0.005** | **<2E-4** | **<2E-4** |
| 20 | TSC22D3 | 0.560 | 0.231 | 0.476 | **<2E-4** | **<2E-4** | **<2E-4** |

Supplementary Table 1: **Genes with largest difference in variance between blood and tumor CD8 T cells.** The columns $\Delta$ {Dispersion, Mean, Variance} show the changes in the corresponding statistic for each listed gene in tumor relative to blood. We performed permutation tests (last three columns) to calculate the significance of the change in mean, dispersion, and variance, and those $p$-values which are significant after Bonferroni correction ($p < 0.01$) are shown in boldface.

Gene Expression in Tumor vs. Blood (T Cells CD4 Memory Resting)

| | | | | | Permutation Test $p$-value (Bonferroni-corrected) | | |
|---|---|---|---|---|---|---|---|
| Rank | Gene | Δ(Variance) | Δ(Dispersion) | Δ(Mean) | Dispersion | Mean | Variance |
| 1 | RGS1 | 1.013 | -0.009 | 1.069 | 1.0 | **<2E-4** | **<2E-4** |
| 2 | DUSP4 | 0.952 | 0.295 | 0.753 | **<2E-4** | **<2E-4** | **<2E-4** |
| 3 | TNFAIP3 | 0.818 | -0.205 | 1.099 | 1.0 | **<2E-4** | **<2E-4** |
| 4 | JUNB | 0.810 | -0.070 | 1.110 | 1.0 | **<2E-4** | **<2E-4** |
| 5 | ZFP36 | 0.809 | 0.039 | 1.045 | 1.0 | **<2E-4** | **<2E-4** |
| 6 | RGCC | 0.779 | 0.101 | 0.922 | 0.0124 | **<2E-4** | **<2E-4** |
| 7 | CREM | 0.762 | 0.111 | 0.937 | 0.1886 | **<2E-4** | **<2E-4** |
| 8 | HSP90AA1 | 0.730 | 0.185 | 0.795 | **<2E-4** | **<2E-4** | **<2E-4** |
| 9 | NR4A2 | 0.669 | 0.216 | 0.737 | **<2E-4** | **<2E-4** | **<2E-4** |
| 10 | HSPA1A | 0.664 | 0.911 | 0.367 | **<2E-4** | **<2E-4** | **<2E-4** |
| 11 | LMNA | 0.656 | 0.455 | 0.556 | **<2E-4** | **<2E-4** | **<2E-4** |
| 12 | DNAJB1 | 0.610 | 0.201 | 0.566 | **<2E-4** | **<2E-4** | **<2E-4** |
| 13 | JUND | 0.598 | -0.049 | 0.979 | 1.0 | **<2E-4** | **<2E-4** |
| 14 | FTH1 | 0.583 | 0.041 | 1.067 | 1.0 | **<2E-4** | **<2E-4** |
| 15 | SLC2A3 | 0.561 | 0.091 | 0.631 | 0.0196 | **<2E-4** | **<2E-4** |
| 16 | ZNF331 | 0.560 | 0.029 | 0.586 | 1.0 | **<2E-4** | **<2E-4** |
| 17 | YPEL5 | 0.557 | 0.052 | 0.657 | 1.0 | **<2E-4** | **<2E-4** |
| 18 | HSPH1 | 0.553 | 0.297 | 0.493 | **<2E-4** | **<2E-4** | **<2E-4** |
| 19 | TSC22D3 | 0.549 | 0.213 | 0.534 | **<2E-4** | **<2E-4** | **<2E-4** |
| 20 | FOSL2 | 0.541 | 0.211 | 0.612 | **<2E-4** | **<2E-4** | **<2E-4** |

Supplementary Table 2: **Genes with largest difference in variance between blood and tumor memory resting CD4 T cells.** The columns Δ {Dispersion, Mean, Variance} show the changes in the corresponding statistic for each listed gene in tumor relative to blood. We performed permutation tests (last three columns) to calculate the significance of the change in mean, dispersion, and variance, and those $p$-values which are significant after Bonferroni correction ($p < 0.01$) are shown in boldface.

Gene Expression in Tumor vs. Blood (T Cells Naïve CD4)

| | | | | | Permutation Test $p$-value (Bonferroni-corrected) | | |
|---|---|---|---|---|---|---|---|
| Rank | Gene | Δ(Variance) | Δ(Dispersion) | Δ(Mean) | Dispersion | Mean | Variance |
| 1 | MT-ATP6 | 1.143 | 0.416 | -0.331 | **<2E-4** | **0.0078** | **<2E-4** |
| 2 | RPS26 | 1.114 | 0.047 | 0.97 | 1.0 | **<2E-4** | **<2E-4** |
| 3 | FTH1 | 1.052 | 0.432 | 0.787 | **<2E-4** | **<2E-4** | **<2E-4** |
| 4 | RPS14 | 0.899 | 0.404 | -0.048 | **<2E-4** | 1.0 | **<2E-4** |
| 5 | CREM | 0.888 | 0.628 | 0.66 | 1.0 | **<2E-4** | **<2E-4** |
| 6 | MT-RNR2 | 0.887 | 0.253 | 0.82 | 0.0128 | **<2E-4** | **<2E-4** |
| 7 | RPLP1 | 0.833 | 0.372 | 0.007 | **<2E-4** | 1.0 | **<2E-4** |
| 8 | RPS27 | 0.832 | 0.299 | -0.097 | **0.0046** | 1.0 | **<2E-4** |
| 9 | RPS3 | 0.828 | 0.457 | -0.198 | **<2E-4** | 0.497 | **<2E-4** |
| 10 | MTRNR2L12 | 0.82 | 0.274 | 0.797 | 0.0194 | <2E-4 | **<2E-4** |
| 11 | CXCR4 | 0.814 | 0.366 | 0.661 | 0.166 | <2E-4 | **<2E-4** |
| 12 | SRGN | 0.81 | 0.377 | 0.678 | 0.129 | <2E-4 | **<2E-4** |
| 13 | MT-ND1 | 0.762 | 0.315 | -0.155 | **0.0042** | 1.0 | **<2E-4** |
| 14 | RPL34 | 0.754 | 0.361 | -0.426 | **<2E-4** | <2E-4 | **<2E-4** |
| 15 | TTC19 | 0.753 | 0.414 | 0.026 | **<2E-4** | 1.0 | **<2E-4** |
| 16 | PABPC1 | 0.744 | 0.481 | -0.099 | **<2E-4** | 1.0 | **<2E-4** |
| 17 | MT-CO2 | 0.733 | 0.307 | 0.407 | **0.0084** | <2E-4 | **<2E-4** |
| 18 | RPL11 | 0.721 | 0.469 | -0.601 | **<2E-4** | <2E-4 | **<2E-4** |
| 19 | MT-CO1 | 0.719 | 0.391 | -0.118 | **<2E-4** | 1.0 | **<2E-4** |
| 20 | RPL27A | 0.716 | 0.387 | -0.289 | **<2E-4** | 0.0138 | **<2E-4** |

Supplementary Table 3: **Genes with largest difference in variance between blood and tumor CD4 naïve T cells.** The columns Δ {Dispersion, Mean, Variance} show the changes in the corresponding statistic for each listed gene in tumor relative to blood. We performed permutation tests (last three columns) to calculate the significance of the change in mean, dispersion, and variance, and those $p$-values which are significant after Bonferroni correction ($p < 0.01$) are shown in boldface.

Gene Expression in Tumor vs. Blood (B Cells Memory)

| | | | | | Permutation Test $p$-value (Bonferroni-corrected) | | |
|---|---|---|---|---|---|---|---|
| Rank | Gene | Δ(Variance) | Δ(Dispersion) | Δ(Mean) | Dispersion | Mean | Variance |
| 1 | RGS1 | 0.882 | 0.644 | 0.764 | **<2E-4** | **<2E-4** | **<2E-4** |
| 2 | NR4A2 | 0.839 | N/A | 1.022 | N/A | **<2E-4** | **<2E-4** |
| 3 | JUNB | 0.742 | -0.074 | 0.896 | 1.0 | **<2E-4** | **<2E-4** |
| 4 | CD83 | 0.674 | 0.230 | 0.771 | 0.224 | **<2E-4** | **<2E-4** |
| 5 | RGS2 | 0.647 | -0.177 | 0.711 | 1.0 | **<2E-4** | **<2E-4** |
| 6 | HSP90AA1 | 0.644 | 0.227 | 0.682 | 0.3564 | **<2E-4** | **<2E-4** |
| 7 | FOSB | 0.635 | N/A | 0.667 | N/A | **<2E-4** | **<2E-4** |
| 8 | JUND | 0.598 | 0.151 | 0.893 | 1.0 | **<2E-4** | **<2E-4** |
| 9 | GPR183 | 0.586 | 0.366 | 0.539 | 0.0246 | **<2E-4** | **<2E-4** |
| 10 | HSPH1 | 0.565 | 0.244 | 0.555 | 0.3082 | **<2E-4** | **<2E-4** |
| 11 | ZNF331 | 0.560 | 0.489 | 0.557 | **<2E-4** | **<2E-4** | **<2E-4** |
| 12 | SRGN | 0.541 | 0.047 | 0.619 | 1.0 | **<2E-4** | **<2E-4** |
| 13 | DUSP4 | 0.529 | N/A | 0.404 | N/A | **<2E-4** | **<2E-4** |
| 14 | TSC22D3 | 0.515 | -0.158 | 0.794 | 1.0 | **<2E-4** | **<2E-4** |
| 15 | NR4A3 | 0.512 | N/A | 0.454 | N/A | **<2E-4** | **<2E-4** |
| 16 | HSPA1A | 0.507 | 0.957 | 0.350 | **<2E-4** | **<2E-4** | **<2E-4** |
| 17 | SLC2A3 | 0.504 | -0.317 | 0.569 | 1.0 | **<2E-4** | **<2E-4** |
| 18 | LY9 | 0.503 | 0.098 | 0.569 | 1.0 | **<2E-4** | **<2E-4** |
| 19 | HSP90AB1 | 0.499 | 0.122 | 0.573 | 1.0 | **<2E-4** | **<2E-4** |
| 20 | MCL1 | 0.496 | 0.215 | 0.492 | 0.3442 | **<2E-4** | **<2E-4** |

Supplementary Table 4: **Genes with largest difference in variance between blood and tumor memory B cells.** The columns Δ {Dispersion, Mean, Variance} show the changes in the corresponding statistic for each listed gene in tumor relative to blood. N/A values in the Δ(Dispersion) and Dispersion $p$-value columns indicate that the gene had zero mean-expression in blood, and so dispersion is undefined. We performed permutation tests (last three columns) to calculate the significance of the change in mean, dispersion, and variance, and those $p$-values which are significant after Bonferroni correction ($p < 0.01$) are shown in boldface.

Gene Expression in Tumor vs. Blood (B Cells Naïve)

| | | | | | Permutation Test $p$-value (Bonferroni-corrected) | | |
|---|---|---|---|---|---|---|---|
| Rank | Gene | Δ(Variance) | Δ(Dispersion) | Δ(Mean) | Dispersion | Mean | Variance |
| 1 | JUNB | 0.967 | 0.264 | 1.094 | 0.322 | **<2E-4** | **<2E-4** |
| 2 | CD83 | 0.852 | -0.013 | 0.937 | 1.0 | **<2E-4** | **<2E-4** |
| 3 | RPS27 | 0.827 | 0.329 | -0.239 | **<2E-4** | 0.6354 | **<2E-4** |
| 4 | NR4A2 | 0.811 | N/A | 0.876 | N/A | **<2E-4** | **<2E-4** |
| 5 | JUND | 0.681 | -0.087 | 1.025 | 1.0 | **<2E-4** | **<2E-4** |
| 6 | FOS | 0.660 | N/A | 0.515 | N/A | **<2E-4** | **<2E-4** |
| 7 | FOSB | 0.652 | 0.320 | 0.573 | 0.4072 | **<2E-4** | **<2E-4** |
| 8 | RPL13A | 0.634 | 0.287 | -0.362 | **<2E-4** | **0.008** | **<2E-4** |
| 9 | DUSP1 | 0.629 | -0.188 | 0.632 | 1.0 | **<2E-4** | **<2E-4** |
| 10 | IGHM | 0.625 | 0.506 | -0.928 | **<2E-4** | **<2E-4** | **<2E-4** |
| 11 | RGS1 | 0.620 | N/A | 0.416 | N/A | **<2E-4** | **<2E-4** |
| 12 | ZNF331 | 0.616 | -0.140 | 0.584 | 1.0 | **<2E-4** | **<2E-4** |
| 13 | TSC22D3 | 0.609 | 0.048 | 0.715 | 1.0 | **<2E-4** | **<2E-4** |
| 14 | JUN | 0.590 | -0.875 | 0.498 | 1.0 | **<2E-4** | **<2E-4** |
| 15 | HERPUD1 | 0.581 | -0.033 | 0.583 | 1.0 | **<2E-4** | **<2E-4** |
| 16 | RPS15A | 0.578 | 0.394 | -0.490 | **<2E-4** | **<2E-4** | **<2E-4** |
| 17 | LY9 | 0.575 | 0.474 | 0.548 | **<2E-4** | **<2E-4** | **<2E-4** |
| 18 | RPS12 | 0.567 | 0.292 | -0.434 | **0.0052** | **<2E-4** | **<2E-4** |
| 19 | SLC2A3 | 0.552 | 0.070 | 0.560 | 1.0 | **<2E-4** | **<2E-4** |
| 20 | RPS19 | 0.549 | 0.374 | -0.403 | **<2E-4** | **<2E-4** | **<2E-4** |

Supplementary Table 5: **Genes with largest difference variance between blood and tumor naïve B cells.** The columns Δ {Dispersion, Mean, Variance} show the changes in the corresponding statistic for each listed gene in tumor relative to blood. N/A values in the Δ(Dispersion) and Dispersion $p$-value columns indicate that the gene had zero mean-expression in blood, and so dispersion is undefined. We performed permutation tests (last three columns) to calculate the significance of the change in mean, dispersion, and variance, and those $p$-values which are significant after Bonferroni correction ($p < 0.01$) are shown in boldface.

T Cells CD8 (Validation)

| | | | | | Permutation Test $p$-value (Bonferroni-corrected) | | |
|---|---|---|---|---|---|---|---|
| Rank | Gene | Δ(Variance) | Δ(Dispersion) | Δ(Mean) | Dispersion | Mean | Variance |
| 1 | RGS1 | 8.099 | -0.782 | 4.155 | 1.0 | **<2E-4** | **<2E-4** |
| 2 | DUSP4 | 4.170 | 0.522 | 1.238 | 1.0 | **<2E-4** | **<2E-4** |
| 3 | FOSB | 4.143 | 0.126 | 1.182 | 1.0 | **<2E-4** | **<2E-4** |
| 4 | RGCC | 3.299 | -0.697 | 0.941 | 1.0 | **<2E-4** | **<2E-4** |
| 5 | NR4A2 | 2.788 | 0.496 | 0.631 | 0.1368 | **<2E-4** | **<2E-4** |
| 6 | TSC22D3 | 2.517 | 0.585 | -0.766 | **<2E-4** | **<2E-4** | **<2E-4** |
| 7 | CREM | 2.475 | -0.437 | 0.765 | 1.0 | **<2E-4** | **<2E-4** |
| 8 | TNFAIP3 | 1.747 | -1.071 | 1.789 | 1.0 | **<2E-4** | **<2E-4** |
| 9 | JUND | 0.785 | -0.334 | 0.335 | 1.0 | **<2E-4** | **<2E-4** |
| 10 | ZNF331 | 0.433 | -0.115 | 0.127 | 1.0 | 0.687 | 0.8736 |
| 11 | DNAJB1 | 0.397 | 0.077 | 0.058 | 1.0 | 1.0 | 0.6108 |
| 12 | JUNB | 0.350 | -0.666 | 0.805 | 1.0 | **<2E-4** | 1.0 |
| 13 | YPEL5 | 0.249 | -0.586 | 0.504 | 1.0 | **<2E-4** | 1.0 |
| 14 | FOSL2 | 0.007 | -0.161 | 0.018 | 1.0 | 1.0 | 1.0 |
| 15 | FTH1 | -0.051 | -0.011 | 0.086 | 1.0 | 1.0 | 1.0 |
| 16 | ZFP36 | -0.286 | -0.287 | 0.687 | 1.0 | **<2E-4** | 1.0 |
| 17 | HSP90AA1 | -0.435 | -0.191 | 0.395 | 1.0 | **<2E-4** | 1.0 |
| 18 | RPS26 | -0.452 | -0.330 | -0.045 | 1.0 | 1.0 | 1.0 |
| 19 | CCL4 | -2.924 | -1.309 | 2.323 | 1.0 | **<2E-4** | 1.0 |

Supplementary Table 6: **Validation of top differentially variable genes between tumor and blood in CD8 T cells on a secondary dataset.** We repeat the tests for difference in mean, variance, and dispersion index of 19 of the top 20 genes from Supplementary Table 1 for CD8 T cells in blood and lung cancer on a secondary dataset from Guo et al. (2018) (IGKC was not found in the dataset). Other cell types we analyzed did not have a close match in this dataset and were omitted from the analysis. The columns Δ {Dispersion, Mean, Variance} show the changes in the corresponding statistic for each listed gene in tumor relative to blood. We performed permutation tests (last three columns) to calculate the significance of the change in dispersion, mean, and variance, and those $p$-values which are significant after Bonferroni correction ($p < 0.01$) are shown in boldface. We see that 9 of the 19 genes are significant in differential variance in this dataset as well and TSC22D3 is also significantly overdispersed in tumor in this dataset.

**T Cells CD8 (GO Enrichment)**

| GO Biological Process Complete | Bgnd | Count | Expected | +/- | Fold Enrich | P-value |
|---|---|---|---|---|---|---|
| negative regulation of interleukin-2 production (GO:0032703) | 24 | 3 | 0.03 | + | >100 | 2.55E-02 |
| response to bacterium (GO:0009617) | 691 | 7 | 0.72 | + | 9.68 | 4.06E-02 |
| negative regulation of transcription by RNA polymerase II (GO:0000122) | 886 | 8 | 0.93 | + | 8.63 | 1.63E-02 |
| negative regulation of RNA metabolic process (GO:0051253) | 1371 | 9 | 1.43 | + | 6.27 | 4.13E-02 |
| regulation of transcription by RNA polymerase II (GO:0006357) | 2255 | 11 | 2.36 | + | 4.66 | 3.96E-02 |
| regulation of nucleobase-containing compound metabolic process (GO:0019219) | 4019 | 15 | 4.21 | + | 3.57 | 4.81E-03 |
| regulation of RNA metabolic process (GO:0051252) | 3768 | 14 | 3.94 | + | 3.55 | 1.88E-02 |
| regulation of cellular macromolecule biosynthetic process (GO:2000112) | 3894 | 14 | 4.08 | + | 3.44 | 2.84E-02 |
| regulation of macromolecule biosynthetic process (GO:0010556) | 4033 | 14 | 4.22 | + | 3.32 | 4.38E-02 |
| negative regulation of cellular process (GO:0048523) | 4768 | 15 | 4.99 | + | 3.01 | 4.78E-02 |
| negative regulation of biological process (GO:0048519) | 5355 | 16 | 5.6 | + | 2.86 | 2.91E-02 |
| regulation of nitrogen compound metabolic process (GO:0051171) | 5821 | 17 | 6.09 | + | 2.79 | 1.11E-02 |
| regulation of primary metabolic process (GO:0080090) | 6004 | 17 | 6.28 | + | 2.71 | 1.79E-02 |
| regulation of macromolecule metabolic process (GO:0060255) | 6140 | 17 | 6.43 | + | 2.65 | 2.53E-02 |
| regulation of cellular metabolic process (GO:0031323) | 6212 | 17 | 6.5 | + | 2.62 | 3.03E-02 |
| **Analysis Type:** | PANTHER Overrepresentation Test (Released 20190711) | | | | | |
| **Annotation Version and Release Date:** | GO Ontology database Released 2020-02-21 | | | | | |

Supplementary Table 7: **GO enrichment analysis of differentially variable genes between tumor and blood for CD8 T cells in the lung cancer dataset.** We obtained the gene ontology (GO) terms significantly enriched in the top twenty genes ranked by increase in variance in tumor v. blood for CD8 T cells (shown in Supplementary Table 1). The analysis was performed using the web service available at: http://geneontology.org/. Significance is calculated using Fisher's exact test, and *p*-values are Bonferroni corrected. Bgnd: Background count

**T Cells CD4 Memory Resting (GO Enrichment)**

| GO Biological Process Complete | Bgnd | Count | Expected | +/- | Fold Enrich | P-value |
|---|---|---|---|---|---|---|
| negative regulation of interleukin-2 production (GO:0032703) | 24 | 3 | 0.03 | + | >100 | 2.55E-02 |
| chaperone cofactor-dependent protein refolding (GO:0051085) | 30 | 3 | 0.03 | + | 95.56 | 4.74E-02 |
| regulation of cellular response to heat (GO:1900034) | 79 | 4 | 0.08 | + | 48.38 | 1.38E-02 |
| response to unfolded protein (GO:0006986) | 166 | 5 | 0.17 | + | 28.78 | 6.92E-03 |
| response to topologically incorrect protein (GO:0035966) | 188 | 5 | 0.2 | + | 25.41 | 1.26E-02 |
| negative regulation of transcription by RNA polymerase II (GO:0000122) | 886 | 8 | 0.93 | + | 8.63 | 1.63E-02 |
| cellular response to stress (GO:0033554) | 1744 | 11 | 1.83 | + | 6.03 | 3.05E-03 |
| regulation of transcription by RNA polymerase II (GO:0006357) | 2255 | 11 | 2.36 | + | 4.66 | 3.96E-02 |
| positive regulation of macromolecule metabolic process (GO:0010604) | 3388 | 13 | 3.55 | + | 3.67 | 4.34E-02 |
| regulation of nucleobase-containing compound metabolic process (GO:0019219) | 4019 | 15 | 4.21 | + | 3.57 | 4.81E-03 |
| regulation of macromolecule biosynthetic process (GO:0010556) | 4033 | 15 | 4.22 | + | 3.55 | 5.04E-03 |
| regulation of cellular macromolecule biosynthetic process (GO:2000112) | 3894 | 14 | 4.08 | + | 3.44 | 2.84E-02 |
| regulation of biosynthetic process (GO:0009889) | 4258 | 15 | 4.46 | + | 3.37 | 1.05E-02 |
| negative regulation of cellular process (GO:0048523) | 4768 | 15 | 4.99 | + | 3.01 | 4.78E-02 |
| regulation of macromolecule metabolic process (GO:0060255) | 6140 | 17 | 6.43 | + | 2.65 | 2.53E-02 |
| **Analysis Type:** | PANTHER Overrepresentation Test (Released 20190711) | | | | | |
| **Annotation Version and Release Date:** | GO Ontology database Released 2020-02-21 | | | | | |

Supplementary Table 8: **GO enrichment analysis of differentially variable genes between tumor and blood for CD4 memory resting T cells in the lung cancer dataset.** We obtained the gene ontology (GO) terms significantly enriched in the top twenty genes ranked by increase in variance in tumor v. blood for CD4 memory resting T cells (shown in Supplementary Table 2). The analysis was performed using the web service available at: http://geneontology.org/. Significance is calculated using Fisher's exact test, and *p*-values are Bonferroni corrected. Bgnd: Background count

**CD4 T Cells Naïve**

| GO biological process complete | Bgnd | Count | Expected | +/- | Fold Enrich | P-value |
|---|---|---|---|---|---|---|
| SRP-dependent cotranslational protein targeting to membrane (GO:0006614) | 96 | 8 | 0.1 | + | 83.55 | 4.25E-10 |
| cotranslational protein targeting to membrane (GO:0006613) | 101 | 8 | 0.1 | + | 79.41 | 6.26E-10 |
| nuclear-transcribed mRNA catabolic process, nonsense-mediated decay (GO:0000184) | 120 | 9 | 0.12 | + | 75.19 | 2.01E-11 |
| protein targeting to ER (GO:0045047) | 110 | 8 | 0.11 | + | 72.92 | 1.2E-09 |
| establishment of protein localization to endoplasmic reticulum (GO:0072599) | 114 | 8 | 0.11 | + | 70.36 | 1.58E-09 |
| viral transcription (GO:0019083) | 115 | 8 | 0.11 | + | 69.75 | 1.68E-09 |
| translational initiation (GO:0006413) | 142 | 9 | 0.14 | + | 63.55 | 8.55E-11 |
| viral gene expression (GO:0019080) | 132 | 8 | 0.13 | + | 60.76 | 4.84E-09 |
| protein localization to endoplasmic reticulum (GO:0070972) | 138 | 8 | 0.14 | + | 58.12 | 6.8E-09 |
| nuclear-transcribed mRNA catabolic process (GO:0000956) | 194 | 9 | 0.19 | + | 46.51 | 1.27E-09 |
| protein targeting to membrane (GO:0006612) | 177 | 8 | 0.18 | + | 45.32 | 4.62E-08 |
| mRNA catabolic process (GO:0006402) | 213 | 9 | 0.21 | + | 42.36 | 2.87E-09 |
| oxidative phosphorylation (GO:0006119) | 109 | 4 | 0.11 | + | 36.79 | 0.0388 |
| RNA catabolic process (GO:0006401) | 247 | 9 | 0.25 | + | 36.53 | 1.04E-08 |
| establishment of protein localization to membrane (GO:0090150) | 289 | 8 | 0.29 | + | 27.75 | 0.00000203 |
| nucleobase-containing compound catabolic process (GO:0034655) | 372 | 9 | 0.37 | + | 24.26 | 0.000000366 |
| translation (GO:0006412) | 378 | 9 | 0.38 | + | 23.87 | 0.000000421 |
| peptide biosynthetic process (GO:0043043) | 403 | 9 | 0.4 | + | 22.39 | 0.000000734 |
| protein targeting (GO:0006605) | 371 | 8 | 0.37 | + | 21.62 | 0.000014 |
| heterocycle catabolic process (GO:0046700) | 429 | 9 | 0.43 | + | 21.03 | 0.00000126 |
| cellular nitrogen compound catabolic process (GO:0044270) | 430 | 9 | 0.43 | + | 20.98 | 0.00000129 |
| aromatic compound catabolic process (GO:0019439) | 444 | 9 | 0.44 | + | 20.32 | 0.0000017 |
| organic cyclic compound catabolic process (GO:1901361) | 478 | 9 | 0.48 | + | 18.88 | 0.00000323 |
| establishment of protein localization to organelle (GO:0072594) | 448 | 8 | 0.45 | + | 17.9 | 0.0000596 |
| peptide metabolic process (GO:0006518) | 519 | 9 | 0.52 | + | 17.39 | 0.00000659 |
| amide biosynthetic process (GO:0043604) | 526 | 9 | 0.52 | + | 17.15 | 0.0000074 |
| protein localization to membrane (GO:0072657) | 514 | 8 | 0.51 | + | 15.6 | 0.000171 |
| mRNA metabolic process (GO:0016071) | 690 | 9 | 0.69 | + | 13.08 | 0.0000767 |
| protein localization to organelle (GO:0033365) | 761 | 9 | 0.76 | + | 11.86 | 0.000178 |
| cellular amide metabolic process (GO:0043603) | 777 | 9 | 0.77 | + | 11.61 | 0.000212 |
| viral process (GO:0016032) | 784 | 9 | 0.78 | + | 11.51 | 0.000229 |
| symbiotic process (GO:0044403) | 876 | 9 | 0.87 | + | 10.3 | 0.00059 |
| cellular macromolecule catabolic process (GO:0044265) | 907 | 9 | 0.9 | + | 9.95 | 0.000793 |
| macromolecule catabolic process (GO:0009057) | 1048 | 9 | 1.05 | + | 8.61 | 0.00269 |
| intracellular protein transport (GO:0006886) | 997 | 8 | 0.99 | + | 8.04 | 0.0254 |
| organonitrogen compound biosynthetic process (GO:1901566) | 1382 | 10 | 1.38 | + | 7.25 | 0.00218 |
| cellular nitrogen compound biosynthetic process (GO:0044271) | 1638 | 10 | 1.63 | + | 6.12 | 0.0105 |
| negative regulation of gene expression (GO:0010629) | 1757 | 10 | 1.75 | + | 5.71 | 0.0199 |
| cellular localization (GO:0051641) | 2393 | 11 | 2.39 | + | 4.61 | 0.0382 |
| **Analysis Type:** | PANTHER Overrepresentation Test (Released 20200407) | | | | | |
| **Annotation Version and Release Date:** | GO Ontology database Released 2020-02-21 | | | | | |

Supplementary Table 9: **GO enrichment analysis of differentially variable genes between tumor and blood for CD4 naïve T cells in the lung cancer dataset.** We obtained the gene ontology (GO) terms significantly enriched in the top twenty genes ranked by increase in variance in tumor v. blood for CD4 naïve T cells (shown in Supplementary Table 3). The analysis was performed using the web service available at: http://geneontology.org/. Significance is calculated using Fisher's exact test, and $p$-values are Bonferroni corrected. Bgnd: Background count

**B Cells Memory**

| GO Biological Process Complete | Bgnd | Count | Expected | +/- | Fold Enrich | P-value |
|---|---|---|---|---|---|---|
| chaperone-mediated protein complex assembly (GO:0051131) | 18 | 3 | 0.02 | + | >100 | 1.00E-02 |
| regulation of cellular response to heat (GO:1900034) | 79 | 4 | 0.08 | + | 50.8 | 1.12E-02 |
| leukocyte activation involved in immune response (GO:0002366) | 615 | 7 | 0.61 | + | 11.42 | 1.30E-02 |
| cell activation involved in immune response (GO:0002263) | 619 | 7 | 0.62 | + | 11.35 | 1.35E-02 |
| response to organic substance (GO:0010033) | 3009 | 12 | 3 | + | 4 | 4.55E-02 |
| **Analysis Type:** | PANTHER Overrepresentation Test (Released 20190711) | | | | | |
| **Annotation Version and Release Date:** | GO Ontology database Released 2020-02-21 | | | | | |

Supplementary Table 10: **GO enrichment analysis of differentially variable genes between tumor and blood for memory B cells in the lung cancer dataset.** We obtained the gene ontology (GO) terms significantly enriched in the top twenty genes ranked by increase in variance in tumor v. blood for memory B cells (shown in Supplementary Table 4). The analysis was performed using the web service available at: http://geneontology.org/. Significance is calculated using Fisher's exact test, and $p$-values are Bonferroni corrected. Bgnd: Background count

**B Cells Naïve**

| GO Biological Process Complete | Bgnd | Count | Expected | +/- | Fold Enrich | P-value |
|---|---|---|---|---|---|---|
| response to cAMP (GO:0051591) | 101 | 6 | 0.1 | + | 59.6 | 6.52E-06 |
| cellular response to calcium ion (GO:0071277) | 85 | 5 | 0.08 | + | 59.02 | 2.14E-04 |
| SRP-dependent cotranslational protein targeting to membrane (GO:0006614) | 96 | 5 | 0.1 | + | 52.26 | 3.83E-04 |
| cotranslational protein targeting to membrane (GO:0006613) | 101 | 5 | 0.1 | + | 49.67 | 4.88E-04 |
| protein targeting to ER (GO:0045047) | 110 | 5 | 0.11 | + | 45.61 | 7.35E-04 |
| establishment of protein localization to endoplasmic reticulum (GO:0072599) | 114 | 5 | 0.11 | + | 44.01 | 8.72E-04 |
| viral transcription (GO:0019083) | 115 | 5 | 0.11 | + | 43.62 | 9.10E-04 |
| response to organophosphorus (GO:0046683) | 142 | 6 | 0.14 | + | 42.4 | 4.63E-05 |
| nuc.-transc. mRNA catab. proc., nonsense-med. decay (GO:0000184) | 120 | 5 | 0.12 | + | 41.81 | 1.12E-03 |
| response to calcium ion (GO:0051592) | 151 | 6 | 0.15 | + | 39.87 | 6.61E-05 |
| response to purine-containing compound (GO:0014074) | 157 | 6 | 0.16 | + | 38.34 | 8.27E-05 |
| viral gene expression (GO:0019080) | 132 | 5 | 0.13 | + | 38.01 | 1.76E-03 |
| protein localization to endoplasmic reticulum (GO:0070972) | 138 | 5 | 0.14 | + | 36.35 | 2.18E-03 |
| translational initiation (GO:0006413) | 142 | 5 | 0.14 | + | 35.33 | 2.51E-03 |
| protein targeting to membrane (GO:0006612) | 177 | 5 | 0.18 | + | 28.34 | 7.23E-03 |
| cellular response to metal ion (GO:0071248) | 193 | 5 | 0.19 | + | 25.99 | 1.10E-02 |
| nuclear-transcribed mRNA catabolic process (GO:0000956) | 194 | 5 | 0.19 | + | 25.86 | 1.12E-02 |
| mRNA catabolic process (GO:0006402) | 213 | 5 | 0.21 | + | 23.55 | 1.76E-02 |
| response to mechanical stimulus (GO:0009612) | 218 | 5 | 0.22 | + | 23.01 | 1.97E-02 |
| cellular response to inorganic substance (GO:0071241) | 221 | 5 | 0.22 | + | 22.7 | 2.10E-02 |
| RNA catabolic process (GO:0006401) | 247 | 5 | 0.25 | + | 20.31 | 3.58E-02 |
| response to metal ion (GO:0010038) | 370 | 6 | 0.37 | + | 16.27 | 1.16E-02 |
| cellular response to hormone stimulus (GO:0032870) | 612 | 7 | 0.61 | + | 11.48 | 1.26E-02 |
| response to bacterium (GO:0009617) | 691 | 7 | 0.69 | + | 10.16 | 2.79E-02 |
| response to organic cyclic compound (GO:0014070) | 926 | 8 | 0.92 | + | 8.67 | 1.46E-02 |
| response to organonitrogen compound (GO:0010243) | 1006 | 8 | 1 | + | 7.98 | 2.70E-02 |
| response to other organism (GO:0051707) | 1322 | 10 | 1.32 | + | 7.59 | 1.43E-03 |
| response to external biotic stimulus (GO:0043207) | 1324 | 10 | 1.32 | + | 7.58 | 1.45E-03 |
| response to biotic stimulus (GO:0009607) | 1356 | 10 | 1.35 | + | 7.4 | 1.81E-03 |
| response to nitrogen compound (GO:1901698) | 1089 | 8 | 1.09 | + | 7.37 | 4.85E-02 |
| interspecies interaction between organisms (GO:0044419) | 1964 | 12 | 1.96 | + | 6.13 | 4.25E-04 |
| cellular nitrogen compound biosynthetic process (GO:0044271) | 1638 | 10 | 1.63 | + | 6.13 | 1.04E-02 |
| RNA metabolic process (GO:0016070) | 1679 | 10 | 1.67 | + | 5.98 | 1.30E-02 |
| cellular macromolecule biosynthetic process (GO:0034645) | 1694 | 10 | 1.69 | + | 5.92 | 1.41E-02 |
| macromolecule biosynthetic process (GO:0009059) | 1753 | 10 | 1.75 | + | 5.72 | 1.93E-02 |
| negative regulation of gene expression (GO:0010629) | 1757 | 10 | 1.75 | + | 5.71 | 1.97E-02 |
| response to external stimulus (GO:0009605) | 2443 | 12 | 2.43 | + | 4.93 | 4.76E-03 |
| negative regulation of macromolecule metabolic process (GO:0010605) | 2682 | 12 | 2.67 | + | 4.49 | 1.32E-02 |
| negative regulation of metabolic process (GO:0009892) | 2942 | 12 | 2.93 | + | 4.09 | 3.58E-02 |
| response to stress (GO:0006950) | 3572 | 13 | 3.56 | + | 3.65 | 3.63E-02 |
| **Analysis Type:** | PANTHER Overrepresentation Test (Released 20190711) | | | | | |
| **Annotation Version and Release Date:** | GO Ontology database Released 2020-02-21 | | | | | |

Supplementary Table 11: **GO enrichment analysis of differentially variable genes between tumor and blood for naïve B cells in the lung cancer dataset.** We obtained the gene ontology (GO) terms significantly enriched in the top twenty genes ranked by increase in variance in tumor v. blood for naïve B cells (shown in Supplementary Table 5). The analysis was performed using the web service available at: http://geneontology.org/. Significance is calculated using Fisher's exact test, and $p$-values are Bonferroni corrected. Bgnd: Background count

### Supplementary Note 1

### Motivating the power-law relationship between embedded and original local radii

We motivate the connection between density-preservation and a power-law relationship between the original and embedding local radii with an example. Suppose for a point $x \in \mathbb{R}^d$ in the original $d$-dimensional dataset, the $K$ points in its neighborhood are uniformly distributed in a ball of radius $\gamma_d$ and volume $V \propto \gamma_d^d$.

Now, suppose we want to embed the dataset into $s < d$ dimensions while preserving structure and density. This means we want $x$ and its neighbors to be mapped to an $s$-dimensional ball of uniform density with radius $\gamma_s$, and, to preserve the *density* of the $K$-neighborhood of $x$, the volume of the $s$-dimensional ball should still be $V$. Since $V \propto \gamma_s^s$, this suggests a power law relationship between $\gamma_s$ and $\gamma_d$, i.e. $\gamma_s \propto \gamma_d^{d-s}$. Taking logarithms, $\log \gamma_s = (d-s)\log \gamma_d + \beta$ for some $\beta$.

Drawing the analogy between the local radius we introduced in Methods and $\gamma$ above, density preservation thus corresponds to a power law relationship between the local radii in the original and the embedded datasets.

### Supplementary Note 2

### Gradient derivation for den-SNE and densMAP

The core of our density-preserving tools lies in the optimization of the Pearson correlation between the log local radius of points in the original dataset and in the embedding (see Methods). Here we compute the gradient of this correlation with respect to the embedding coordinates for optimization. Let $X = \{x_i\}_{i=1}^N$ be our original dataset and $y_i = s(x_i)$ be our embedding, where $s \in \{\text{den-SNE}, \text{densMAP}\}$ is our algorithm of choice.

Let $\{R_i^o\}_{i=1}^N$ and $\{R_i^e\}_{i=1}^N$ be measures of pointwise density in the original and embedded spaces respectively. We discuss specific density functions at the end, but allow full generality here. Let $r_i^e = \log R_i^e$. We center the original densities, so we let $r_i^o = \log R_i^o - N^{-1}\sum_{k=1}^N \log R_i^o$.

Since we want the densities in the embedded and original dataset to have a power-law relationship, (Supplementary Note 1) we maximize the correlation between $r^o = \{r_i^o\}_{i=1}^N$ and $r^e = \{r_i^e\}_{i=1}^N$, denoted $\rho_{e,o}$. We write:

$$\rho_{e,o} = \frac{\text{Cov}(r^e, r^o)}{\sigma^o \sigma^e} = \frac{\sum_{k=1}^N (r_k^e - \mu^e) r_k^o}{s^o (N-1)^{\frac{1}{2}} \left[\sigma^2 + \sum_{k=1}^N (r_k - \mu^e)^2\right]^{\frac{1}{2}}} \tag{1}$$

where $\mu^e$ is the average of $r^e$; $\sigma^o$ and $\sigma^e$ are the sample standard deviations of $r^o$ and $r^e$ respectively, and $\sigma^2$ is a user-specified constant for regularization (this ensures that the standard deviation of the embedded local radii does not go to zero).

Now, we compute the gradient of the correlation with respect to pairwise squared distances of the embedded datapoints, $d_{ij}^2 = \|y_i - y_j\|^2$:

$$\frac{\partial \rho_{e,o}}{\partial d_{ij}^2} = (s^o)^{-1}(N-1)^{-\frac{1}{2}} \left[\widetilde{\text{Var}}(r^e)^{-\frac{1}{2}} \frac{\partial \widetilde{\text{Cov}}(r^e, r^o)}{\partial d_{ij}^2} - \frac{1}{2}\widetilde{\text{Cov}}(r^e, r^o)\widetilde{\text{Var}}(r^e)^{-\frac{3}{2}} \frac{\partial \widetilde{\text{Var}}(r^e)}{\partial d_{ij}^2}\right],$$

where $\widetilde{\text{Var}}(r^e) = (N-1)\left[\sigma^2/(N-1) + \text{Var}(r^e)\right]$ and $\widetilde{\text{Cov}}(r^e, r^o) = (N-1)\text{Cov}(r^e, r^o)$ (these are sample variances and covariances, hence the normalization by $N-1$ instead of $N$).

Now consider the component parts:

$$
\begin{aligned}
\frac{\partial \widetilde{\mathrm{Cov}}(r^e, r^o)}{\partial d_{ij}^2} &= \sum_{k=1}^{N} r_k^o \left( \frac{\partial r_k^e}{\partial d_{ij}^2} - \frac{\partial \mu^e}{\partial d_{ij}^2} \right) \\
&= \sum_{k=1}^{N} r_k^o \frac{\partial r_k^e}{\partial d_{ij}^2} - \frac{\partial \mu^e}{\partial d_{ij}^2} \sum_{k=1}^{N} r_k^o \qquad (2) \\
&= \sum_{k=1}^{N} r_k^o \frac{\partial r_k^e}{\partial d_{ij}^2},
\end{aligned}
$$

where the second term in (2) is zero since $r^o$ is centered. Similarly,

$$
\begin{aligned}
\frac{\partial \widetilde{\mathrm{Var}}(r^e)}{\partial d_{ij}^2} &= 2 \sum_{k=1}^{N} (r_k^e - \mu^e) \left( \frac{\partial r_k^e}{\partial d_{ij}^2} - \frac{\partial \mu^e}{\partial d_{ij}^2} \right) \\
&= 2 \sum_{k=1}^{N} (r_k^e - \mu^e) \frac{\partial r_k^e}{\partial d_{ij}^2} - 2 \frac{\partial \mu^e}{\partial d_{ij}^2} \sum_{k=1}^{N} (r_k^e - \mu^e) \qquad (3) \\
&= 2 \sum_{k=1}^{N} (r_k^e - \mu^e) \frac{\partial r_k^e}{\partial d_{ij}^2},
\end{aligned}
$$

where, similar to before, the second sum in (3) is zero.

For many density functions (and certainly the one we use), $r_i$ will only depend on $\{d_{ik}, d_{ki}\}_{k=1}^N$, so we can further simplify the above expressions to:

$$
\begin{aligned}
\frac{\partial \widetilde{\mathrm{Cov}}(r^e, r^o)}{\partial d_{ij}^2} &= r_i^o \frac{\partial r_i^e}{\partial d_{ij}^2} + r_j^o \frac{\partial r_j^e}{\partial d_{ij}^2} \\
\frac{\partial \widetilde{\mathrm{Var}}(r^e)}{\partial d_{ij}^2} &= 2 \left( (r_i^e - \mu^e) \frac{\partial r_i^e}{\partial d_{ij}^2} + (r_j^e - \mu^e) \frac{\partial r_j^e}{\partial d_{ij}^2} \right).
\end{aligned}
$$

Under this scenario, putting this all together, we get:

$$
\frac{\partial \rho_{e,o}}{\partial d_{ij}^2} = \frac{\widetilde{\mathrm{Var}}(r^e) \left( r_i^o \frac{\partial r_i^e}{\partial d_{ij}^2} + r_j^o \frac{\partial r_j^e}{\partial d_{ij}^2} \right) - \widetilde{\mathrm{Cov}}(r^e, r^o) \left( (r_i^e - \mu^e) \frac{\partial r_i^e}{\partial d_{ij}^2} + (r_j^o - \mu^e) \frac{\partial r_j^e}{\partial d_{ij}^2} \right)}{s^o (N-1)^{\frac{1}{2}} \widetilde{\mathrm{Var}}(r^e)^{\frac{3}{2}}}. \qquad (4)
$$

The measure of local density for the embedded points we have used is the squared distance, weighted by the embedding distribution $Q$ for either t-SNE or UMAP:

$$
R_k^e = \left( \sum_{\ell=1}^{N} (1 + a d_{k\ell}^{2b})^{-1} \right)^{-1} \sum_{m=1}^{N} d_{km}^2 (1 + a d_{km}^{2b})^{-1} = \mathcal{Z}_k^{-1} \sum_{m=1}^{N} d_{km}^2 (1 + a d_{km}^{2b})^{-1},
$$

where $\mathcal{Z}_k = \sum_{m=1}^{N} (1 + a d_{km}^{2b})^{-1}$. We also write $\widetilde{Q}_{k\ell} = \mathcal{Z}_k^{-1} (1 + a d_{k\ell}^{2b})^{-1}$. Note that these are related to the $Q$ matrix and $\mathcal{Z}$ partition functions of t-SNE and UMAP, equations (5) and (6) in Methods, by

$$
\begin{aligned}
\mathcal{Z} &= \sum \mathcal{Z}_i \\
Q_{ij} &= \widetilde{Q}_{ij} \frac{\mathcal{Z}_i}{\mathcal{Z}}.
\end{aligned}
$$

To finish (4), we need to evaluate $\frac{\partial r_i^e}{\partial d_{ij}^2}$.

$$\begin{aligned}
\frac{\partial r_i^e}{\partial d_{ij}^2} &= \frac{\partial}{\partial d_{ij}^2}\log(\mathcal{Z}_i R_i^e) - \frac{\partial}{\partial d_{ij}^2}\log \mathcal{Z}_i \\
&= (R_i^e \mathcal{Z}_i)^{-1}\frac{\partial(\mathcal{Z}_i R_i^e)}{\partial d_{ij}^2} - \mathcal{Z}_i^{-1}\frac{\partial \mathcal{Z}_i}{\partial d_{ij}^2} \\
&= (R_i^e \mathcal{Z}_i)^{-1}\left[(1+ad_{ij}^{2b})^{-1} - abd_{ij}^{2b-2}d_{ij}^2(1+ad_{ij}^{2b})^{-2})\right] + \mathcal{Z}_i^{-1}abd_{ij}^{2b-2}(1+ad_{ij}^{2b})^{-2} \\
&= \frac{\widetilde{Q}_{ij}}{R_i^e}(1-abd_{ij}^{2b}(1+ad_{ij}^{2b})^{-1}) + abd_{ij}^{2b-2}\widetilde{Q}_{ij}^2\mathcal{Z}_i \\
&= (1+ad_{ij}^{2b})^{-1}\widetilde{Q}_{ij}\left[1+ad_{ij}^{2b} - ad_{ij}^{2b}\right] + \widetilde{Q}_{ij}^2\mathcal{Z}_i \\
&= \widetilde{Q}_{ij}^2\mathcal{Z}_i\left[1+\frac{1+ad_{ij}^{2b}(1-b)}{R_i^e}\right]
\end{aligned}$$

Note that when $a = b = 1$, as in t-SNE, this simplifies to:

$$\frac{\partial r_i^e}{\partial d_{ij}^2} = \widetilde{Q}_{ij}^2\mathcal{Z}_i\left[1+\frac{1}{R_i^e}\right]. \tag{5}$$

As discussed in Methods, in both den-SNE and densMAP, for the sake of efficiency, we assume for the local radius computation that, for a point $i$ with embedding $y_i$ and original coordinates $x_i$, $d_{ij}^2 \neq 0$ only for $j$ such that $i$ and $j$ are in the edge set $E$ of the $k$-nearest neighbors graph produced by each algorithm. Since the objective functions of each algorithm prioritize preserving local structure, they should encourage $i$ and $j$ to be nearest neighbors in the embedding as well, and so we only need to consider density with respect to those points.

We can now write the full den-SNE gradient by combining the gradient of the correlation with the gradient of the original t-SNE objective function [1] (we discuss adaptations needed for the densMAP gradient below):

$$\nabla_{y_i}\mathcal{L}^{\text{den-SNE}} = \sum_{\{i,j\}\in E} P_{ij}Q_{ij}\mathcal{Z}(y_i - y_j) - \sum_{j\neq i} Q_{ij}^2\mathcal{Z}(y_i-y_j) - \lambda \sum_{\{i,j\}\in E}\frac{\partial \rho_{e,o}}{\partial d_{ij}^2}(y_i - y_j),$$

where $\lambda$ is a user-provided parameter that determines the weight of the density-preservation objective, and $\frac{\partial \rho_{e,o}}{\partial d_{ij}^2}$ is given in (4) with (5) plugged in.

#### 2.1 Stochastic gradient descent for densMAP

Here we detail how densMAP adapts the stochastic gradient descent (SGD) formulation of UMAP. The cross-entropy loss function for UMAP and its gradient with respect to the squared distance $d_{ij}^2$ between points $i, j$ in the embedding [2], given $E$, the set of edges in the nearest-neighbors graph, is:

$$\begin{aligned}
\mathcal{L} &= \sum_{\{i,j\}\in E} \underbrace{P_{ij}\log Q_{ij}}_{\text{attractive}} + \underbrace{(1-P_{ij})\log(1-Q_{ij})}_{\text{repulsive}} \\
\frac{\partial \mathcal{L}}{\partial d_{ij}^2} &= P_{ij}\frac{\partial}{\partial d_{ij}^2}[\log Q_{ij}] + (1-P_{ij})\frac{\partial}{\partial d_{ij}^2}[\log(1-Q_{ij})],
\end{aligned}$$

where $P$ and $Q$ are the distributions on the original and embedded data respectively. Note that in UMAP, unlike t-SNE, the value $Q_{ij}$ depends *only* on distance $d_{ij}^2$ and does not involve a normalization term over all edges.

To optimize the attractive term of the objective function, at each step, UMAP draws an edge $\{i, j\} \in E$ randomly according to the distribution $P$, and computes the gradient $\frac{\partial}{\partial d_{ij}^2}(\log Q_{ij})(y_i - y_j)$. This means that, over the course of the optimization, edge $\{i, j\}$ will be chosen with proportion $P_{ij}/Z$, where $Z = \sum_{i\neq i} P_{ij}$. To estimate the repulsive term, a set of points $S = \{k_s\}_{s=1}^{n_s}$ is chosen *uniformly* at random and the algorithm computes the gradient $\frac{1}{|S|}\sum_{k\in S}\frac{\partial}{\partial d_{ik}^2}[\log(1-Q_{ik})](y_i - y_j)$. The size of $S$ is a tunable parameter $n_s$.

Now, incorporating the density-preservation term into this objective function means taking the gradient of the correlation (1) and adding it to the UMAP gradient. The full gradient becomes:

$$\nabla_{y_i}\mathcal{L} = \sum_{\{i,j\}\in E}\left(P_{ij}\frac{\partial}{\partial d_{ij}^2}[\log Q_{ij}] + (1-P_{ij})\frac{\partial}{\partial d_{ij}^2}[\log(1-Q_{ij})] + \lambda\frac{\partial \rho_{e,o}}{\partial d_{ij}^2}\right)(y_i - y_j).$$

Note that, for the correlation term of the optimization, each *edge* is given equal weight (i.e. the term is not weighted by $P_{ij}$). Since, in the stochastic descent algorithm of UMAP, an edge is actually chosen with proportion $P_{ij}/Z$, we *re-weight* the gradient estimate of the correlation term by multiplying by $Z/(NP_{ij})$ (we divide by the number of points $N$ for numerical stability, since $Z$ grows with $N$).

The densMAP gradient estimate for an edge $\{i,j\}$ at each iteration of the SGD is then:

$$\nabla_{y_i}\mathcal{L}|_{\{i,j\}} = \left(\frac{\partial}{\partial d_{ij}^2}[\log Q_{ij}] + \lambda\frac{Z}{NP_{ij}}\frac{\partial \rho_{e,o}}{\partial d_{ij}^2} - \frac{1}{|S|}\sum_{k\in S}\frac{\partial}{\partial d_{ik}^2}[\log(1-Q_{ik})]\right)(y_i - y_j),$$

where $S$ is a set of edges adjacent to $i$ chosen uniformly at random. This ensures that, over the course of the optimization, the edges are weighted equally when optimizing the correlation term.

Next, we consider the $Q$ distribution. Since, unlike t-SNE, UMAP does *not* normalize $Q_{ij}$ over all the edges, it treats the term as a Bernoulli random variable over each edge. For the calculation of the local radius, however, we need a probability distribution over the nearest neighbors of each point. In other words, we need $Q_{ij}/\sum_{k:\{i,k\}\in E} Q_{ik} = Q_{ij}/\mathcal{Z}_i$. To achieve this, we compute $\mathcal{Z}_i$ at the start of each epoch and take it as fixed for all the edges that are updated in that epoch, which is akin to performing coordinate descent with the update for $\mathcal{Z}_i$ happening once per epoch. Similarly, we compute the local radius $R_i^e$, and global variance and covariance terms at the start of each epoch. These techniques allow us to use SGD to optimize densMAP in a similar manner as UMAP.

### Supplementary Note 3

### Theoretical motivation for the local radius

Here, we motivate density-preservation by more rigorously showing that t-SNE does not preserve density due to its use of a constant perplexity for choosing the length-scale. Unfortunately, UMAP's use of a non-standard distribution for its $P$ matrix (the re-scaled exponential distribution) precludes this analysis from being extended to UMAP. The setup is as follows.

Assume that for a given point $X$, we draw its $n$-nearest neighbors (which are used in the computation of t-SNE's $P$ matrix) as iid random variables $\mathbf{X} = \{X_1, \ldots, X_n\}$ where each $X_i \in \mathbb{R}^d$ is drawn from a Gaussian distribution with mean $X$ and covariance matrix $\Sigma$. Let $P_X$ be a row of the un-symmetrized probability matrix induced by t-SNE, as in (1) in Methods:

$$(P_X)_j = Z_X^{-1}\exp\Big(-\|X - X_j\|^2/\sigma_X^2\Big)$$
$$Z_X = \sum_{j=1}^{n}\exp\Big(-\|X - X_j\|^2/\sigma_X^2\Big).$$

The length-scale term $\sigma_X$ is chosen to make the perplexity, Perp, constant as in (7) in Methods:

$$\log \mathrm{Perp} = \mathscr{H}_X = -Z_X^{-1}\sum_{j=1}^{n}(P_X)_j\log(P_X)_j + \log Z_X, \tag{6}$$

where $\mathscr{H}_X$ is the *entropy* of $P_X$.

#### 3.1 Scaling of $\sigma$ in t-SNE

We can consider $\sigma_X$ as a function of the covariance of the generative model for $\mathbf{X}$, i.e. $\sigma_X = \sigma(\Sigma)$, since it is based on the length-scale, so $\sigma_X$ itself can be thought of as a random variable. Notably, Vladymyrov and Carreira-Perpiñán (2013) [3] show that $\sigma$ has a unique value that satisfies (6), so we just need to find *any* $\sigma$ that satisfies the equation.

We first show that $\sigma$ scales as the variance.

**Proposition 3.1.1.** *Suppose, we draw $\mathbf{X} = \{X_1, \ldots, X_n\}$, $X_i \in \mathbb{R}^d$ from a Gaussian distribution $\mathcal{N}_X$ with mean $X$ and covariance $\Sigma$, and given $\alpha > 0$, we now draw $\mathbf{Y} = \{Y_1, \ldots, Y_n\}$, $Y_i \in \mathbb{R}^d$ from a Gaussian distribution $\mathcal{N}_Y$ with mean $Y$ and covariance $\alpha\Sigma$. Let $\sigma_X$ be chosen so that $\mathscr{H}_X = \log \mathrm{Perp}$ for a constant value of Perp. Then, for any $\phi, \delta \in (0, 1/4)$, with $n = O(\log(1/\delta)/\phi^2)$, setting $\sigma_Y = \alpha\sigma_X$ will yield $(1-\phi)\mathscr{H}_X < \mathscr{H}_Y < (1+\phi)\mathscr{H}_X$ with probability at least $1-\delta$.*

Our proof involves approximating the sums on the right-hand side of (6) by expectations over the generating Gaussian distribution. To that end, we define:

$$\mathcal{G}_X(\sigma^2) = \frac{\mathbb{E}_{z\sim\mathcal{N}_X}\left[\exp\left(-\|z-X\|^2/\sigma^2\right)\|z-X\|^2/\sigma^2\right]}{\mathbb{E}_{z\sim\mathcal{N}_X}\left[\exp\left(-\|z-X\|^2/\sigma^2\right)\right]} + \log \mathbb{E}_{z\sim\mathcal{N}_X}\left[\exp\left(-\|z-X\|^2/\sigma^2\right)\right]. \tag{7}$$

We will show that $\mathscr{H}_X - \log n \to \mathcal{G}_X$ in the large-sample limit. We first show the behavior of $\mathcal{G}$ under dilations of the length-scale, i.e. $\mathcal{G}(\alpha\sigma^2)$:

**Lemma 3.1.2.** *Given the distributions $\mathcal{N}_X$ and $\mathcal{N}_Y$ defined above, and length-scale term $\sigma$:*

$$\mathcal{G}_Y(\alpha\sigma^2) = \mathcal{G}_X(\sigma^2).$$

*That is, scaling $\sigma$ by the same amount as the covariance results in a constant value for $\mathcal{G}$.*

*Proof.* (Lemma 3.1.2) Introducing the simplifying notations $d(z) := \|z - X\|$, $p(z,\sigma) := \exp\left(-d(z)^2/\sigma^2\right)$, and $\mathbb{E}_X[\cdot] := \mathbb{E}_{z\sim\mathcal{N}_X}[\cdot]$, we write:

$$\mathcal{G}_X(\sigma^2) = \frac{\mathbb{E}_X\left[p(z,\sigma)\frac{d^2(z)}{\sigma^2}\right]}{\mathbb{E}_X[p(z,\sigma)]} + \log \mathbb{E}_X[p(z,\sigma)]. \tag{8}$$

Letting $f(z;\mu,\Sigma)$ denote the pdf of the multivariate normal distribution with mean $\mu$ and covariance $\Sigma$, we can thus compute the expectations explicitly:

$$\begin{aligned}\mathbb{E}_X\left[p(z,\sigma)\frac{d^2(z)}{\sigma^2}\right] &= \frac{1}{\sigma^2}\int f(z;X,\Sigma)p(z,\sigma)d^2(z)\,\mathrm{d}z \\ &\propto \frac{1}{\sigma^2 \det \Sigma^{1/2}}\int \exp\left(-\frac{1}{2}((z-X)^T(\Sigma^{-1}+2\sigma^{-2}I)(z-X))\right)d^2(z)\,\mathrm{d}z\,. \end{aligned} \tag{9}$$

The factors hidden by the proportionality symbol throughout are constants that depend only on the dimension and not on $X$, $\sigma$, or $\Sigma$.

We show now that we can, without loss of generality, assume that the covariance matrix $\Sigma$ is diagonal. Intuitively, this is because the calculation of $\sigma$ relies only on *distances*, which are invariant under orthogonal transformations, and we can transform to a coordinate system where $\Sigma$ is diagonal. More formally, assume we have a non-diagonal covariance matrix. Then we can take its singular value decomposition $\Sigma = U\Lambda U^T$ where $\Lambda$ is diagonal and $U$ orthonormal. Then, replacing the integration variable in (9) with $\theta = Uz$ and $\Phi = UX$, we see:

$$\mathbb{E}_X\left[p(z,\sigma)\frac{d^2(z)}{\sigma^2}\right] \propto \frac{1}{\sigma^2 \det \Lambda^{1/2}}\int \exp\left(-\frac{1}{2}((\theta-\Phi)^T(\Lambda^{-1}+2\sigma^{-2}I)(\theta-\Phi))\right)d^2(\theta)\,\mathrm{d}\theta\,, \tag{10}$$

which is identical to (9).

The exponential term in the integral in (9) is the (unnormalized) pdf of a normal distribution with mean $X$ and covariance matrix $(\Sigma^{-1} + 2\sigma^{-2}I)^{-1} = \sigma^2\Sigma(\sigma^2 I + 2\Sigma)^{-1}$ (by the Woodbury matrix identity). Thus, (9) is:

$$\propto C(\sigma,\Sigma)\frac{1}{\sigma^2 \det \Sigma^{1/2}}\int f(x; X, \sigma^2\Sigma(\sigma^2 I + 2\Sigma)^{-1})\|x - X\|^2 \, \mathrm{d}x \,,$$

where we define $C$ as the normalization factor for the normal distribution $f$ inside the integral. The integral is thus the expectation of the sum of the variances in each dimension of the distribution, i.e. the total variance, given by the trace of the variance matrix:

$$\mathbb{E}_X\left[p(x,\sigma)\frac{d^2(x)}{\sigma^2}\right] \propto \underbrace{\det\left[\sigma\Sigma^{1/2}(\sigma^2 I + 2\Sigma)^{-1/2}\right]}_{C(\sigma,\Sigma)}\frac{1}{\sigma^2\det\Sigma^{1/2}}\underbrace{\mathrm{Tr}\left[\sigma^2\Sigma(\sigma^2 I + 2\Sigma)^{-1}\right]}_{\text{total variance of } f}. \tag{11}$$

Similarly,

$$\mathbb{E}_X\left[p(x,\sigma)\right] \propto \frac{1}{\det\Sigma^{1/2}}\int \exp\left(-\frac{1}{2}((x-X)^T(\Sigma^{-1} + 2\sigma^{-2}I)(x-X))\right)\mathrm{d}x \,,$$

and thus the integral is just the normalization of $f(x; X, \sigma^2\Sigma(\sigma^2 I + 2\Sigma)^{-1})$, so:

$$\mathbb{E}_X\left[p(x,\sigma)\right] \propto \underbrace{\det\left[\sigma\Sigma^{1/2}(\sigma^2 I + 2\Sigma)^{-1/2}\right]}_{C(\sigma,\Sigma)}\frac{1}{\det\Sigma^{1/2}}. \tag{12}$$

We thus have the form of $\mathcal{G}_X(\sigma^2)$ by plugging (11) and (12) into (8):

$$\mathcal{G}_X(\sigma^2) = C_1\,\mathrm{Tr}\left[\Sigma(\sigma^2 I + 2\Sigma)^{-1}\right] + C_2\log\left(\frac{\det\left[\sigma\Sigma^{1/2}(\sigma^2 I + 2\Sigma)^{-1/2}\right]}{\det\Sigma^{1/2}}\right), \tag{13}$$

where $C_1$ and $C_2$ are constants that do not depend on $X$, $\sigma$, or $\Sigma$ (and will be used throughout, but can take on different values). Now, we turn to $Y$, whose neighbors are drawn from a normal distribution with mean $Y$ and covariance $\alpha\Sigma$. Repeating the analyses above, it is easy to see then that:

$$\mathcal{G}_Y(\tau^2) = C_1\,\mathrm{Tr}\left[\alpha\Sigma(\tau^2 I + 2\alpha\Sigma)^{-1}\right] + C_2\log\left(\frac{\det\left[\tau(\alpha\Sigma)^{1/2}(\tau^2 I + 2\alpha\Sigma)^{-1/2}\right]}{\det(\alpha\Sigma)^{1/2}}\right). \tag{14}$$

Plugging in $\tau = \alpha\sigma$, we see:

$$\begin{aligned}
\mathcal{G}_Y(\alpha\sigma^2) &= C_1\,\mathrm{Tr}\left[\alpha\Sigma(\alpha\sigma^2 I + 2\alpha\Sigma)^{-1}\right] + C_2\log\left(\frac{\det\left[\alpha\sigma(\alpha\Sigma)^{1/2}(\alpha\sigma^2 I + 2\alpha\Sigma)^{-1/2}\right]}{\det(\alpha\Sigma)^{1/2}}\right)\\
&= C_1\,\mathrm{Tr}\left[\cancel{\alpha}\Sigma\cancel{\alpha^{-1}}(\sigma^2 I + 2\Sigma)^{-1}\right] + C_2\log\left(\frac{\cancel{\alpha^{d/2}}\det\left[\sigma\Sigma^{1/2}(\sigma^2 I + 2\Sigma)^{-1/2}\right]}{\cancel{\alpha^{d/2}}\det(\Sigma)^{1/2}}\right)\\
&= \mathcal{G}_X(\sigma^2).
\end{aligned}$$

□

Thus, when the variance of the underlying distribution is scaled by $\alpha$, the length-scale selection needs to also scale by $\alpha$ to keep $\mathcal{G}$ constant.

Next, we now show that, for sufficiently large $n$, the entropy $\mathscr{H}_X$ calculated on a sample and scaled appropriately by $n$, does not differ much from $\mathcal{G}_X$.

**Lemma 3.1.3.** *Suppose we draw* $\mathbf{X} = \{X_1, \ldots, X_n\}$ *iid from* $\mathcal{N}_X$. *Then, for any* $\alpha, \delta \in (0, 1/4)$ *and* $n > O\left(\frac{\log(1/\delta)}{\epsilon^2}\right)$, *we have:*

$$(1-\alpha)\mathcal{G}_X + \log(1-\alpha) < \mathscr{H}_X - \log n < (1+\alpha)\mathcal{G}_X + \log(1+\alpha)$$

*with probability at least* $1 - \delta$.

*Proof.* (Lemma 3.1.3) Using our notation from above, we have:

$$\mathcal{G}_X(\sigma^2) = \frac{\mathbb{E}_X[p(z,\sigma)d^2(z)]}{\sigma^2\mathbb{E}_X[p(z,\sigma)]} + \log \mathbb{E}_X[p(z,\sigma)]$$
$$\mathscr{H}_X = \frac{\sum_{j=1}^n p(X_j,\sigma)d^2(X_j)}{\sigma^2\sum_{j=1}^n p(X_j,\sigma)} + \log\left(\sum_{j=1}^n p(X_j,\sigma)\right).$$

We use Hoeffding's inequality to show that:

$$\mathbb{E}_X[p(z,\sigma)d^2(z)] \approx \frac{1}{n}\sum_{j=1}^n p(X_j,\sigma)d^2(X_j)$$
$$\mathbb{E}_X[p(z,\sigma)] \approx \frac{1}{n}\sum_{j=1}^n p(X_j,\sigma).$$

Given $X_1,\ldots,X_n$ iid random variables drawn from a distribution $F$, bounded in $[0,s]$, Hoeffding's inequality quantifies how far the sample mean $\overline{X} = \frac{1}{n}\sum_i X_i$ deviates from its expectation $\mu = \mathbb{E}_{X\sim F}[X]$:

$$\Pr[\overline{X} > (1+\epsilon)\mu] < \exp\left(-\frac{\delta^2 n\mu}{3s}\right)$$
$$\Pr[\overline{X} < (1-\epsilon)\mu] < \exp\left(-\frac{\delta^2 n\mu}{2s}\right).$$

Taking the weaker bound (the first one) and setting the probability to $\delta$, we can solve for $n$ to see that $\overline{X}$ will be between $(1\pm\epsilon)\mu$ when $n > \frac{3s\log(1/\delta)}{\mu\epsilon^2}$. Now, we note that the random variable $p(z,\sigma)$ lies within $[0,1]$, and the random variable $p(z,\sigma)\frac{d^2(z)}{\sigma^2}$ lies within $[0,1/e]$, so we can set the range $s$ to be 1. Thus, with sufficiently large $n$, we have that:

$$(1-\epsilon)\mathbb{E}_X[p(z,\sigma)d^2(z)] < \frac{1}{n}\sum_{j=1}^n p(X_j,\sigma)d^2(X_j) < (1+\epsilon)\mathbb{E}_X[p(z,\sigma)d^2(z)], \tag{15}$$

and

$$(1-\epsilon)\mathbb{E}_X[p(z,\sigma)] < \frac{1}{n}\sum_{j=1}^n p(X_j,\sigma) < (1+\epsilon)\mathbb{E}_X[p(z,\sigma)]. \tag{16}$$

Now, the above two equations (15) and (16) provide bounds for the numerator and the denominator of the first term of $\mathcal{G}_X(\sigma^2)$, respectively. Thus, in the worst case, we see that:

$$\frac{(1-\epsilon)\mathbb{E}_X[p(z,\sigma)d^2(z)/\sigma^2]}{(1+\epsilon)\mathbb{E}_X[p(z,\sigma)]} + \log(1-\epsilon) + \log\mathbb{E}[p(z,\sigma)]$$
$$< \mathscr{H}_X - \log n$$
$$< \frac{(1+\epsilon)\mathbb{E}_X[p(z,\sigma)d^2(z)/\sigma^2]}{(1-\epsilon)\mathbb{E}_X[p(z,\sigma)]} + \log(1+\epsilon) + \log\mathbb{E}[p(z,\sigma)].$$

To write these bounds in terms of $\mathcal{G}_X$, we note that $1+2\sqrt{\epsilon} > \frac{1+\epsilon}{1-\epsilon}$ and $1-2\sqrt{\epsilon} < \frac{1-\epsilon}{1+\epsilon}$ for $\epsilon < 1/4$, we can combine the above and compare $\mathscr{H}_X$ and $\mathcal{G}_X$:

$$(1-2\sqrt{\epsilon})\mathcal{G}_X + \log\big(1-2\sqrt{\epsilon}\big) < \mathscr{H}_X(\sigma) - \log n < (1+2\sqrt{\epsilon})\mathcal{G}_X + \log\big(1+2\sqrt{\epsilon}\big)..$$

Setting $\alpha = 2\sqrt{\epsilon}$ gives us the desired bounds. □

With these two results, we can prove Proposition 3.1.1.

*Proof.* (Proposition 3.1.1) By Lemma 3.1.2, $\mathcal{G}_Y(\alpha\sigma_X^2) = \mathcal{G}_X(\sigma_X^2)$. By our concentration bounds in Lemma 3.1.3, we take enough samples so that $\mathscr{H}_X(\sigma_X^2)$ is within a $(1+\beta)$ multiplicative factor of $\mathcal{G}_X$. Similarly, with as many samples, we know that $\mathscr{H}_Y(\alpha\sigma^2)$ is within a $(1+\beta)$ multiplicative factor of $\mathcal{G}_Y$. Thus in the worst case:

$$(1-\beta)^2\mathscr{H}_X < \mathscr{H}_Y < (1+\beta)^2\mathscr{H}_X$$

Taking $\beta = \frac{1}{3}\phi$ (which allows us to discard the quadratic terms), we see the above can be relaxed to

$$(1-\phi)\mathscr{H}_X < \mathscr{H}_Y < (1+\phi)\mathscr{H}_X,$$

as required. □

For the remainder of this section we will assume that the quantities calculated can be well approximated by their expectations, as we do above by approximating $\mathscr{H}_X$ with $\mathcal{G}_X$. For all the below propositions, Hoeffding's inequality can be used as in Lemma 3.1.3 to achieve arbitrarily close approximations, logarithmic in the number of samples.

We extend the above analysis to the case of *uniformly* distributed data.

**Proposition 3.1.4.** *Let $X$ be such that its $n$-nearest neighbors are distributed uniformly in a ball $B_X$ of radius $\gamma$, and $Y$ another point whose neighbors are distributed uniformly in a ball $B_Y$ of radius $\sqrt{\alpha}\gamma$. Then, $\mathcal{G}_Y(\alpha\sigma^2) = \mathcal{G}_X(\sigma^2)$ where $\mathcal{G}$ is as defined analogously to* (8)*:*

$$\mathcal{G}_X(\sigma^2) = \frac{\mathbb{E}_{z\sim B_X}\left[p(z,\sigma)\frac{d^2(z)}{\sigma^2}\right]}{\mathbb{E}_{z\sim B_X}[p(z,\sigma)]} + \log \mathbb{E}_{z\sim B_X}[p(z,\sigma)].$$

*Proof.* As in Proposition 3.1.1, we then explicitly compute the terms $\mathbb{E}_X\left[p(z,\sigma)\frac{d^2(z)}{\sigma^2}\right]$ and $\mathbb{E}_X\left[p(z,\sigma)\right]$, assuming a uniform distribution now instead of a Gaussian:

$$\begin{aligned}
\mathbb{E}_X\left[p(z,\sigma)\frac{d^2(z)}{\sigma^2}\right] &\propto \frac{1}{\gamma^d\sigma^2} \underbrace{\int \exp\left(-\|z-X\|^2/\sigma^2\right) d^2(z)\,\mathrm{d}z}_{\text{total variance of (unnormalized) } \mathcal{N}(0,(1/2)\sigma^2 I)} \\
&\propto \frac{\sigma^d}{2^{d/2}\gamma^d\sigma^2}\frac{d}{2}\sigma^2 && (17)\\
&\propto \frac{\sigma^d}{\gamma^d}, && (18)
\end{aligned}$$

where the factors of 2 and $d$ are absorbed into the proportionality constant. Similarly:

$$\begin{aligned}
\mathbb{E}_X\left[p(z,\sigma)\right] &\propto \frac{1}{\gamma^d} \underbrace{\int \exp\left(-\|z-X\|^2/\sigma^2\right) \mathrm{d}z}_{\text{normalization factor of } \mathcal{N}(0,(1/2)\sigma^2 I)} \\
&\propto \frac{\sigma^d}{2^{d/2}\gamma^d} \\
&\propto \frac{\sigma^d}{\gamma^d}.
\end{aligned}$$

Repeating the above calculations for $Y$ shows that:

$$\mathbb{E}_Y\left[p(y,\sigma)\frac{d^2(y)}{\sigma^2}\right] \propto \mathbb{E}_Y\left[p(x,\sigma)\right] \propto \frac{\sigma^d}{\alpha^d\gamma^d}.$$

Choosing $\sigma_X^2$ to achieve target perplexity for $X$, we see that setting $\sigma_Y^2 = \alpha\sigma_X^2$ makes the above terms equal to their corresponding terms in $X$ and thus achieves the target perplexity. □

#### 3.2 Scaling of the local radius with variance and length-scale

We now turn to the definition of local radius. Recall from Equation (8) in Methods that our local radius is defined as:

$$R_{P_X} = Z_X^{-1} \sum_j d^2(X_j) p(X_j, \sigma_X).$$

So, assuming the samples are drawn from generating distribution $F_X$, when we approximate this in the large-sample limit by $T_X(\sigma^2)$, as we did for the length-scale, we have:

$$T_X(\sigma^2) = \frac{\mathbb{E}_{z\sim F_X}[d^2(z)p(z,\sigma)]}{\mathbb{E}_{z\sim F_X}[p(z,\sigma)]}. \tag{19}$$

For the Gaussian and uniform generating distributions discussed in Propositions 3.1.1 and 3.1.4, it is straightforward to show that the local radius scales with the variance of the underlying distribution.

**Proposition 3.2.1.** *Let $F_X$ and $F_Y$ be a Gaussian or a spherical uniform distribution centered at $X$ and $Y$ respectively. For the Gaussian case, assume $X$ has a covariance matrix $\Sigma$, and for the uniform distribution assume a radius $\gamma$; $Y$ has covariance $\alpha\Sigma$ or radius $\sqrt{\alpha}\gamma$. Then, given $\sigma_X^2$, the length-scale for $X$, and $\sigma_Y^2 = \alpha\sigma_X^2$ as in Propositions 3.1.1 and 3.1.4, $T_Y(\sigma_Y^2) = \alpha T_X(\sigma_X^2)$.*

*Proof.* Note that for a given distribution,

$$T_X(\sigma_X^2) = \sigma_X^2 \mathcal{G}_X(\sigma_X^2) - \log \mathbb{E}_X[p(z,\sigma_X)].$$

If $F$ is Gaussian, then plugging the value of $\mathcal{G}_X(\sigma_X^2)$ from (13):

$$T_X(\sigma_X^2) \propto \sigma_X^2 \operatorname{Tr}\left[\Sigma(2\sigma_X^2 I + \Sigma)^{-1}\right],$$

and if $F$ is uniform, then from (18):

$$T_X(\sigma_X^2) \propto \sigma_X^2 \frac{\sigma_X^d}{\gamma^d \sigma_X^2} \sigma_X^2 \propto \frac{\sigma_X^d}{\gamma^d} \sigma_X^2.$$

Now, plugging in $\sigma_Y^2 = \alpha\sigma_X^2$ into corresponding equations for $T_Y(\sigma_Y^2)$, we see, in the Gaussian case:

$$T_Y(\alpha\sigma_X^2) = C_1 \alpha\sigma_X^2 \operatorname{Tr}\left[\alpha\Sigma(\alpha\sigma^2 I + 2\alpha\Sigma)^{-1}\right] = \alpha T_X(\sigma_X^2),$$

where the proportionality constant is the same for $T_X$ and $T_Y$ (as it does not depend on the covariance). In the uniform case:

$$T_Y(\alpha\sigma_X^2) = C_1 \left(\frac{\alpha\sigma_X}{\alpha\gamma}\right)^d \alpha\sigma_X^2 = \alpha T_X(\sigma_X^2),$$

as required. □

For more general distributions we are unable to show the explicit *linear* scaling of local radius with the variance of the underlying distribution as above. However, we can still connect the local radius to the length-scale parameter $\sigma$ chosen by t-SNE, which itself is known to empirically proxy the length scale at each point.

We show that the local radius $R_o$ in the original space is an increasing function of the length-scale parameter $\sigma$. Thus, the local radius recaptures the length-scale information lost when normalizing by $\sigma$. The proposition below builds off the connection made by Vladymyrov and Carreira-Perpiñán (2013) [3] between the t-SNE Gaussian kernel and the partition function from thermodynamics.

Here, we assume again that given a point $X$, its $n$-nearest neighbors are given by $X_1, \dots, X_n$, but unlike before, we assume *no* knowledge of the distribution of the $X_j$. Thus, we cannot use expectations of known distributions for large-sample approximations and instead consider a discrete distribution. To elucidate, define $\beta = \frac{1}{\sigma^2}$ and rewrite $R_X(\beta) := R_{P_X}(\sigma^2)$ from above suggestively as:

$$R_X(\beta) = \sum_{j=1}^{n} \frac{\tilde{q}(X_j, \beta)}{Z_X(\beta)} d^2(X_j),$$

where $\tilde{q}(z, \beta) = \exp\bigl(-\beta d^2(X_j)\bigr) = p(z, \sigma)$, and we make clear the sum $Z_X$'s dependence on $\beta$. Now, we note that since $Z_X(\beta) = \sum_j \tilde{q}(X_j, \beta)$, we can treat $q(X_j, \beta) := \tilde{q}(X_j, \beta) Z_X^{-1}(\beta)$ as probabilities from a *discrete* distribution, i.e. since $\sum_j q(X_j, \beta) = 1$. Using the moments of this discrete distribution, we can show that the local radius is a decreasing function of $\beta$ (and so an increasing function of the length-scale $\sigma$).

**Proposition 3.2.2.** *Let $R_X(\beta) = \sum_j q(X_j, \beta) d^2(X_j)$, where $\sigma = \beta^{-1/2}$ is chosen as before, to ensure constant entropy. Then $\frac{\partial R_X}{\partial \sigma} > 0$.*

*Proof.* Consider the expectation of $d^2$, taken over the distribution $q$:

$$\mathbb{E}_{X_j \sim q(\beta)}[d^2] = \sum_{j=1}^{n} q(X_j) d^2(X_j) = R_X(\beta). \tag{20}$$

Thus, $R_X$ is expectation of the variable $d^2$ over the discrete distribution $q$.

Now, we aim to understand the derivative of $R_X$ with respect to $\beta$. First it is straightforward to verify:

$$R_X(\beta) = Z_X^{-1}(\beta) \sum_j d^2(X_j) \exp\bigl(-\beta d^2(X_j)\bigr) = -\frac{1}{Z_X} \frac{\partial Z_X}{\partial \beta}.$$

Now, take the derivative:

$$\begin{aligned} R_X'(\beta) &= \frac{1}{Z_X^2} \left(\frac{\partial Z_X}{\partial \beta}\right)^2 - \frac{1}{Z_X} \frac{\partial^2 Z_X}{\partial \beta^2} \\ &= R_X^2 - \frac{1}{Z_X} \frac{\partial^2 Z_X}{\partial \beta^2}. \end{aligned}$$

By (20), we see that the first term is $\bigl(\mathbb{E}_{q(\beta)}[d^2]\bigr)^2$. Expanding the second term we see:

$$\begin{aligned} \frac{1}{Z_X} \frac{\partial^2 Z_X}{\partial \beta^2} &= \frac{1}{Z_X} \sum_j d^4(X_j) \exp\bigl(-\beta d^2(X_j)\bigr) \\ &= \mathbb{E}_{X_j \sim q(\beta)}[(d^2)^2], \end{aligned}$$

so this term is the second moment of the variable $d^2$ over the distribution $q$.

Thus, we have

$$R_X'(\beta) = \bigl(\mathbb{E}_{q(\beta)}[d^2]\bigr)^2 - \mathbb{E}_{q(\beta)}[(d^2)^2] = -\mathrm{Var}_q(d^2) < 0.$$

where $\mathrm{Var}_q(d^2)$ is the variance of $d^2$ calculated over $q$ and is therefore positive.

Since $R_X$ is a decreasing function of $\beta$, that means it is an increasing function of $\sigma$, since $\sigma = 1/\sqrt{\beta}$ is monotonic.

□
